# Efficient Multi-Kilobase Knockins in Mice and Cell Lines using CRISPR/Cas9 and rAAV Donors with Unbiased Whole-Genome characterization by LOCK-seq

**DOI:** 10.1101/2024.11.26.625457

**Authors:** Monica F. Sentmanat, Zi Teng Wang, Evguenia Kouranova, Samuel T. Peters, Wan Ching Chan, Jed Lin, Yong Miao, J. Michael White, Mia Wallace, Xiaoxia Cui

## Abstract

Multi-kilobase knock-ins (KIs) are a necessary, yet challenging type of genome editing to create and characterize in cell lines and animals. The combination of rAAV donor transduction and electroporation of single-cell mouse embryos with Cas9/gRNA ribonucleoprotein complex (RNP) enables highly efficient KI, but the insert size is limited by the viral packaging capacity. Here, we report the creation of up to 6.6 kb precise KI achieved in one step by using three rAAVs designed to insert one after the other. To fully characterize the edited genome with large KIs, we developed LOCK-seq (LOng-read sequencing of Captured Kilo-base targets), where relevant genomic regions are enriched via hybridization, achieving over 100-fold greater coverage compared to other long-read methods with enrichment. LOCK-seq simultaneously detects the presence of the KI allele, genotypes non-KI alleles and more importantly, uniquely identifies donor concatenation in the KI allele, and localizes random integration of the full or partial donor. Additionally, the multi-rAAV donor approach is successfully applied to cell lines, including lines intolerant of plasmid DNA, whereas LOCK-seq reliably and efficiently screens for KI clones. Together, the two approaches significantly improve the creation and precision of knock-in models.

**GRAPHICAL ABSTRACT:** 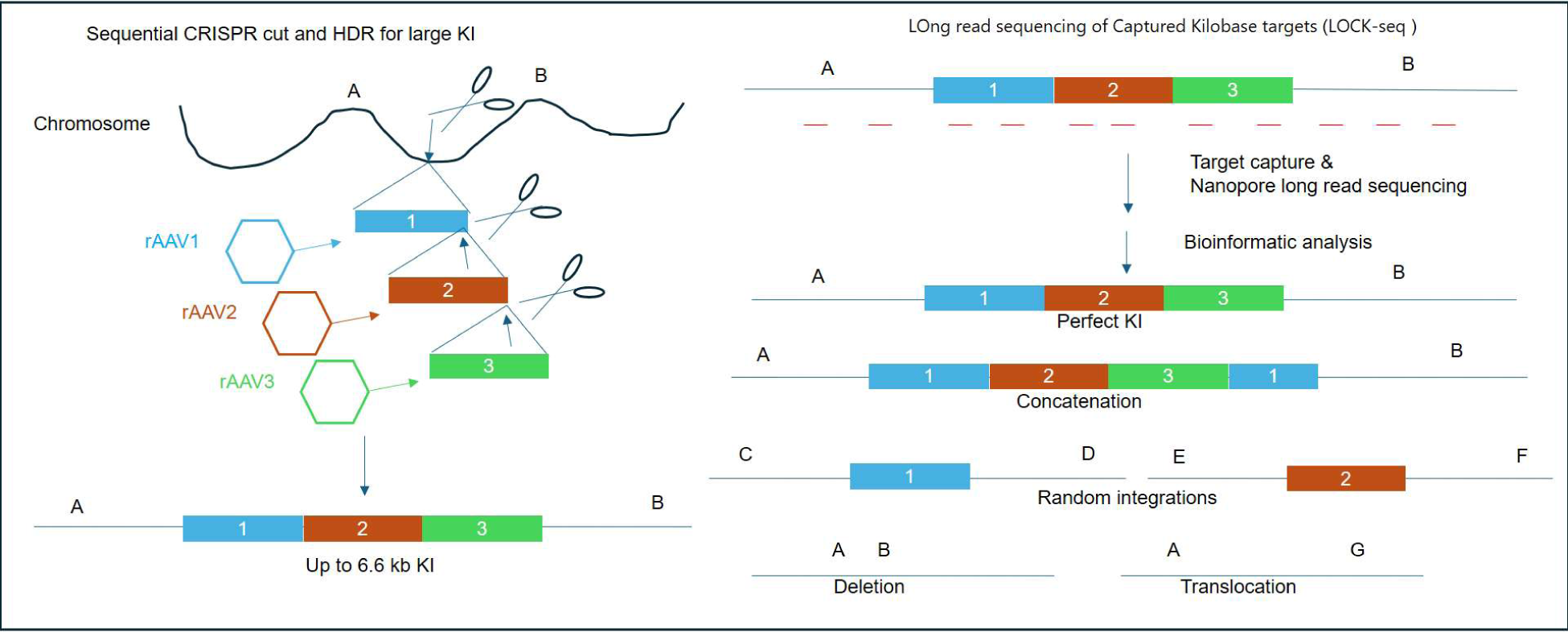

## INTRODUCTION

Genome engineering is indispensable for both emerging medical applications, such as gene and cell therapies, and the creation of research tools, including cell lines and model organisms. The continuous advancement of programmable nucleases, especially since the advent of CRISPR technologies in 2012(1), enables the introduction of increasingly sophisticated genomic modifications, allowing the recapitulation or correction of disease-relevant patient mutations, delivery of precise therapeutic cargos, or generation of a wide variety types of research models (e.g., floxed alleles, reporter lines, inducible expression) to facilitate mechanistic studies.

However, site-specific large insertions remain far more challenging than simpler modifications such as knockouts, deletions, and point mutations. Compared to editing cultured cells, editing single-cell mouse embryos is limited by the number of available embryos and by means of delivery. Electroporation of CRISPR RNPs and small single-stranded DNA donors into mouse embryos circumvents the need for microinjection and enables more embryos to be edited with remarkable efficiency for point mutations and small inserts, such as loxP site(2, 3). Plasmid DNA donors, however, must be microinjected into individual embryos, limiting throughput. The issue was partially addressed when rAAV donors were shown to sufficiently mediate HDR via simple incubation prior to electroporation of embryos with RNPs, bypassing the need for pronuclear microinjection(4–6).

rAAV as a donor format, however, has a maximum payload of 4.7 kb, restricting insert size. To create mice with larger KIs in a single step, we designed an approach to co-deliver multiple rAAV donors and Cas9/gRNA RNPs, building upon a previous report that achieved two-AAV mediated KIs in cells (7). Each donor carries a portion of the desired insert and mediates an HDR event upon proper cleavage by a respective RNP so that consecutive homology-directed repair (HDR) events reconstitute the full insert at the genomic locus. We demonstrate for the first time that co-delivery of up to three rAAV donors to mouse embryos reliably mediate multi-kilobase insertions. Specifically, we report the creation of 108 single rAAV, 13 two-rAAV, and 3 three-rAAV mouse models, achieving up to 6.7 kb precise insertions.

Successful large insertions are commonly identified by junction PCRs spanning from the insert into genomic sequences adjacent to homology arms. PCRs do not always work across the insertion junctions even when an integration occurs, depending on the locus and sequence context. This leads to substantial challenges with false negatives, as a failed PCR cannot be distinguished from a failed integration, and there are no obvious positive controls for these reactions. In addition, DNA templates in either plasmid or viral forms can randomly integrate into the genome, which may confound the intended use of the engineered research models(8–11).

While high fidelity, long-read, whole genome sequencing with deep coverage is ideal for thorough characterization of the edited genome, it is currently too costly and low-throughput as a screening tool. Targeted alternatives have been described, namely Cas9-targeted sequencing (nCATS) with or without long DNA size-selection steps (12, 13), but the on-target coverage and throughput are too low for screening multiple founders or clones.

To obtain more efficient target enrichment and enable thorough characterization of on-target large KIs and simultaneously counter-screen for undesired edits (imperfect KIs, indels, donor random integrations), we developed LOCK-seq (LOng read sequencing of Captured Kilo-base targets) on the affordable and accessible Oxford Nanopore MinION platform. Fragmented and barcoded genomic DNA samples are pooled for hybridization with biotinylated probes tiling the insert, homology arms and flanking genomic regions. Up to 100 samples for the same target or mixed targets can be enriched in the same hybridization reaction and sequenced in a single MinION flow cell. In addition to validating on-target precise KI events, LOCK-seq readily detects imperfect or partial insertions, donor concatenation and random integration, all of which can undermine model precision. As importantly, the per sample cost of LOCK-seq is only a fraction of that of nCATS. LOCK-seq is a practical method for fast and cost-effective screening of large numbers of samples.

Both methodologies are also applicable to editing stem and cancer cell lines. Some cell lines are highly sensitive to naked exogenous DNA and do not survive transfections with plasmid DNA. This sensitivity makes it difficult to introduce large insertions in these cell lines by using plasmid donors or linear templates with end modifications designed to enhance delivery efficiency(14, 15). Using rAAV donors circumvents the lethality from DNA transfection in cell lines, such as BV2. LOCK-seq, on the other hand, allows quick and cost-effective screen for correctly targeted clones, regardless of DNA donor format.

In summary, we report two complementary methods to significantly improve the precision of mouse and cell line models with large KIs: co-delivery of multiple rAAV donors and CRISPR/Cas9 RNPs to enable sequential HDR events and achieve multi-kilobase insertions and LOCK-seq for a much-needed, thorough characterization of the edited genome.

## MATERIAL AND METHODS

### gRNA design

gRNAs were designed using an in-house algorithm that incorporates off-target MIT specificity scores developed by the Zhang lab (16). The selection of gRNAs prioritized higher specificity of spacer sequences with low off-target scores and proximity of the cut site to the insertion site. Synthetic gRNAs were purchased as Alt-R single gRNAs from IDT (Coralville, Iowa). All gRNAs used for projects in this study are listed in **Table S1**.

### rAAV donor design and viral production

Donors were designed to have 400 to 1200 bp homology arms, with the majority in 700-800 bp range. It is critical to ensure the gRNA target site is not present in the donor. rAAV donor plasmids were synthesized by VectorBuilder (Chicago, IL), BioBasic (Amherst, NY), or Genscript (Piscataway, NJ) with flanking canonical AAV2 ITR sequences or cloned into a vector carrying canonical ITRs (5’cctgcaggcagctgcgcgctcgctcgctcactgaggccgcccgggcaaagcccgggcgtcgggcgacctttggtcgcccggcctcagtgagcgagcgagcgcgcagagagggagtggccaactccatcactaggggttcct-3’) or one canonical and one alternative ITR (5’-aggaacccctagtgatggagttggccactccctctctgcgcgctcgctcgctcactgaggccgggcgaccaaaggtcgcccgacgcccgggcggcctcagtgagcgagcgagcgcgcagctgcctgcagg-3’). All donors used for projects in the study are listed in **Table S1**. Production of rAAV6 virus was performed by the Hope Viral Vectors Core (Washington University, St. Louis, MO) using iodixanol gradient purification, or by VectorBuilder (Chicago, IL) or Genscript (Piscataway, NJ) as a crude preparation concentrated from supernatant. We did not observe a difference in efficiency between these two types of preparations, both using qPCR of ITRs for titer determination.

### Validation of sgRNAs and rAAV donors in N2a cells

N2a cells were cultured in DMEM media +10% FBS, +1% Glutamax, +1% Pen-Strep at 37°C in a humidified incubator with 5% CO_2_ and passaged using 0.25% Trypsin-EDTA using standard cell culture techniques. Cas9/gRNA RNPs were complexed at room temperature by mixing 1 µl of recombinant Cas9 protein (40 µM, from QB3 MacroLab, UC Berkeley) with 1 µl of each sgRNA (100 µM, resuspended in IDT Duplex Buffer, cat.#11-051-12). Nucleofections were performed using 150,000 cells in 20 µl OPTI-MEM or P3 solution and DS-137 program. After nucleofection, cells were immediately seeded in a well of a 24-well plate with 500 µl of growth medium supplemented with 1 µl of each rAAV donor and harvested 72 hr later for genotyping. All reagents for mouse projects were validated in N2a cells first before being used in mouse production.

### Amplicon NGS for CRISPR cleavage activity at the target site

Transfected N2a cells or tail clips of mice were lysed in QuickExtract Solution (Biosearch Technologies, cat.#QE09050), following manufacturer’s instructions. The target regions were PCR amplified by tailed primers appended with 5′-CACTCTTTCCCTACACGACGCTCTTCCGATCT-3′ for forward and 5′-GTGACTGGAGTTCAGACGTGTGCTCTTCCGATCT-3′ for reverse to genomic-specific primer sequences (PCR 1), which allows unique indexes and Illumina P5/P7 adapter sequences to be added in a second round of PCR. All primer sequences are listed in **Table S2**. PCR amplifications were performed with JumpStart REDTaq ReadyMix (MilliporeSigma, cat.#P0982), following the manufacturer protocol. The following cycling conditions were used: 9°C for 2 min, followed by five cycles of 94°C for 30 s, 54°C for 30 s, and 72°C for 40 s. We generated 2 × 250 reads with the Illumina MiSeq platform at the Center for Genome Sciences and Systems Biology (Washington University, St. Louis, MO). The extracted FASTQ files are analyzed using a Python-based script that outputs editing efficiency(17).

### Junction PCRs to confirm targeted integration

The same N2a lysates above were used as templates in junction PCRs for detecting knock-in events. Each pair of junction PCR primers contain one that anneals to the flanking genomic region and the other, to the insert. PCR amplification was performed using primers listed in **Table S2** with Platinum SuperFi II Green PCR Master Mix (ThermoFisher, cat.#12369050) or JumpStart REDTaq ReadyMix (MilliporeSigma, cat.#P0982).

### Mouse husbandry and embryo manipulation

All animals at Washington University in St. Louis are housed under SPF barrier conditions in AALAC-accredited facilities. All required breeding, experiments and interventions are included in IACUC approved protocols. The Department of Comparative Medicine provides basic husbandry in accordance with their procedures. Three to four weeks-old C57BL/6J mice (JAX Laboratories, Bar Harbor ME, USA) were superovulated by intraperitoneal injection of 5 IU pregnant mare serum gonadotropin, followed 47h later by intraperitoneal injection of 5 IU human chorionic gonadotropin (PMS from SIGMA, HGC from Millipore USA). Mouse zygotes were obtained by mating C57BL/6J stud males with superovulated C57BL/6J females at a 1:1 ratio.

rAAV transduction of zygotes was performed as previously reported(4, 5). Briefly, 10^9^-10^10^ viral particles were added to each KSOM droplet containing 20-30 zygotes covered with paraffin oil and allowed to incubate for 5-6 hours in a humidified tissue culture incubator (37°C, 5% CO_2_). The zygotes were then washed and electroporated with Cas9/sgRNA RNP complexes as previously described(3). Briefly, one-cell fertilized embryos were electroporated with RNPs containing 12 μg of Cas9 protein complexed with 3.6 μg of gRNA (1:1.5 molar ratio). Between 30 to 40 post-transduction zygotes in 10 µl of OPTI-MEM are mixed with 10 µl of RNP in OPTI-MEM before being loaded into a 1 mm electroporation cuvette. With a Bio-Rad Gene Pulser Xcell electroporator, we use 2-6 pulses of 30V for 3 ms with 100 ms internals. Post-electroporation embryos were kept in 25 μl drops of KSOM overlayed with oil and in a humidified tissue culture incubator (37°C, 5% CO_2_) for 1-2 hours before being surgically transferred to 0.5d PC pseudopregnant surrogates.

For projects using multiple rAAVs, all rAAV donors were combined at equal number of viral particles before adding onto the zygotes. After incubation, the washed zygotes were electroporated with all necessary RNPs at equal molar ratio, with the total RNP mass per electroporation held constant at 15.6 μg.

### Founder genotyping by junction PCRs and detection of random integration by ITR PCRs

Founder genotyping was done using junction PCR conditions optimized during validation. To detect ITR-containing donor random integration, a primer that binds to the hairpin of the ITR, 5’-TGGCCAACTCCATCACTAGG-3’ (**Table S2**, XCC216c.AAV2.ITR.F2-R2), is paired with an insert specific primer at each end of the donor.

### Cas9-targeted sequencing (nCATS)

Genomic DNA from tail clips was isolated with the Monarch Genomic DNA Purification Kit (NEB, cat.#T3010S). nCATS was performed using the Oxford Nanopore’s Cas9 Sequencing Kit (cat. #SQK-CS9109) according to the manufacturer’s instructions. Briefly, 5 μg of extracted DNA was 5’ dephosphorylated and cut with a pair of RNPs for the ROSA locus (Project XCC113, **Table S1**) (5’-TAAGGGAGCTGCAGTGGAGT*NGG*-3’, 5’ GGATTTAGCCACATCCATAG*NGG*-3’) flanking the region of interest. RNPs were each prepared using 100 μM of gRNA complexed with 30 μM of Cas9 protein (QB3 MacroLab, UC Berkeley) and added in a dA-tailing cocktail at 37°C for 35 min. Adapter ligation and clean-up was performed using the Ligation Sequencing Kit V14 (Oxford Nanopore Technologies, cat.#SQK-LSK114) and the final library run on an R10.4.1 MinION flow cell (Oxford Nanopore Technologies, cat.#R10.4.1).

### LOCK-seq (LOng-read sequencing of Captured Kilo-base targets)

Genomic DNA was isolated from tissue and cell pellets using the Monarch Genomic DNA Purification Kit (NEB, cat.#T3010S) or with QuickExtract DNA Extraction Solution (Biosearch Technologies, cat.#QE0901L), according to the respective manufacturer’s instructions.

#### Tn5 tagmentation*Tagmentation*

Tagmentation with Tn5 LongPlex Long Fragment Multiplexing Kit (seqWell, cat.#301312) was used according to the manufacturer’s protocol or an in-house protocol for multiplexing. For in-house tagmentation, Tn5 protein (QB3 MacroLab, UC Berkeley) was complexed with annealed oligonucleotide adapters as described by Picelli *et al.*, 2014(18). Briefly, 16.5 μM of adapter primers (Tn5ME-A 5’-TCGTCGGCAGCGTCAGATGTGTATAAGAGACAG-3’ and Tn5ME-B 5’-GTCTCGTGGGCTCGGAGATGTGTATAAGAGACAG-3’, each annealed with Tn5MErev 5’-[phos]CTGTCTCTTATACACATCT-3’) was added to Tn5 protein (500 ng/μl) and incubated at room temperature, shaking at 200 rpm. Tagmentation was performed with 10 μl of crude lysate in 10 mM TAPS-NaOH, pH 8.5, 5 mM MgCl2, and 7% PEG 8000 in a total reaction volume of 20 μl for 5 min at 55°C. The resulting fragmented genomic DNA was cleaned up using 0.7x JetSeq Clean magnetic beads (Meridian Biosciences, cat.#BIO-68032) and eluted in 20 μl.

Barcodes for dual indexing were introduced by PCR using KOD Xtreme Hot Start DNA Polymerase (MilliporeSigma, cat.#71975-3), with primers specific to the universal adapter tail sequences introduced by Tn5. Each PCR reaction used the full volume of fragmented DNA from the previous step with 2 mM of each dNTP and 10 μM of each primer in a final volume of 50 μl. Cycling conditions were 94°C for 2 min, 8 cycles of 98°C for 10 sec, 68°C for 10 min, and a final 72°C hold for 10 min.

#### Mechanical fragmentation

Up to 5 μg of genomic DNA prepared with the Monarch Genomic DNA Purification Kit (NEB, cat.#T3010S) was loaded onto a Covaris g-TUBE (Covaris, cat.#520079) and centrifuged at 4900 x g for 1 min to generate fragments in the 10 kb size range. Fragmented DNA was end-repaired and A-tailed using the TWIST Mechanical Fragmentation Kit (TWIST, cat.#101281), ligated to TruSeq compatible adapters (TWIST, cat.#101310) and dual-index barcoded using PCR with KOD Xtreme Hot Start Polymerase, following the manufacturer’s protocols. Specifically, the cycling conditions include 94°C for 2 min, 8 cycles of 98°C for 10 sec, 58.8°C for 30 sec and 68°C for 10 min, and a final 68°C hold for 10 min. Indexed samples were pooled to a final mass of 4-6 μg for hybridization.

#### Biotinylated probes

Custom capture probes were acquired as 120-mer, end-labeled xGen oligos from IDT or generated in-house using a two-step PCR and tiled at densities of 1-4 probes/kb (commercially purchased and home-made probe sequences listed in **Table S3**).

In-house generated LOCK-seq probes presented in **Fig.6** used tailed primers appended with 5’ ggcatttcagtcaggtgcccaatgtacc-3’ for forward and 5’-gttccgggtaggcagttcgctccaagct-3’ for reverse to genomic-specific sequences in PCR 1, followed by an asymmetric PCR 2 using universal primers, 5’biotin-agtcaggtgcccaatgtacc-3’ and 5’biotin-taggcagttccgctccaagc-3’. Primer sequences for all in-house generated probes with tails used are listed in **Table S4**. PCR amplifications were performed with JumpStart REDTaq ReadyMix (MilliporeSigma, cat.#P0982), according to the manufacturer’s protocol. The asymmetric PCR was performed using 0.1x volume of PCR product (no clean-up step) from PCR 1 with the forward primer at 10 μM and reverse primer at 0.5 μM and melting at 94°C for 2 min, followed by twenty cycles of 94°C for 30 s, 60°C for 10 s, and 72°C for 20 s. Amplicons were then pooled and cleaned-up using the Zymo DNA clean and Concentrator kit (Zymo Research, cat#.D4004). To generate multiple biotin moieties across probes during the asymmetric PCR step, KOD Xtreme Hot Start DNA Polymerase was used with 2 mM each dATP, dCTP, and dGTP, 1.6 mM dTTP and 0.4 mM Biotin-11-dUTP (ThermoFisher cat.#R0081), with the forward primer at 10 μM and reverse primer at 0.5 μM and melting at 98°C for 2 min, followed by 20 cycles of 98°C for 10 s, 60°C for 10 s, and 68°C for 20 s.

#### Hybridization

Multiplexed samples were hybridized with biotinylated probes using Twist Standard Hyb and Wash Kit (cat.#104446), and hybridization reactions were performed for 16-18 hr according to the manufacturer’s protocol. Briefly, the pooled DNA was incubated with Universal Blockers (TWIST, cat.#100578) and 5 μg of Mouse Cot-1 DNA (Invitrogen, cat.#18440016) at 95°C for 5 minutes, cooling at room temperature for 5 min to minimize non-specific capture in the hybridization reagent. The reactions were then heated to 70°C, and upon adding the probe panel, maintained at 70°C for 16-18 hr.

#### Post-hybridization enrichment

Target-captured libraries were enriched following the TWIST (catalog #104446) post-capture workflow. Briefly, biotinylated DNA–probe hybrids were captured with Dynabeads M-270 Streptavidin (ThermoFisher, cat.# 65305) at 68°C for 10 minutes. Beads were washed according to manufacturer’s instructions, and bound DNA was released with 0.2 N NaOH. The eluate was neutralized and used directly for post-capture PCR with KOD Xtreme Hot Start DNA Polymerase using TWIST universal adapter primers. Thermocycling conditions were 94°C for 2 min, 18 cycles of: 98°C for 10 sec; 58.8°C for 30 sec; 68 °C for 10 min, with a final extension of 68°C for 10 min. PCR products were purified by DINOMAG SPRI (Labscoop, cat.#DN9004-5ML) bead cleanup at 0.5× bead/sample ratio.

For **Fig3b**, **Fig.S9**, and **Fig.S12** (Project XCB528) a double hybridization was performed using the captured output from the first round as the input for a second round of hybridization, repeating the process with a fresh aliquot of probes. This can be performed to increase the percentage of on-target reads.

#### ONT library preparation

Ligation of Oxford Nanopore sequencing adapters was performed for all libraries using the Ligation Sequencing Kit V14 (Oxford Nanopore Technologies, cat.#SQK_LSK114) and loaded onto an Oxford Nanopore Technologies flow cell (FLO-MIN114 or FLO-PRO114HD).

#### LOCK-seq analysis

Base calling was performed using Guppy or Dorado (v0.7), and samples were demultiplexed with Nanoplexer or an in-house python script. The code for demultiplexing and the analysis pipeline is deposited at github.com/msentmanat/LOCK-seq-analysis-pipeline. Briefly, no trimming was performed, and for the in-house demultiplexing script, exact index matches for both indexes were required. Reads with >1 match for either index or lacking one index were excluded. Capture efficiency was determined using flagstat output (total reads mapped to transgene/total reads) and length of mapped reads calculated using FASTQ files derived from transgene mapped BAM files for each sample. FASTQ files were aligned to the transgene first using Minimap2 (v2.28), mapped reads (.fastq) were pulled using Samtools (v1.20), and then mapped to the reference genome (mm39 or hg38) using Minimap2 (v2.28) and Samtools (v1.20). Using the Samtools depth command, the top 10 genomic coordinates (based on highest read depth, ∼20 kb window) were used to generate a bed file to pull reads from the bam files. Those regions were investigated by manual curation on IGV and Canu to identify random integration events (**Fig.S1**). FASTQ files were used to build consensus sequences using Canu (v2.2) and visualized using Snapgene. The per base coverage was plotted with normalized reads from Samtools depth output in R.

### Gene editing in N2a, BV2, HEK293T, and iPSCs

N2a, BV2 and HEK293T cells were cultured in DMEM media +10% FBS, +1% Glutamax, +1% Pen-Strep at 37°C in a humidified incubator with 5% CO_2_ and passaged using 0.25% Trypsin-EDTA using standard culture conditions. iPSC cells were cultured on Matrigel-coated (Corning, cat.#354277) 6-well plates with mTeSR Plus (STEMCELL Technologies, cat.#100-0276) media at 37°C in a humidified incubator with 5% CO_2_, and passaged using ReLeSR (STEMCELL Technologies, cat.#100-0483) according to the manufacturer’s instructions. Cas9 RNPs were complexed at room temperature by mixing 1 µl of recombinant Cas9 protein (40 µM, from QB3 MacroLab, UC Berkeley) with 1 µl of each sgRNA (100 µM, resuspended in IDT Duplex Buffer, cat.#11-051-12).

Nucleofections were performed using 150,000-250,000 cells per 20 µl reaction. Cells were dissociated into single cells, washed twice with PBS, then resuspended into Opti-MEM media (N2a and HEK293T) or P3 solution (Lonza, cat.# V4SP-3096 for iPSCs) and nucleofected using a Lonza 4D nucleofector with standard programs (N2a: DS-137, HEK293T: CM-130, iPSCs: CA-137).

Nucleofected N2a, BV2 and HEK293T cells were seeded into 500 µl of complete culture medium in a well of a 24-well tissue culture plate, supplemented with 1 µL of each AAV donor, and harvested 72h post-nucleofection for genotyping and expansion for sorting. Nucleofection iPSC cells were seeded into 500 µl of mTeSR Plus media supplemented with 10% CloneR2 (STEMCELL Technologies, cat.#100-0691) for 24h, and the medium was changed to mTeSR Plus without CloneR2.

To generate clonal edited lines, pools were single-cell dissociated with 0.25% trypsin (N2a, BV2 and HEK293T cells) or 0.75x TrypLE (Fisher Scientific, cat.#12-604-013, iPSC cells), and single cells were sorted into 96-well tissue culture plates on a Sony SH800 cell sorter. N2a cells were sorted into complete media supplemented with 10 µM ROCK inhibitor (MilliporeSigma cat#SCM075) and 1 mM sodium pyruvate (ThermoFisher, cat#11360070) while HEK293T cells were sorted into 50% conditioned media, and iPSCs were sorted into mTeSR Plus supplemented with 10% CloneR2. Clones were monitored for growth, with changes of complete media every 3-5 days (N2A and HEK293T) or 2-3 days (iPSCs) until >40% confluent, then harvested, consolidated, and genotyped by junction PCRs as in described above.

## RESULTS

### rAAV donors mediate efficient HDR in mouse embryos

By transduction of mouse embryos with a single rAAV donor, followed by Cas9/gRNA RNP electroporation, we successfully created 108 mouse models with insert sizes ranging from 400 bp to 3.5 kb (**Fig.1a, Table S5**). Founder rates among live births were 1% to 80%, identified by junction PCRs (**Fig.1b**). There is a trend that the longer the insert, the less efficient the insertion. There is an inverse correlation between KI efficiency and length of insert/total homology. Each doubling of homology length relative to the insert results in a 30% or greater increase in efficiency of generating founders (R=-0.28, p-value =0.004; **Fig.1c**). Homology arms were designed between 400 and 1200 bp each, with a preference for longer arms – typically ∼ 800 bp – when compatible with AAV packaging limits. This design choice was informed by previous work using plasmid or AAV-based donors with programmable nucleases, including zinc finger nucleases and CRISPR/RNPs (7, 19, 20). These studies report robust HDR within this range, and comparative analysis of arm lengths for plasmid donors indicate that increased length enhances repair efficiency (21) but reducing them in rAAV donors to accommodate packaging limits may not significantly impact HDR (22). Fifteen out of the 108 models were gene replacements, generated using one donor and two gRNAs that excise the region targeted for replacement while facilitating HDR (**Fig.S2**). Gene replacement was as efficient as direct insertion among these models (**Fig.1d**).

**Figure 1.**
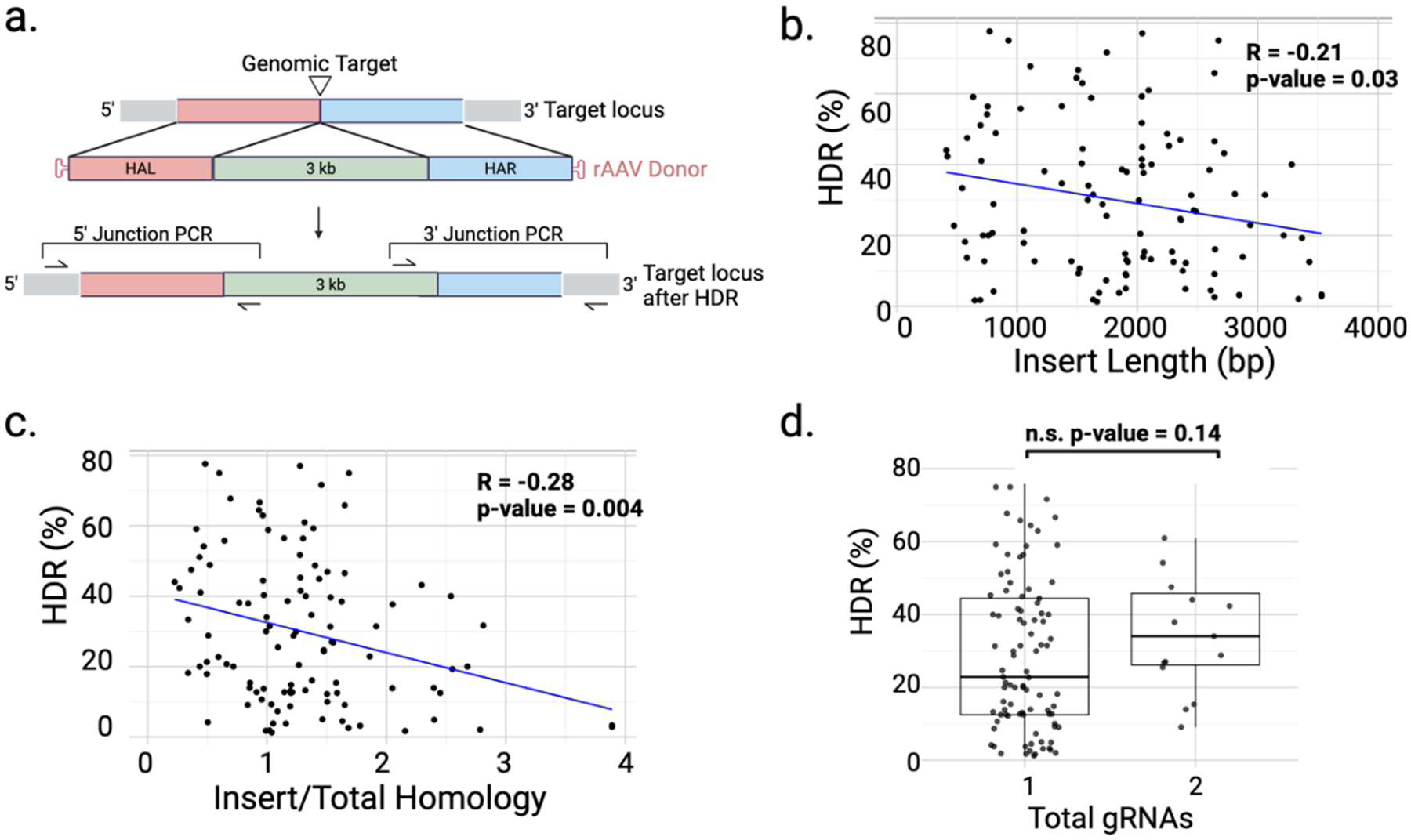
rAAV is an efficient donor format for editing mouse embryos. **a**. Schematic of rAAV donor and junction PCR designs. **b**. Scatter plot of KI efficiency by junction PCRs vs. insert length in 108 single rAAV KI models and **c**. KI efficiency vs. Insert length/total homology length. R represents Pearson’s correlation coefficient between x- and y-axis values shown. **d**. Boxplots comparing HDR efficiency of direct insertion using a single gRNA vs. gene replacement KI models using two gRNAs. n.s. represents not significant with a p-value cut-off of >0.05 for significance.

### Sequential insertion of multiple rAAV donors for larger inserts

To overcome the 4.7 kb size limitation of an rAAV genome, we designed a sequential insertion strategy, where the first donor delivers a portion of the insert along with a gRNA target site, upon cleavage, to mediate the subsequent insertion from the second donor (**Fig.2a**). The gRNA target site in the donor can be identical to the genomic target site, or unrelated, such as a previously validated site from a different species or a new site created by the insertion. The gRNA target is not cleavable in the single-stranded rAAV genome. For even larger inserts, the second rAAV donor brings in an additional gRNA target site to mediate the third insertion (**Fig.2a**). All rAAV donors and RNPs are delivered to single-cell embryos in a single step. The consecutive HDR events happen one after the other upon respective CRISPR cleavage within the embryos. To date, we have completed 13 models with two rAAV donors and 3 models with three rAAV donors.

**Figure 2.**
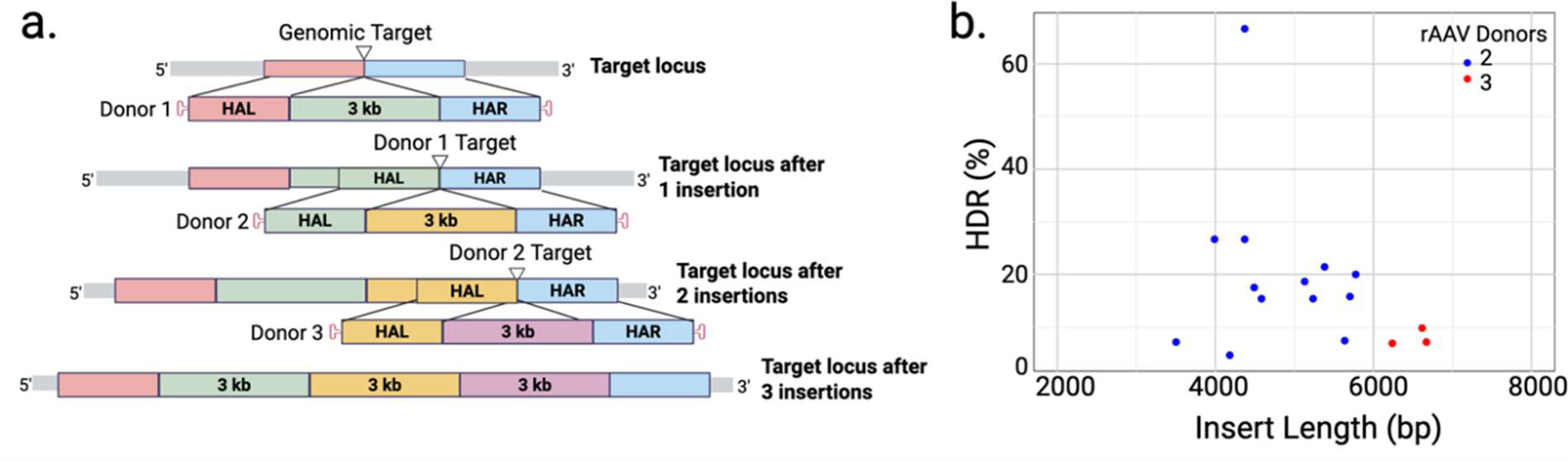
Co-delivery of multiple rAAV donors greatly increases the size of KI insert achieved in one-step editing. **a**. The schematic of sequential insertions using two or three co-delivered rAAV donors. **b**. Scatter plot of KI efficiency by junction PCRs vs. insert length in 16 models with two-(blue dots) and three-rAAVs (red dots).

### Detectable recombination between rAAV donors

The increasing insert size via sequential HDR events renders genotyping by junction PCRs tedious and inconclusive for several reasons, including increased number of junctions, difficult sequences, mosaicism for the KI allele in founders, and partially correct inserts. The use of multiple rAAV donors further complicates screening, as each additional donor introduces two more new junctions. Multiple PCRs are required to screen founders, targeting both the insert and genomic junctions as well as junctions between donors. Results can be difficult to interpret when animals are positive for only a subset of PCRs (**Fig.S4)**, because negative PCR reactions can be the result of lack of junctions or tricky sequences to amplify. Adding to the complexity, founder animals are often mosaic and harbor more than two alleles. On the other hand, the presence of positive junctions does not guarantee a full-length insertion event (**Fig.S5**). And the correctly targeted allele, if much less abundant, can be missed in PCR detection. A reliable method with minimal bias is required for screening such complex events with confidence. Additionally, rAAV random integration has been reported previously(23, 24). To better prioritize animals without viral genome integration, we designed primers that bind to the hairpin region of the ITRs and donor sequence so that animals negative for the ITR-specific PCR and positive for junctions are prioritized (**Fig.S6, Table S5**). On average, 37% of junction positive animals were also positive for ITRs. However, the PCR-based method does not reveal the random integration sites in the genome and will miss random integrations without ITRs.

### LOCK-seq

To better characterize rAAV-mediated KI events, we first used the Cas9 Sequencing Kit (a.k.a. nCATS) from Oxford Nanopore (SQK-CS9109), which relies on Cas9/gRNA binding and cleavage close to the target. Although on-target reads were captured, the low coverage and throughput and high cost make it impractical for screening multiple founders (**Fig.S7**). A new method with significantly higher efficiency of target enrichment than using Cas9/CRISPR recognition is needed.

We developed a target-capture approach using 5′-biotinylated 120-mer DNA oligos complementary to the insert, homology arms, flanking genomic regions, and ITRs, which we term LOng read sequencing of Captured Kilo-base targets (LOCK-seq). High molecular weight genomic DNA is fragmented (by using Tn5 transposition or physical sharing with Covaris g-TUBEs), ligated to adapters with barcodes, and undergoes low-cycle PCR amplification to enrich adapter-ligated products. Barcoded samples are pooled and hybridized to biotinylated probes for selective pull-down of sequences of interest. Then a second, low-cycle PCR is done to further enrich fragments ligated with adaptor sequences before motor protein is ligated and sequencing is run.

In a direct side-by-side comparison, LOCK-seq yielded >100-fold on-target coverage than nCATS (**Fig. S7**). This is attributable to two main factors. First, higher on-target capture efficiency by LOCK-seq (10% or greater), whereas for nCATS is about 1%(12, 25, 26). Second, there is excess adapter-free DNA in nCATS library, which we hypothesize can transiently interact with pores and motor-protein bound DNA, reducing pore occupancy, without occupying the pores productively. The more efficient target enrichment and in turn increased read depth by LOCK-seq enables multiplexing of many barcoded samples to run on a single MinION flow cell, making large-scale genotyping of founders feasible for both throughput and cost considerations (**Fig. 3a**). Conveniently, as little as 250 ng of genomic DNA per sample is required for LOCK-seq, compared to 1-5 µg for nCATS, which is a significant amount to isolate from routine tissue biopsies of founder animals. Critically, the probes are designed against the insert, homology arms as well as genomic regions flanking homology arms, allowing LOCK-seq to identify both the knock-in allele and alleles containing indels at the target site in a mosaic animal (**Fig.S8**), as well as random integration of the full or partial donor(s) and the location of each random integration in the genome.

**Figure 3.**
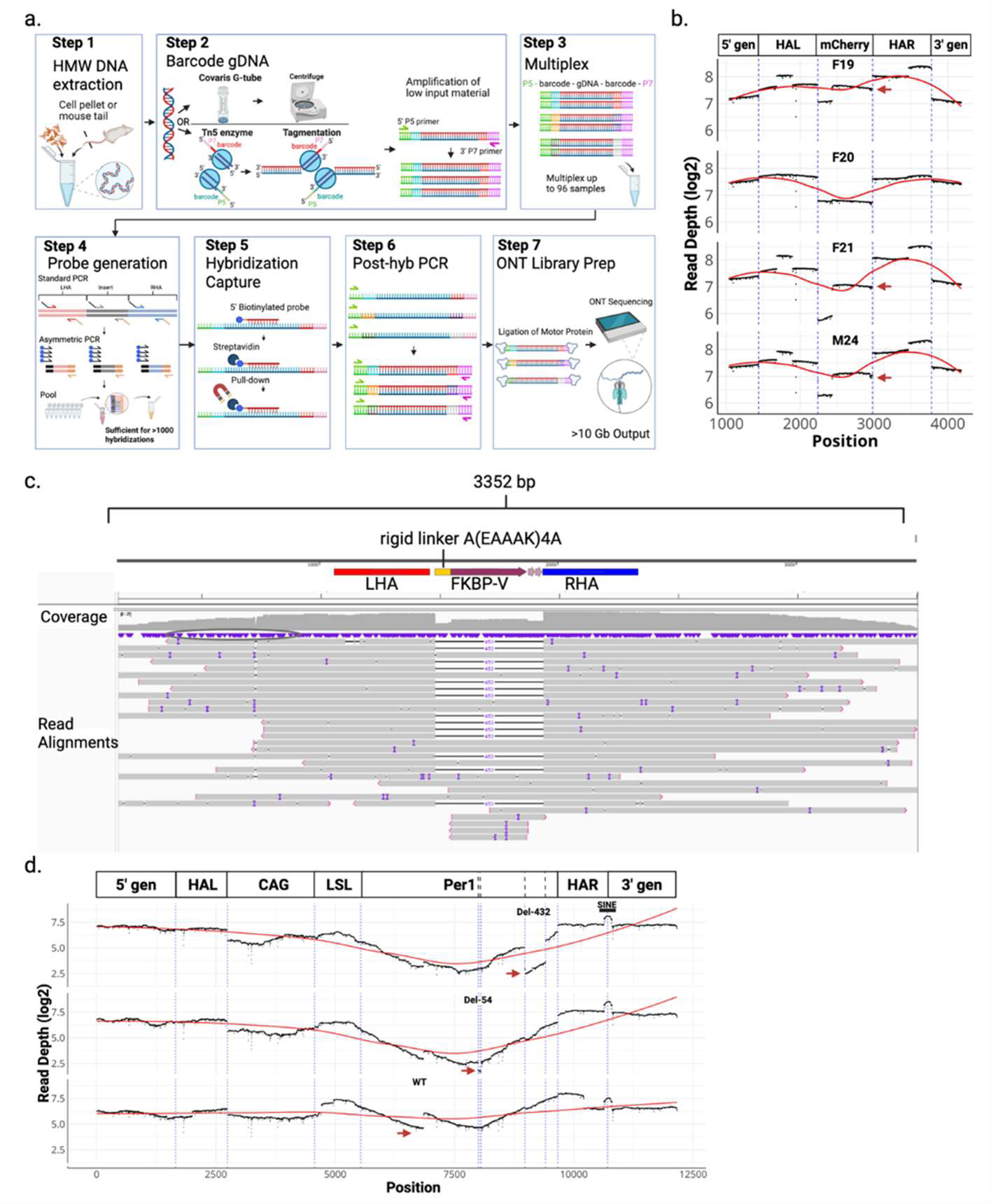
LOCK-seq for detection of on-target KI events. **a**. Schematic of LOCK-seq. **b**. Dotplot of read depth (log2 normalized) per base across the XCB528 mCherry insertion (schematic on top). Dashed vertical lines (blue) mark boundaries of insert and homology arms. Red arrows point to copy number variation. **c**. IGV screenshot of aligned bam file for Project MS2757 illustrating multiple individual raw reads (grey bars) that cover the length of the cassette and extend into flanking genomic regions for both KI and wild-type alleles. The black horizonal bar across read indicates absence of FKBP-V insert, illustrating the heterozygous state of the F1 animal sequenced. Raw, uncorrected reads are shown, vertical purple bars within reads indicate sequencing error. **d**. Dotplot of read depth (log2 normalized) per base across the longer insert mediated by XCC113 3-rAAV donors. Dashed vertical lines (blue) mark boundaries of genetic regions: gen5, 5’ genomic; L and R, left and right homology arms, CAG, CAG promoter; LSL: floxed polyA signals; Per1, cDNA variants (hashed boxes); pA, polyA signal. Red arrows mark copy number variation.

We first analyzed four mouse samples (a founder, M24, and three F1s of M24: F19, F20 and F21) with a 741 bp mCherry cassette inserted at the Syk gene, one of the 108 mouse models created using a single rAAV (**Table S5**). The genomic DNA was fragmented using Covaris g-TUBEs. Biotinylated 120-mer single stranded oligo probes tiled the entire length of the 2.4 kb rAAV genome and 2 kb of flanking sequences. The samples were pooled and underwent a double hybridization reaction where the enriched products from the first found of hybridization underwent a second hybridization with fresh probes to maximize specificity. The post-hybridization, single-stranded DNA was then PCR amplified, ligated to motor protein and run on a PromethION flow cell (8x the number of pores compared to MinION flow cell), given previous poor depth obtained using nCATS.

The average length for mapped reads was 2.7 kb across these four samples, with the maximum read-lengths ranging from 84-102 kb, average on-target coverage of 254K reads/sample and resulting in a capture efficiency of 85% (**Table S7**). Full-length, continuous reads spanning the donor and flanking genomic DNA outside of both homology arms were recovered for all samples (**Fig.S9**). Out of the four samples, F20 was closest to a heterozygous insertion with equivalent coverage across the insert, supporting the absence of aberrant integration events (**Fig.3b**). The other two F1 samples, F19 and F21, shared similar patterns in coverage variation as the founder, indicative of random integration events that were further analyzed below. Our initial workflow, which paired a PromethION flow cell with a double-hybridization protocol, provided substantially higher sequencing depth than was necessary for multiplexed samples. Accordingly, all later samples were processed using MinION flow cells and a single-hybridization workflow.

**Fig.3c** shows an IGV screenshot of raw aligned reads (bam files) used to generate coverage plots and identify deviations from expected read depth consistent with a single copy number of an FKBP-V insertion (**Table S5**, Project MS2757). The data identifies the founder has both a knock-in and wild-type allele, demonstrating that LOCK-seq can resolve zygosity (detailed description of IGV alignments **Fig.S10-S11**).

We then analyzed three related models generated by co-delivery of three rAAV donors for conditional expression of a full-length Per1 cDNA or the same Per1 cDNA with 54 bp and 432 bp deletions, respectively. For each model, only the third donor is unique. The longest insert of the series is 6.7 kb, and the founder rate was 2-7% HDR (**Table S6**, Project XCC113). With this set of samples, we tested an alternative fragmentation method using Tn5, called tagmentation, to simultaneously fragmentate and barcode the DNA samples for convenient multiplexing. We obtained an average read length of 1 kb across samples, with maximum read lengths between 10-14 kb. The capture efficiency was 11-19%, with an on-target sequencing depth range across samples of 6K to 38K reads (**Table S7**). The 54 bp and 432 bp deletions are evident in respective samples (**Fig.3d**). An increase in coverage for a portion of the wild type Per1 cassette implies concatenation of donor 3, which was further analyzed below. Compared to Covaris g-TUBEs, enzymatic tagmentation is convenient and high throughput, with the downside of shorter average fragment size, hence short read lengths. However, it is a useful method for quickly screening tens of samples.

### Various undesired modifications detected only by LOCK-seq

We next looked deeper at undesired edits among the samples at or away from the target sites. The per-base read depth for M24, F19 and F21 from project XCB528 targeting mCherry to the 3’ of the endogenous mouse Syk locus suggests partial integration of the 3’ of mCherry and homology arms (**Fig.3b**). Alignments of mapped reads to the mm39 mouse genome revealed a random integration event within the first intron of Cd63 (chr10: 128,743,596, mm39) and 5 kb upstream of neurotransmitter transporter Slc6a15 (chr10: 103,198,832) in M24 (mosaic founder) that were transmitted to F19 and F21, but not F20 (**Fig.4a**). Both random integrations had the viral ITR present and were confirmed by site-specific PCRs of the insertion junctions (**Fig.S12**). A third random integration at a transcript with unknown function, 49305000H12Rik (chr16:73,129,317), was detected in Founder M24 but not in these three F1s analyzed.

**Figure 4.**
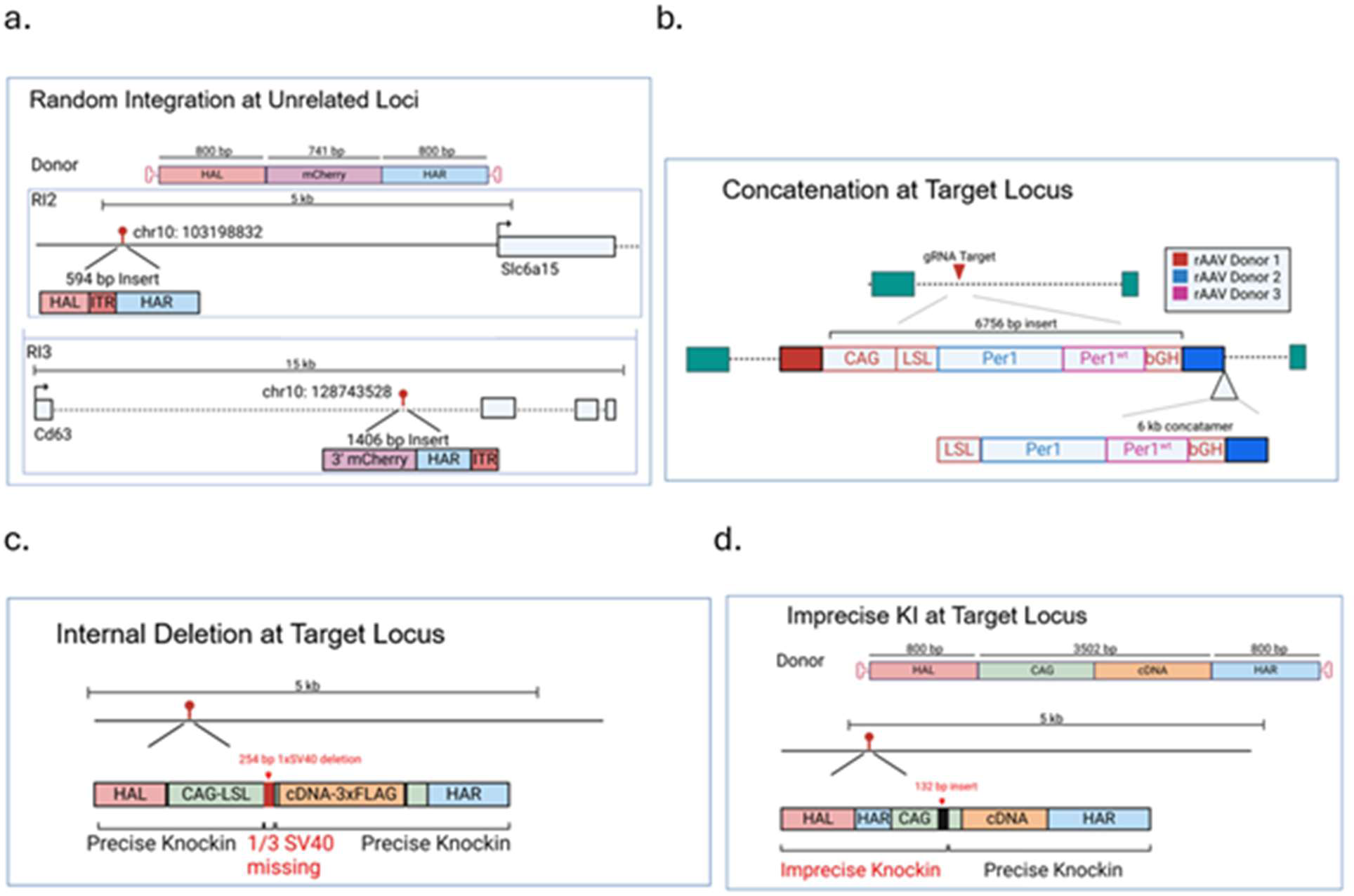
Schematics of various unintended editing outcomes in mice: **a**. random integration at unrelated, non-target loci, **b**. deletion of repetitive element, **c**. donor concatenation at the target site, and **d**. imprecise KI resulting in additional sequence at 5’ end of transgene.

The variation in coverage across the cDNA cassette in the WT Per1 cDNA model proved to be a concatenated insertion event at the target site that was evident after de novo assembly of reads, despite the shorter average read length using Tn5 tagmentation (**Fig.4b**). The concatenation junctions were confirmed to be present only in the WT model by PCR, not in the two other models with shorter cDNAs (**Fig.S13**). Besides concatenation, we also observed frequent internal deletions of one of the three SV40 polyA signal across multiple projects. An example is shown for the conditional CAG promoter-driven construct (CAG-loxP-3xSV40 termination signal-loxP) used for the 1 kb Ldhb-3xFLAG mouse cDNA KI (Project XCD722a;**Fig.4c**), which would likely be missed by junction PCRs. Additionally, we observed one-sided imprecise KIs. For example, in one founder of the 3.5 kb pH-sensitive fluorescent reporter Mito-SypHer, which includes a mitochondrial targeting sequence and was inserted at the at the ROSA26 locus (Project MS2750; **Table S5**), we identified the insertion of an additional right homology arm along with a short extraneous sequence of the CAG promoter (**Fig.4d**).

We also explored the possibility of technical artifacts, such as fusion reads that could be mistaken as indicate random integration or concatenation of the donor(s). Such artifacts have previously been reported to occur in ∼2% of ONT data generated from amplicon sequencing (27). We ran two MinION flow cells, each combining two LOCK-seq libraries prepared separately from different KI projects with unique inserts (**Table S8**) and detected less than 2% of reads to be fusion or split reads, i.e. single reads aligning to both inserts belonging to the two different libraries (**Fig.S14**).

### Flexible and cost-effective probe design and generation

The biotinylated 120-mer oligo probes are costly, especially when used to tile across the genomic region with a large insert. To help reduce cost, we designed amplicons spaced ∼300 bp apart along the genomic region and primers with universal tails to create probe templates for a second, asymmetric PCR with biotinylated primer that anneals to the universal tail, resulting in low density tiling across the region of interest (**Fig.5a**). The pooled and cleaned up asymmetric PCR reactions are now a probe set sufficient for hundreds of hybridizations. As shown at the bottom of **Fig.5a**, four probes ranging from 140-270 bp were tiled across 1.3 kb region with 300 bp or greater spacing for LOCK-seq on samples for Project MS2826 (**Table S5**). We obtained good uniformity of coverage across all samples, achieving 55x coverage for 99% of bases and 100x coverage for >90% (**Fig.5b-e**). Using this small panel, average read lengths were 2-4 kb across samples with a range of 2.5-17K mapped reads and capture efficiency of 10-20%. A MinION (R10.4.1) flow cell outputs 30-50 Gb of data, which averages more than 1M total 5-10 kb reads per run. This accommodates greater than 100 samples, with roughly 1000 reads per sample at a 10% capture efficiency, surpassing the efficiency obtained from Cas9-based capture methods (**Table 1**).

**Figure 5.**
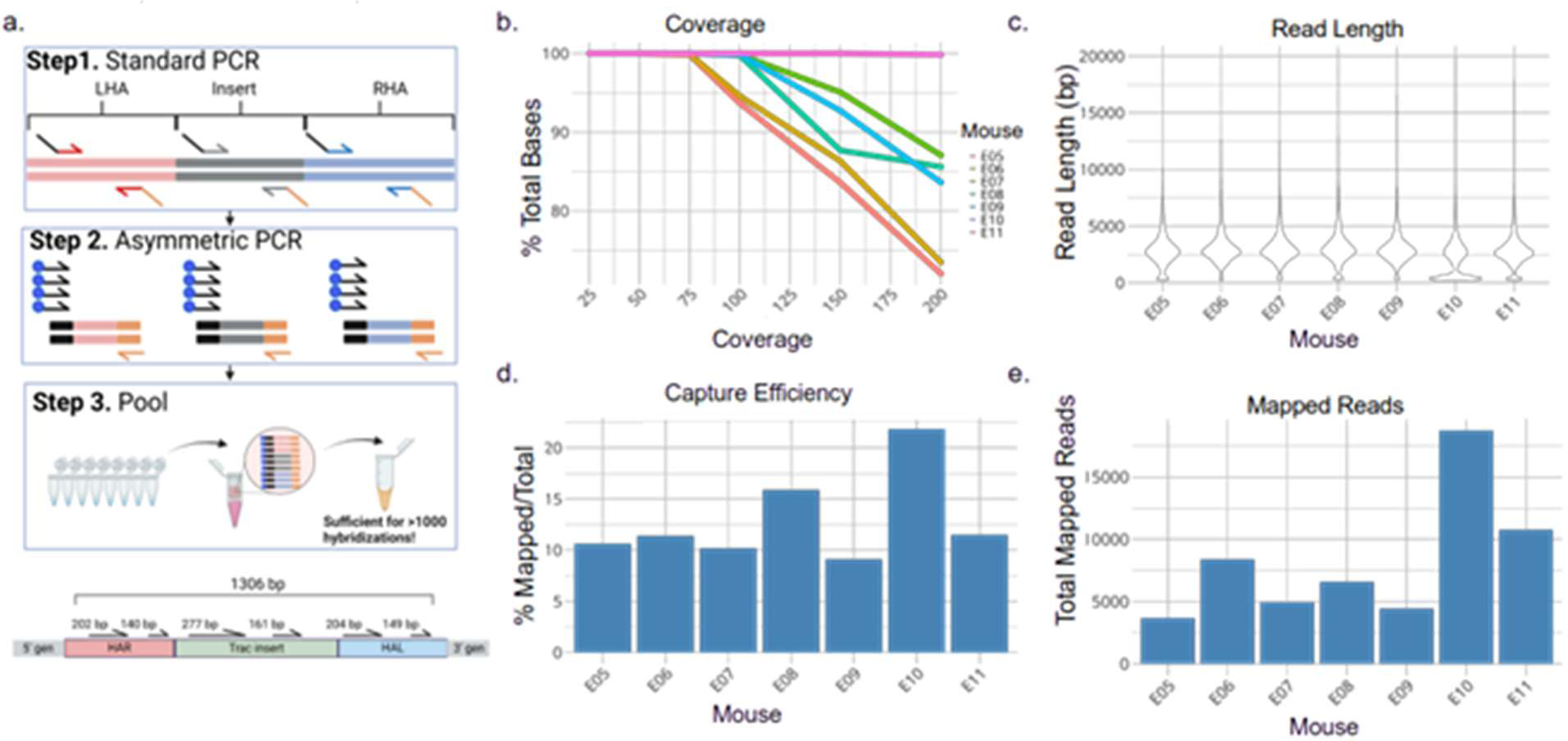
LOCK-seq metrics generated using in-house probes **a**. Schematic of probe generation and tiled probes across transgene **b**. Line graph of uniformity of coverage across samples **c**. Violin plot of lengths (bp) of mapped reads **d**. total mapped read counts and **e**. capture efficiency (mapped reads/total reads).

**Table 1.** LOCK-seq by using PCR generated probes significantly reduces costs while facilitating design flexibility, compared to multiple synthetic gRNAs for nCATS.

| Feature | nCATS (Cas9 RNP-based) | LOCK-seq (Probe-based capture) |
| --- | --- | --- |
| **Capture Mechanism** | Cas9/gRNA RNPs bind and cleave dephosphorylated genomic DNA to release regions of interest | Biotinylated DNA probes hybridize to target; captured fragments amplified via adapter-specific primers |
| **Sequencing Platform** | ONT long-read | ONT long-read |
| **Multiplexing Capacity** | Low (1-3 samples per MinION) | High (≤ samples on a MinION at $\sim 1000\times$ coverage for 5-10 kb read lengths) |
| **Versatility of Design** | Constrained by PAM sequences; multiple pairs of RNPs to improve efficiency | High flexibility: probes can be placed anywhere (insert, HA, ITR, flanks), unconstrained by PAM |
| **Capture Efficiency** | $\sim 1\%$ | $\geq 10\%$ enables deep coverage at target locus |
| **Number of Reactions** | $\sim 20$ reactions (2 nmol gRNA stock) | **Commercial panels:** $\sim 16$ captures **Asymmetric PCR in-house:** hundreds of captures possible |
| **DNA Input Requirement** | 1-5 ug genomic DNA | 250 ng genomic DNA |
| **Cost per 20 kb Target Locus** | $\sim \$500$ (Cas9 protein + synthetic gRNAs; 4 RNPs per locus) | **Commercial panels:** $\sim \$1000$ (120-mer biotinylated ssDNA panel, $\sim 120$ probes tiled at 30 bp spacing) **Asymmetric PCR in-house:** $\sim \$200$ (average 250 bp amplicons, $\sim 70$ probes tiled at 30 bp spacing) |

**Table S7** shows LOCK-seq runs on samples across five donor targets using probes generated with asymmetric PCR. The average read length was highly reproducible at 4.5 kb, and each sample had an enhanced on-target coverage of 1×10^4^-1×10^5^ reads. In these cases, hundreds of reads in the upper quartile of read length exceed the length of the donor from homology arm to homology arm, producing single reads that traverse the entire region of interest. This adds a higher level of support that the on-target knock-in sequence is contiguous with the genomic-specific flanking sequences, and therefore not an artifact of assembly.

### rAAV approach is successful in cancer and stem cell lines

Like mouse KI models, large insertions are sometimes required for cell line models. We applied the same strategy to engineer cancer and stem cell lines, including those that are too sensitive to plasmid DNA to survive transfection, such as mouse BV2 microglial cell line. We found rAAV added immediately after RNP nucleofection was 30-fold more efficient than a donor plasmid for inserting a cDNA-GFP into iPSC cells, measured for GFP by FACS a week after nucleofection (**Fig.S15a**), and confirmed by junction PCRs in single-cell clones (**Fig.S15b**). BV2 cells are highly sensitive to exogenous DNA and show low efficiency HDR when using ssODNs. A side-by-side comparison of ssODN-mediated KI in BV2 and N2a cells showed comparable gRNA activity in both cell lines but drastically different HDR rates, from barely detectable in BV2 cells to 10-30% HDR in N2a cells (**Fig.6a**). Likewise, plasmid DNA transfections often lead to nearly no survival of BV2 cells and undetectable HDR, making large cassette KI impossible to obtain. In contrast, we targeted BV2 with three 3-AAV sequential donors used for mouse projects to KI >6 kb and obtained >10% junction positive single-cell derived clones across all 3-AAV targeting projects (**Fig.6b** & **Fig.S16**).

**Figure 6.**
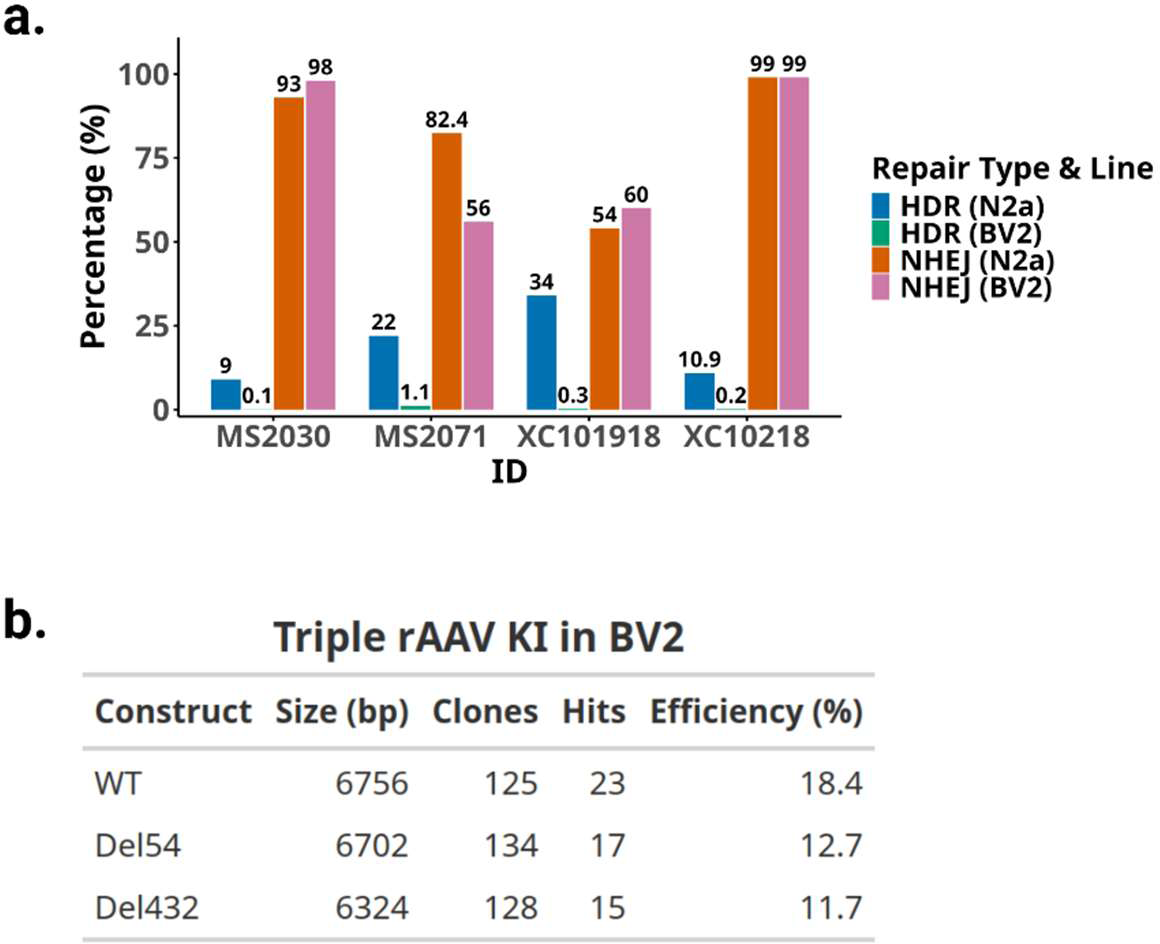
rAAV donors enabled knockin in BV2 cells that are highly sensitive to exogenous DNA exposure. **a**. Bar graph of HDR and NHEJ for BV2 and N2a cell lines using an ssODN template with CRISPR/Cas9. Projects MS2030 and XC10218, are an insertion of a loxP site with a BamHI site; MS2071 and XC101918, are a two base pair change from AG to CC and CC to TG, respectively. **b**. Table of KI efficiency of three related 3-rAAV projects, Project XCC113, in BV2 cells, reporting the insert size in each donor (bp) and total number of clones screened, resulting hits, and percent efficiency (hits/total clones).

### Detectable recombination between rAAV donors

Above we demonstrated that each double strand break generated by a Cas9/sgRNA cleavage of the chromosome mediates one HDR event, and up to three consecutive HDR events can occur in one delivery, achieving sequential rAAV-based KI. Single-stranded rAAV genome carrying a gRNA target site is not cleavable until KI occurs and the target site becomes double stranded, orchestrating the order of HDR events (**Fig.S17a**). We next sought to determine whether additional double-stranded breaks are needed for the second and/or third insertion events. In the absence of the second CRISPR/Cas9 complex, there is no double-strand break occurring after the first knock-in. It would require an interaction between donor 1 and 2, through inter-molecular recombination for example, to bypass the need of two independent HDR events for each half of the full-length composite sequence (**Fig.S17b**). Using two AAV donors, we tested project XCD47c, to knock in EF1a-Cre-ERT-T2A-luciferase-bGH pA signal, with or without the second RNP in HEK293T cells and iPSCs. Surprisingly, precise KI can occur at the target site with both donors without the need of a second double-stranded break, but much less efficiently (**Fig.S18**). The percentage of junction PCR positive clones for all junctions is 3-fold higher with two gRNAs in HEK293T and 5-fold higher in iPSC cells.

### LOCK-seq on cell lysates for quick, first-round clonal screen

We tested the feasibility of performing LOCK-seq on crude cell extract to significantly increase throughput, which would not be possible using the nCATS. Clones grown in a 96-well plate were extracted with proteinase K, treated with Tn5 transposase to add barcodes before PCR amplification, pooling for target capture and sequencing. We screened 59 iPSC clones on a single MinION flow cell and identified the same nine positive clones confirmed by junction PCRs (**Fig.S19**). These results demonstrate that LOCK-seq provides a practical and scalable approach for initial clonal screening.

### Optimization of Probe Design and Production for better capture efficiency

We next sought to optimize probe design to further improve capture efficiency by testing eight different probe designs in parallel (**Fig.S20a**). In theory, using probes that hybridize both strands could improve capture by enabling hybridization to both DNA strands with equal efficiency. To test this, we compared a top-stranded probe design to alternating top/bottom and double-stranded probe configurations (Groups A-C, **Fig.S20b**). All probe sets were 125 nt in length with 20 bp spacing between probes. Unexpectedly, the top-stranded probe design yielded the highest capture efficiency and coverage, outperforming both alternating and double-stranded designs by 1.4- and 3-fold in two different projects, respectively (**Fig.S20c-f, Table S7**).

We also examined three additional parameters affecting capture efficiency: (1) increasing the spacing between 125 nt top-stranded probes (Groups D and E, **Fig.S20b**), (2) extending probe length to 500 nt or 1,000 nt while maintaining 20 bp spacing (Groups F and G, **Fig.S20b**), and (3) additional biotin moieties (Group H, **Fig.S20b**). Increasing the probe spacing to 500 bp or 1,000 bp decreased performance, reducing capture efficiency by over twofold. Longer probes produced mixed results—decreasing capture for one target while improving it for another. However, increasing the biotin marks along the probe markedly improve capture efficiency for one target (6.7x) while modestly improving efficiency for the other (**Fig.S20c,d**). Notably, probes with additional biotin moieties outperformed other designs in achieving improved coverage across GC-rich regions (**Fig.S20e-h**). The overall length distribution of mapped reads was minimally affected by the parameters tested (**Fig.S20i-j**). However, when examining the longest top 10% of reads across conditions, probes more heavily modified with biotin skewed toward shorter reads (**Fig.S20k-l**). Our results support that the 125-nt probes at 20 bp spacing on the same strand with additional biotin moieties result in the best overall capture and coverage. For particularly GC-rich targets (>70%, >50 bp stretches), moderate addition of biotin during probe synthesis can improve coverage with the caveat of a slight shift towards shorter reads. The improved capture efficiency and coverage by using optimized probe design will allow higher levels of multiplexity and further reduce costs.

## DISCUSSION

Multi-kilobase KIs are critical for the generation of various types of mouse and cell line models, such as reporter lines, humanization, gene replacement, inducible overexpression and conditional KI. The use of rAAV donors for editing single cell embryos eliminates the need for microinjection when coupled with RNP electroporation. As importantly, both birth rate and founder rate are greatly improved. The caveat is the size limit of cargo to 4.7 kb including homology arms and ITRs, leaving maximum insert to be about 3 kb, which as we demonstrated here, can be overcome by consecutive insertions of two or three rAAV donors.

We generated >100 mouse models using 1-, 2- or 3-rAAV donors in a single-step delivery to insert up to over 6.6 kb. The most significant correlation is between KI efficiency and insert/homology length with one rAAV (**Fig.1d**). Previous work using plasmid donors supports a positive correlation between length of homology arms and HDR, with efficiency plateaus achieved at 1-2 kb, but the most marked effect is the poor performance of shorter arms (100-500 bp) that are <0.5 in insert/total homology, lowering HDR rates by half or more when compared to longer arms(28, 29). The homology arm length requirement further limits the insert size to around 3 kb. When more than one rAAV donor is used, donor 1 and/or donor 2 each bring in an additional gRNA site for recutting upon integration (**Fig.2a**). Efficiency was related to insert size, with efficiency decreasing as insert size increased. Nonetheless, most projects only required two mouse sessions to complete (∼200-300 embryos).

Even though we got overall excellent efficiencies, it is not failproof with two sessions to obtain founders for a given model. Sufficient homology length to insert size correlates positively to better efficiency. Additional contributing variables can include purity and accuracy of titer of AAV preps, gRNA cutting efficiency, sequence context at target site, variation in manipulation of the embryos during a given session, etc., or any combinations of the above. For example, among the failed projects using one or two donors, each had a project that the donor differed only by a point mutation from the one worked well, likely because of either rAAV quality/titer and/or arbitrary issues during the mouse session. Additionally, almost all multi-donor projects target the ROSA26 locus with very similar lengths of homology arms (∼800 bp each), yet the founder rate varies from 0-70%. Specifically, two of eight projects shared the same donor 1 and homology arms for donor 2 failed, yet the other six had founder rates ranging from 7.5% to 20%. rAAV titer is determined by qPCR of the ITR sequences, which does not always reflect the concentration of functional virus that can efficiently enter the nucleus and act as repair templates. Given the sequential nature of multiple insertion events, titer accuracy may be a leading contributing factor to variance in targeting efficiency.

The same rAAV strategy worked in iPSCs and cancer cell lines. Even though plasmid donors can be nucleofected, and editing is achievable, we often observed low survival post nucleofection and low targeting efficiency. In contrast, project XCB910b used a single rAAV to insert a GFP tag at the AAVS1 site, and showed a 10-fold increase in targeting efficiency compared to plasmid donor (**Fig.S15**).

As importantly, in cells like BV2 and T cells, where exogenous DNA is cytotoxic, rAAV donors make a drastic difference. Using three rAAV donors, BV2 can be edited to have 6.7 kb large insertion (**Fig.6** & **Fig.S16)**. And XCD47c, using two rAAV donors, reached about 14% positive clones in HEK293 cells and iPSCs (**Figs.S17-18**), compared to using a plasmid donor at less than 1% (not shown). Sequential insertion with AAV requires smaller plasmid constructs that are generally easier to synthesize than a single plasmid donor. Intermolecular recombination between monomeric circularized rAAV genomes have been reported as intermediate products that contribute to episomal persistence in muscle tissue, and rAAV genome concatenation is hypothesized to precede integration into the host genome (30, 31). Homology between rAAV genomes is suggested to mediate intermolecular concatenation via base pairing between ITRs, which account for the head-to-tail configuration of episomal rAAV genomes reported in liver and skeletal muscle(32, 33). Interestingly, when we left out the second gRNA in nucleofection of XCD47c, positive junctions were obtained in three out of 72 clones in HEK293 cells, compared to 10 out 75 with both gRNAs, implying the possibility of recombination between rAAV donors happens before HDR at the endogenous locus. The same targeting in iPSCs produced junction positive clones with both gRNAs, and without the second gRNA. Recent findings show that rAAV genome concatenation can bridge cis-acting transcriptional modifiers to rAAV expression cassettes delivered in trans, with nuclear concatemer formation detectable as early as 12 hours post-transduction(34). These concatemers may undergo further processing, providing a template for precise knock-in at a lower efficiency.

Genotyping becomes increasingly challenging with increasing insert size by junction PCRs, using one primer annealing to the target locus outside of a homology arm and the other in the insert to ensure site-specific insertion. Overlapping junction amplicons are more diagnostic but quickly become several kb in length with large inserts. Furthermore, multiple pairs of junction PCRs are necessary for genotyping sequential insertions (**Fig.S5**) and to confirm the final insert, yet some target sites and/or inserts are difficult to PCR amplify, and positive junctions do not guarantee the two junctions were amplified from the same allele, i.e. partial insertions can lead to false positives. Additionally, CRISPR-mediated targeted integration processes can introduce undesired by-products, such as random integration of donor templates, homology-independent insertion in full or partially into off-target cleavage sites, on-target large deletions and potentially chromosomal translocations, none of which can be detected by junction PCRs. Genotyping can be a time and resource-consuming obstacle for establishing sophisticated research models and potentially skewing data obtained if genotyping is inaccurate.

We developed cost-effective, high-throughput LOCK-seq to not only screen for accurate KI allele but also detect large indels and donor random integration, providing comprehensive characterization of the genome. By using less than 1 μg of genomic DNA per sample, clonal cell populations and founder animals with the same or different targets can be multiplexed, enriched by target capture using biotinylated probes designed against the insert, homology arms and flanking genomic sequences and loaded into the same flow cell. A MinION flow cell typically produces 20-30 Gb of total output that translates into thousands of reads per sample, sufficient even for regions that may have lower sequencing depth due to high GC content, sequence context, and less-than-optimal probe performance. With an average turnaround of a typical run being just under a week, LOCK-seq enables quick identification of knock-in clones and founder animals, thus minimizing culture time and animal husbandry costs.

LOCK-seq significantly reduces per sample cost compared to nCATS and related methods for three main reasons. First, LOCK-seq libraries result in more productive sequencing events on the MinION than those of nCATS. nCATS is amplification-free and relies on CRISPR/Cas9 to bind and expose the DNA ends of select loci for ligation of the motor protein required for ONT sequencing. We hypothesize that even though the majority of DNA loaded on the flow cell is not competent for sequencing, it can transiently interact with pores or adapter ligated molecules, interfering with sequencing. On the other hand, LOCK-seq achieves greater than 10-fold increase in capture efficiency via hybridization-mediated target enrichment and higher output per flow cell with significantly more efficient ligation between the PCR amplicons of enriched sequences and the motor protein to result in greater than 100-fold increase in coverage than nCATS. Second, LOCK-seq enables low cost and flexible probe generation. We first used custom synthetic oligo probes labeled with 5’ end biotin, which incurs a higher reagent cost. We then showed that probes generated by asymmetric PCR are versatile and efficient. Panels of 5-10 asymmetric PCR probed (<500 bp in length, 5’ biotinylated, spaced ∼20 bp apart along the flanking genomic regions, homology arms and the insert) can be produced in under an hour. Each set of asymmetric PCRs generates enough probes for hundreds of hybridizations. In contrast, nCATS and related methods require one or more gRNAs that are specific to the target (**Table 1**). To further improve capture efficiency, we tested different probe length, density, strand distribution and with additional biotin-modified bases incorporated (**Fig.S14**). Our data shows that probes of 125 nt, with 20 bp spacing on the same strand with additional biotin-modified internal bases resulted in the highest capture efficiency with minimal increase in probe costs. The preference for probes designed against the same strand may imply the impact of probe density, whereas more than one biotin/streptavidin interaction/probe seems to be needed for maximal capture of multi-kilobase DNA molecules. There is still room to further optimize probe design and production. Finally, fragmentation and barcoding are straightforward. Using Tn5 or Covaris g-TUBEs for fragmentation, barcodes can be ligated to the DNA to facilitate multiplexed target capture. Tn5 fragmentation typically yields post-hybridization reads of 1-2 kb, while Covaris g-TUBEs yield 4-6 kb-long reads, sufficient to obtain single contiguous reads with coverage across the entire transgene and its flanking genomic sequences. In cases where read lengths were shorter than the transgene size, assembly tools like Canu can be used to generate a high-quality consensus sequence, proving the presence of an insertion event, or lack thereof. Both Tn5 tagmentation and the Covaris g-TUBE fragmentation work for LOCK-seq. The latter only requires a one-minute centrifugation and yields longer fragment lengths, averaging around 10 kb. Tn5 tagmentation has higher throughput, is more suitable for an initial screen and can use cell lysates generated with Quick Extract instead of column-purified genomic DNA preparations (**Fig.S19**). We found that it is critical to use low cycle numbers for both PCR steps to achieve longer read length, 6-8 cycles for barcoding, and 16-18 cycles for post-hybridization PCR.

In addition to on-target insertion events, which can be precise or with imperfections (usually various sized deletions in the insert), we used LOCK-seq to identify concatenation of donors at the target site and random integrations of donors as well as genomic locations of the randomly inserted donors (**Fig.5**). Both random integration and concatenation of the donors potentially skew phenotypes of the cell or mouse models. Using LOCK-seq to screen and choose correct clones and founders are critical to the precision of the research models. LOCK-seq is agnostic to the format of donors used, and we have had equal success at genotyping models generated with plasmid donors. And the method should be readily adaptable to be used for genomic localization of transgenes created by various means, such as via lentiviral integration and transposition (35, 36), and for detection and confirmation of genomic rearrangement events. The upper limit of insert size for reliable detection in our hands is currently at 10-20 kb.

The presence of artifacts due to fusion reads was explored and found to be <2%. The difference between artifacts and real fusion events is that real fusions have a distinct junction breakpoint that is supported by multiple independent reads, as opposed to artifacts which lack a consistent fusion junction breakpoint. When mapping to the whole genome, split reads will inevitably arise at low complexity sites, so we use unique sequence elements as anchors when mapping, ensuring that multiple independent reads traverse these regions. This increases confidence when curating the data. Critically, we follow-up with PCR to confirm putative off-target sites identified from LOCK-seq are not sequencing artifacts.

Nucleases potentially have off-target activities, which are not fully understood or well predicted, possibly leading to additional insertion of the donor template, with or without relying on homology(37, 38). Using LOCK-seq, recurring random insertion sites among different founders and clones are likely to be off target sites, instead of truly “random”. So far, we have not detected potential off-targeting cleavage-mediated insertion sites in either mice or cell lines.

We have demonstrated that LOCK-seq works well to characterize the post-edit genome. Yet several limitations of LOCK-seq exist. First, the need for two rounds of low-cycle PCRs restricts read length (**Fig.S7a**) and renders LOCK-seq to be more susceptible to coverage dropouts in GC-rich regions, which can be partially overcome by incorporation of more biotinylated bases in the probe (**Fig.S20**). nCATS, on the other hand, is PCR-free, has longer read length and preserves methylation readout. With our current conditions, we obtained read lengths of 4-6 kb on average, which limits the chance to obtain individual reads spanning from the genomic region outside one homology arm, across large inserts, especially with concatemers, to the genomic region outside of the other homology arm. Optimization of fragmentation methods, PCR conditions and probe design can yield and enrich longer fragments more efficiently. Additionally, it is difficult to apply this method to a transfected pool shortly after transfection, given the abundance of rAAV genome or plasmid donor, compared to the copy number of genomic targets. Finally, analysis of undesired events requires manual curation. We are currently developing a more automated pipeline to facilitate analysis. Even though the advantages of LOCK-seq compensate for these limitations enough to make it a powerful method, further improvements can enable LOCK-seq to be even more valuable.

In conclusion, we have shown that large, multi-kilobase KIs can be achieved efficiently in mouse embryos and cell lines via HDR using more than one rAAV donor and CRISPR/Cas9, and that LOCK-seq reliably verifies on-target, precise KIs and identifies undesired partial KIs, donor concatenation and random integrations, enabling accurate selection of founders and clones to ensure model precision.

## SUPPLEMENTARY DATA

All supplementary data are available via the Supplementary Materials link.

## AUTHOR CONTRIBUTIONS

Monica Sentmanat: Conceptualization, Formal analysis, Methodology, Investigation, Writing—original draft, Writing—review & editing. Zi Teng Wang: Investigation. Evguenia Kouranova: Investigation. Samuel Peters: Investigation, software. Wan Chin Chen: Investigation. Jed Lin: Investigation, software. Yong Miao: Investigation. Michael White: Investigation. Mia Wallace: Investigation. Xiaoxia Cui: Conceptualization, Methodology, Investigation, Writing—original draft, Writing—review & editing.

## DATA AVAILABILITY

The ONT data has been deposited in GenBank under BioProject number PRJNA1282668.

## ACKNOWLEDGEMENTS

We thank Joe Dougherty, Ben Humphreys, Gaya Amarasinghe and Daisy Leung and for comments and suggestions, Sophia DeGeorgia, Robert Fulton, Catrina Fronick, Paul Cliften for technical assistance, Mingjie Li at Hope Center Viral Core, Washington University in St. Louis, for rAAV preps and Jessica Hoisington-Lopez at DNA Sequencing Innovation Lab at The Edison Family Center for Genome Sciences & Systems Biology, Washington University in St. Louis for amplicon NGS analysis.

