## Supplemental Figures for "Efficient Multi-Kilobase Knockins in Mice and Cell Lines using CRISPR/Cas9 and rAAV Donors with Unbiased Whole-Genome characterization by LOCK-seq"

**List of supplemental tables**

**Table S1.** List of gRNA and donor sequences used in this study.

**Table S2.** List of genotyping primers used in this study.

**Table S3.** List of 120-mer biotinylated oligo probes used for hybridization in this study.

**Table S4.** List of primers used for in-house probe generation.

**Table S5.** Stats for single rAAV donor projects.

**Table S6.** Stats for projects using two or three rAAV donors.

**Table S7.** Mapping statistics for LOCK-seq in this study.

**Table S8. Reference sequences to query for fusion reads in Fig.S14.** Full sequence of references used for **Fig.S14**.

**Figure S1**

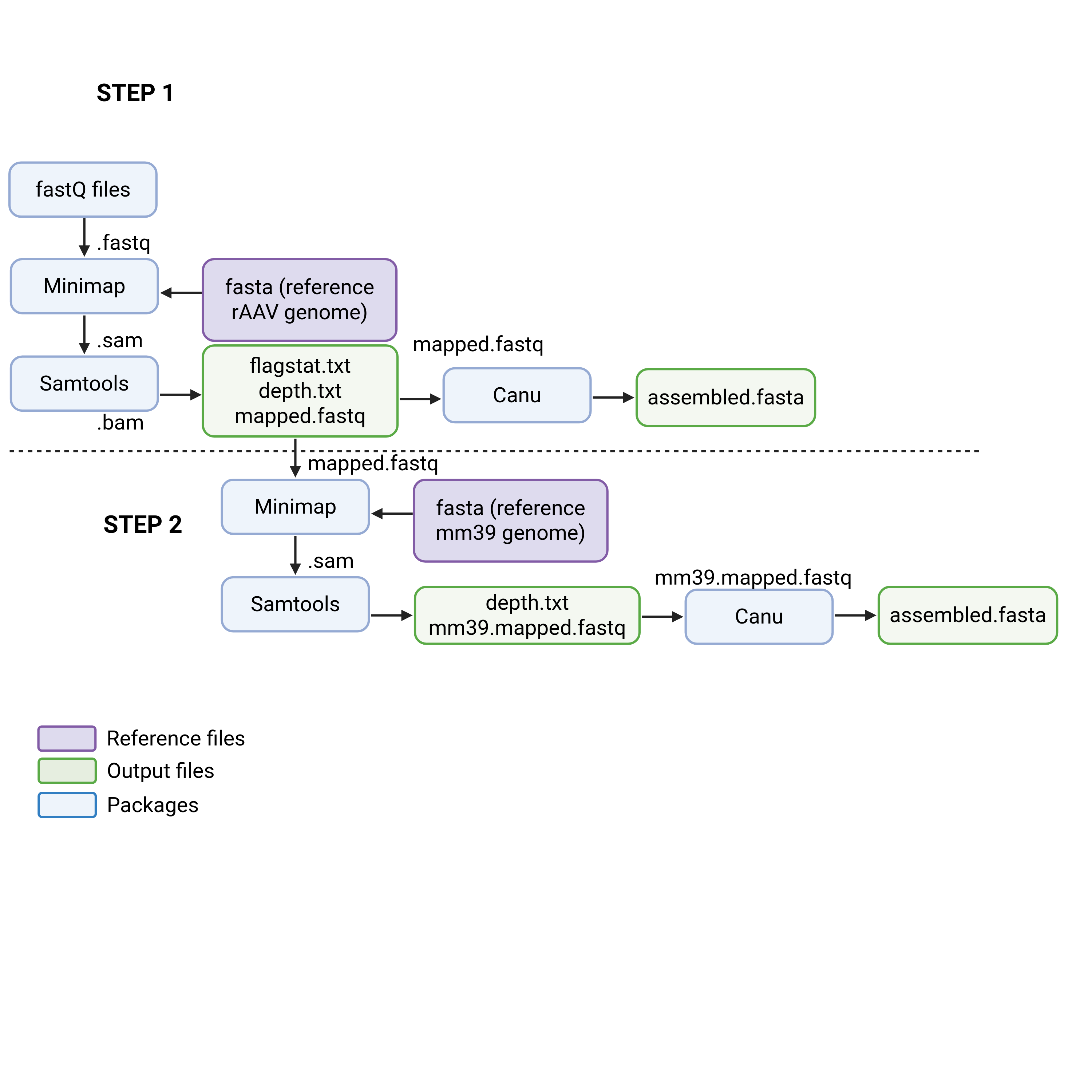

**Supplemental Figure S1.** First, reads are aligned to the donor construct (including homology arms), then mapped reads (fastq) are pulled to assemble (fasta) the on-target editing events using Canu. Not all editing events are precise or on-target. When indels or mutations are present, they are evident in the assembly. These donor-specific mapped reads (fastq) will contain overhangs that map to the flanking genomic regions if an on-target event occurred or will have genomic regions that map elsewhere if a random integration occurred. In the second step, the mapped reads (fastq) are mapped to mm39 to determine where in the genome donor-specific reads map. If both on-target and random integrations occurred, high read depth at the on-target site (>40% of total mapped reads) will be identified, in addition to high read depth at the candidate random integration sites (>20% of total mapped reads).

**Figure S2**

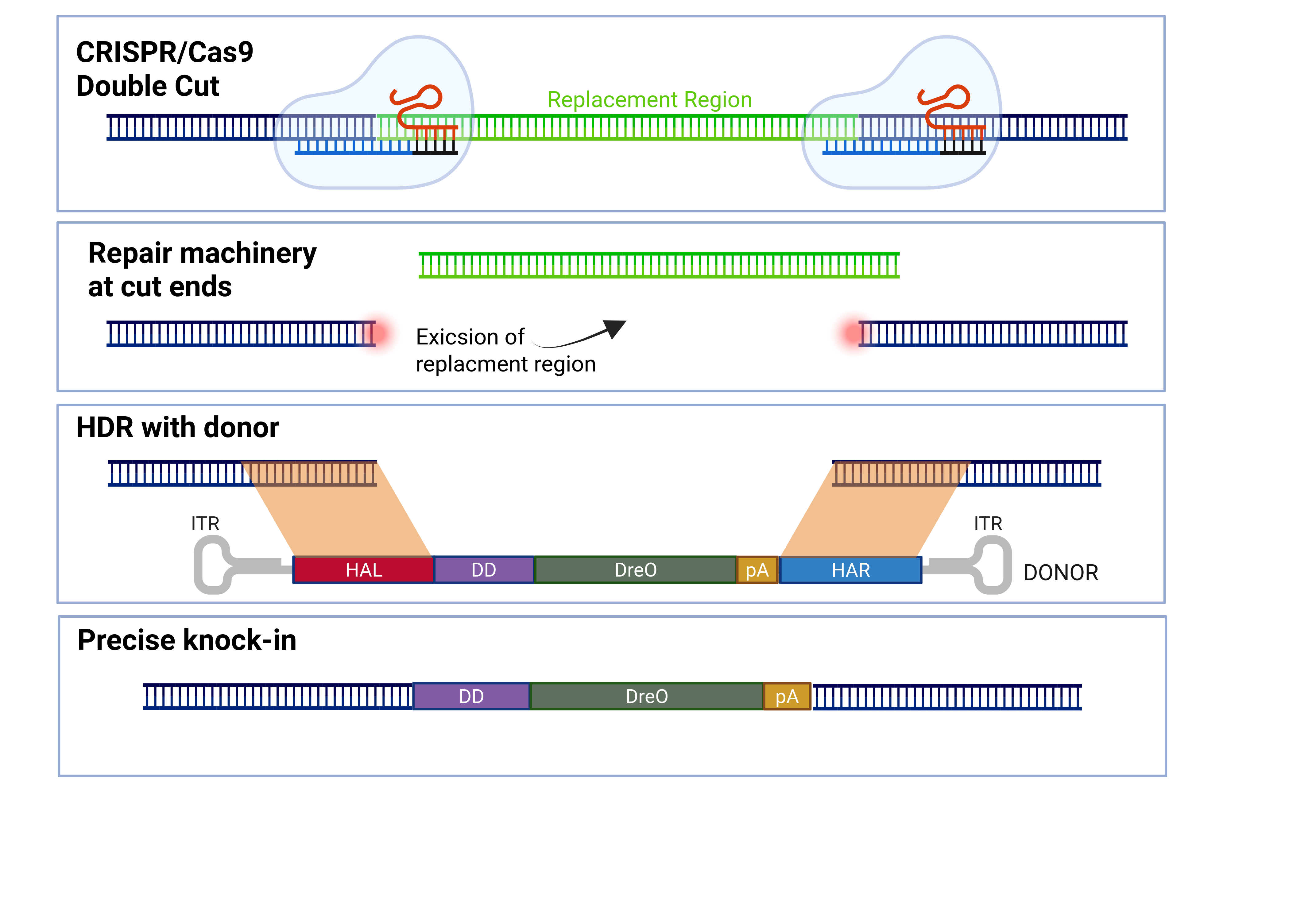
**Supplementary Figure 2.** The schematic of gene replacement strategy used in this study where two gRNAs cleave at the boundaries of the region marked for replacement, i.e. concomitant deletion and knock-in (e.g. MS2742/MS2746 listed in **Table S5**). The homology arms are designed to flank the replacement region, outside of both cut sites and with 500-1000 bp in length each.

**Figure S3**

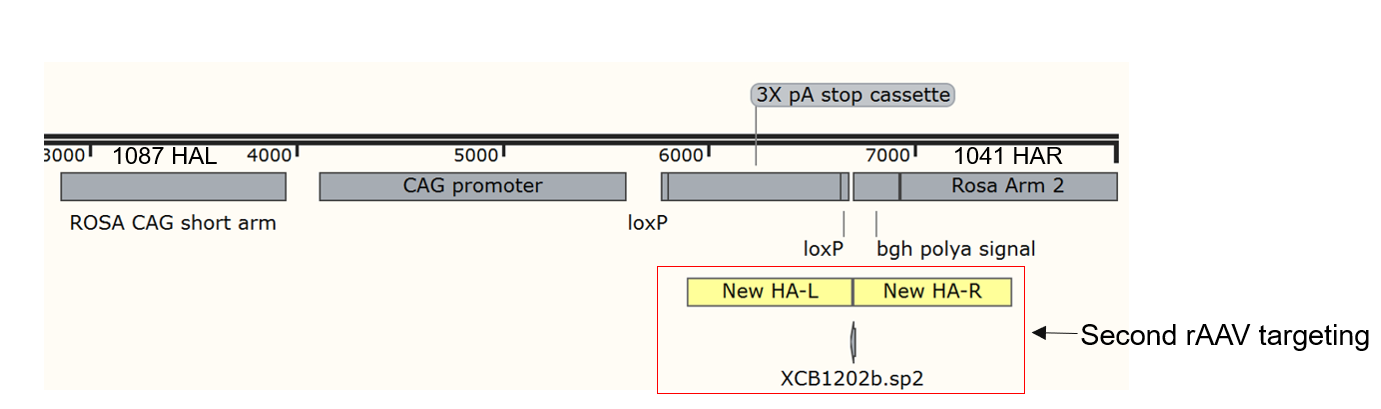

**Supplementary Figure 3.** Schematic of conditional expression configuration in the ROSA26 locus, inserted with the first rAAV donor of the cassette containing the CAG promoter, floxed 3xSV40 poly A and a BGH poly A signal. The cDNA to be expressed will be inserted into the XCB1202b.sp2 gRNA site on a second donor. Sperm from males harboring this “recipient” site was frozen down to be used in IVT for fertilized eggs so that only rAAV donors delivering cDNAs are needed for future targeting.

**Figure S4**

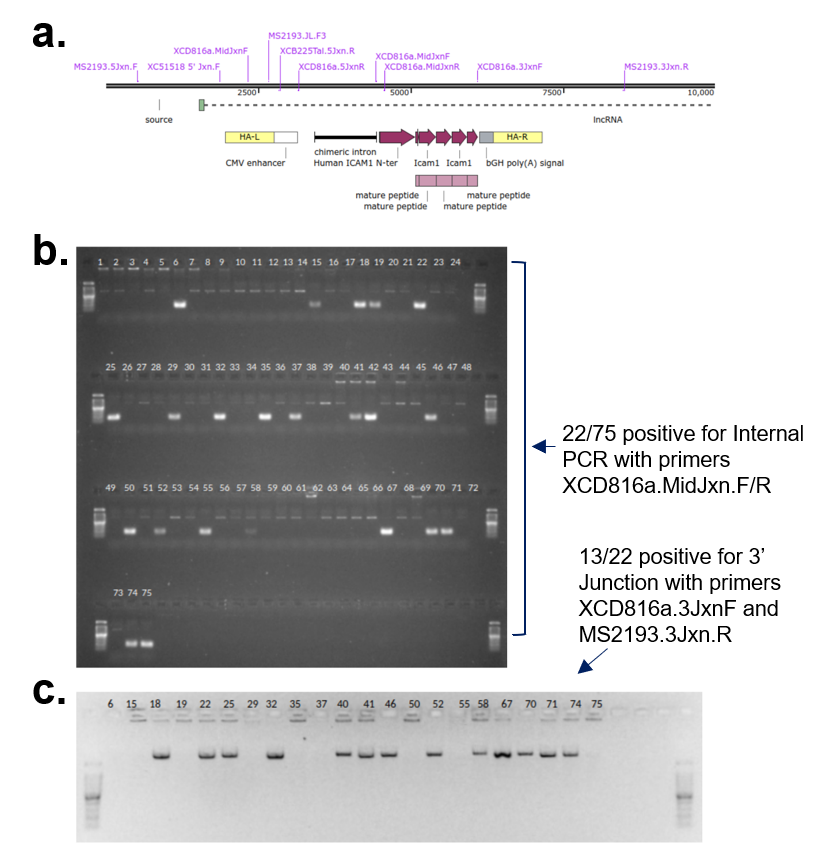

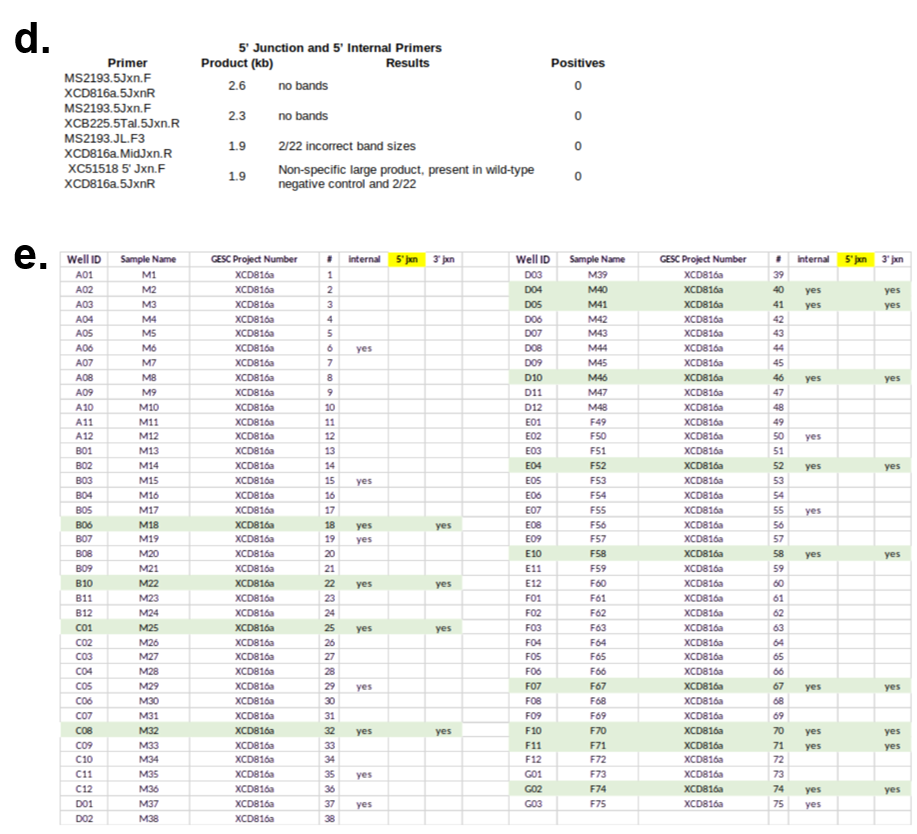

**Supplementary Figure 4** PCR-based screening for multi-kilobase KI of XCD816a with single donor junction detected across several animals. **a**. Schematic of rAAV donor and junction PCR designs of a “failed” project. **b.** Gel image of 22 animals positive for internal PCRs specific to cDNA transgenes, and **c.** 13/22 of internal PCR positive animals also positive for 3’ junction PCR. **d.** Table of 5’ junction PCRs attempted and negative results across 22 animals positive for internal and/or 3’ junction. **e.** Summary table of PCR results illustrating a labor intensive and inconclusive screening odyssey.

**Figure S5**

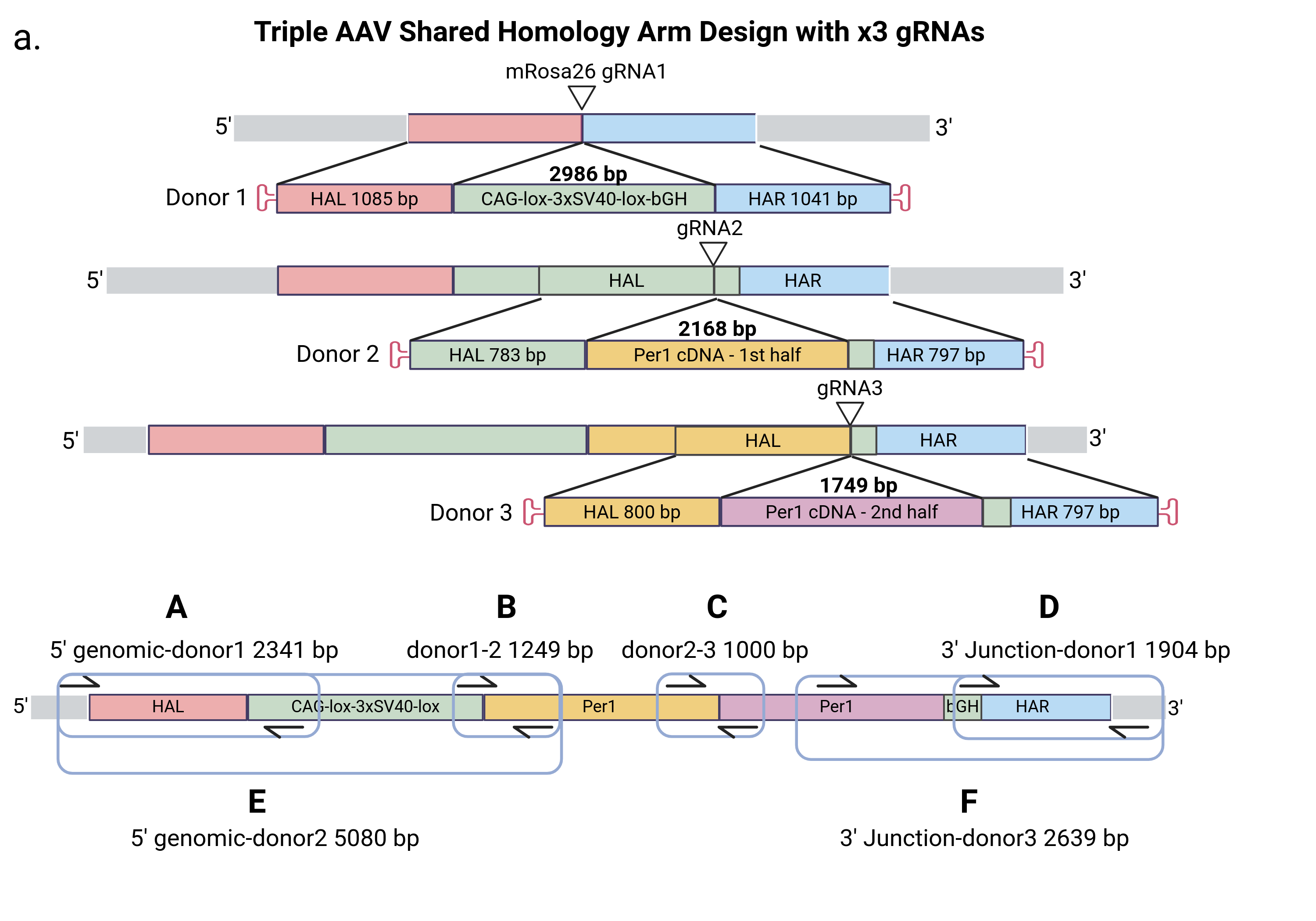

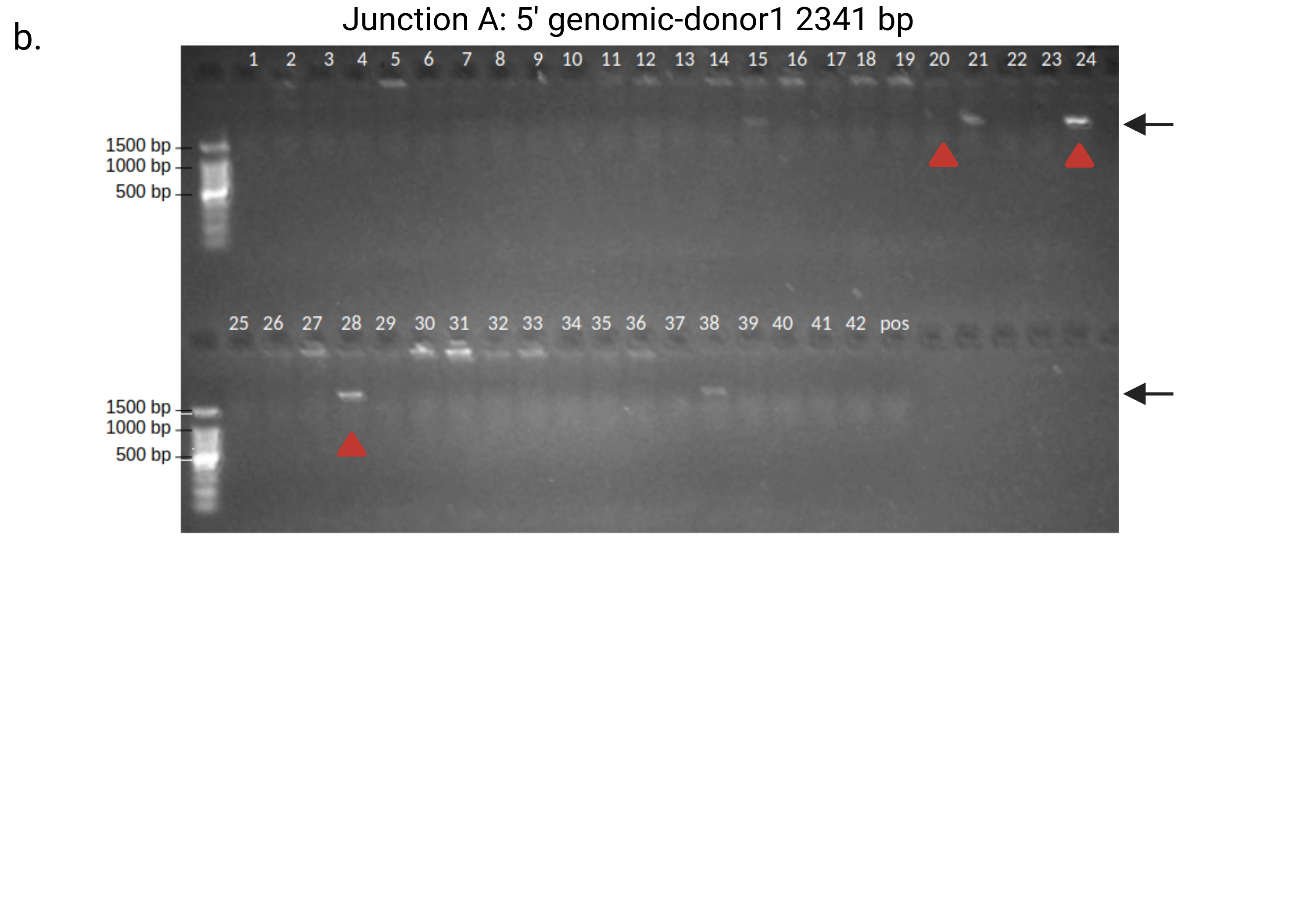

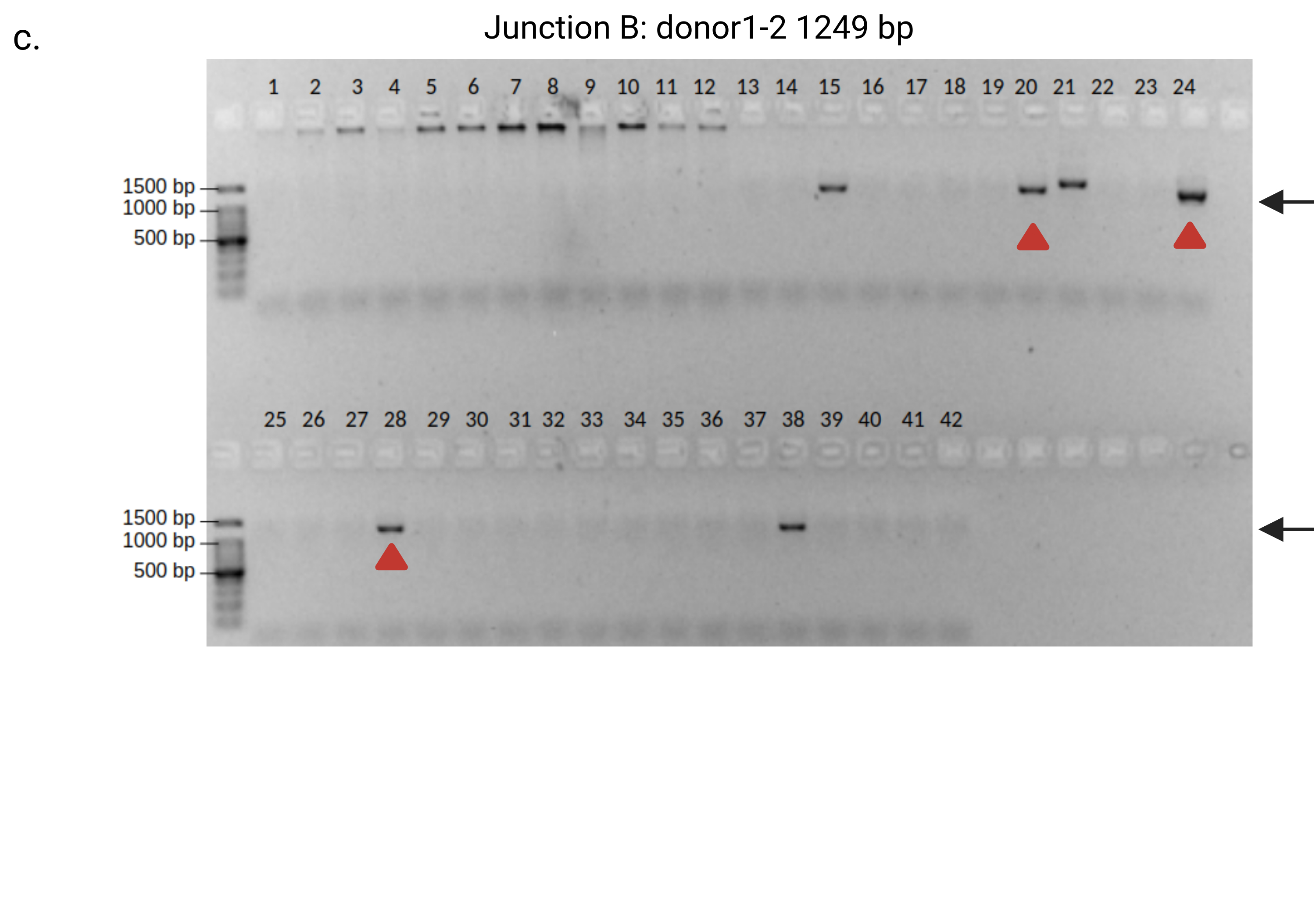

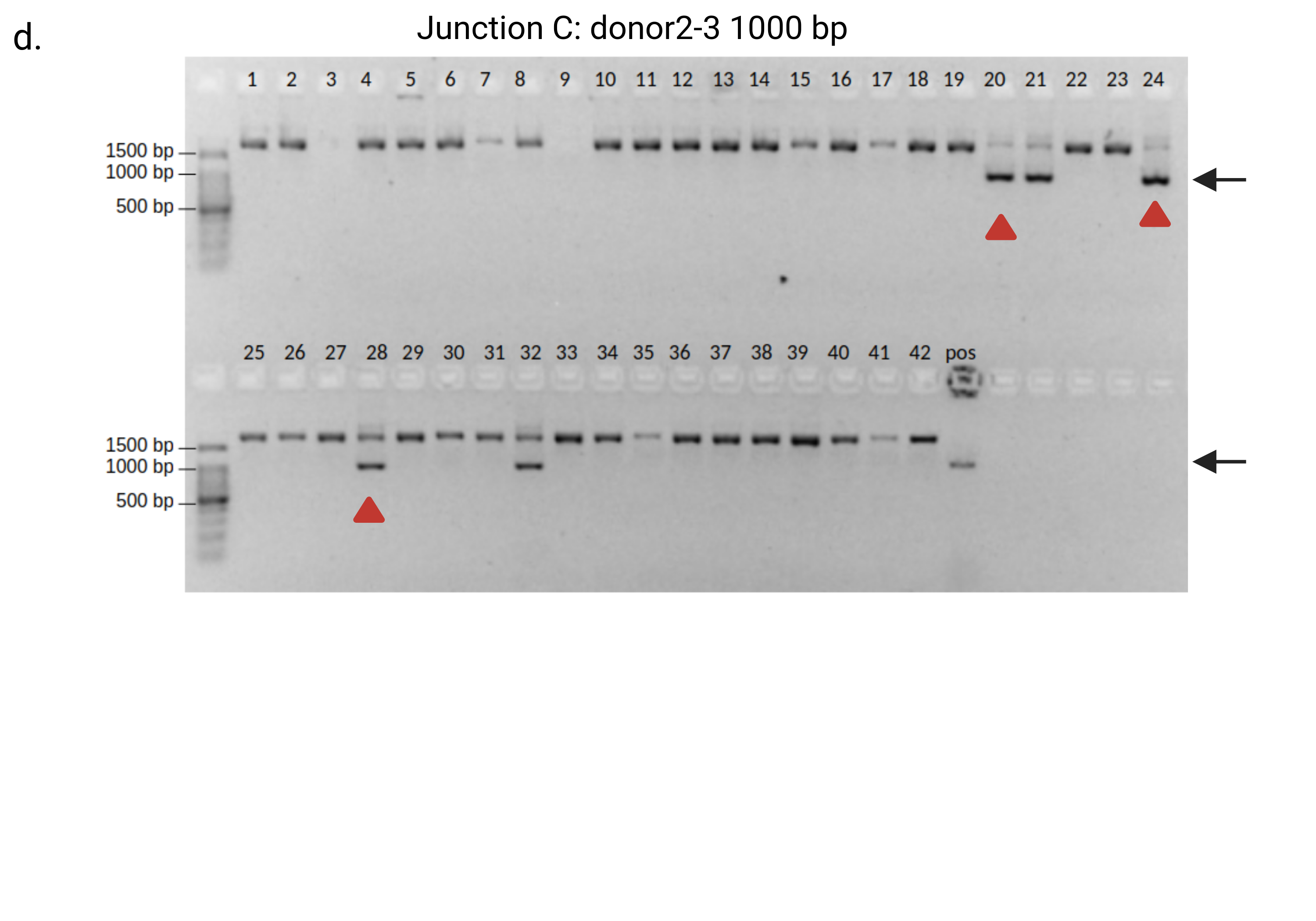

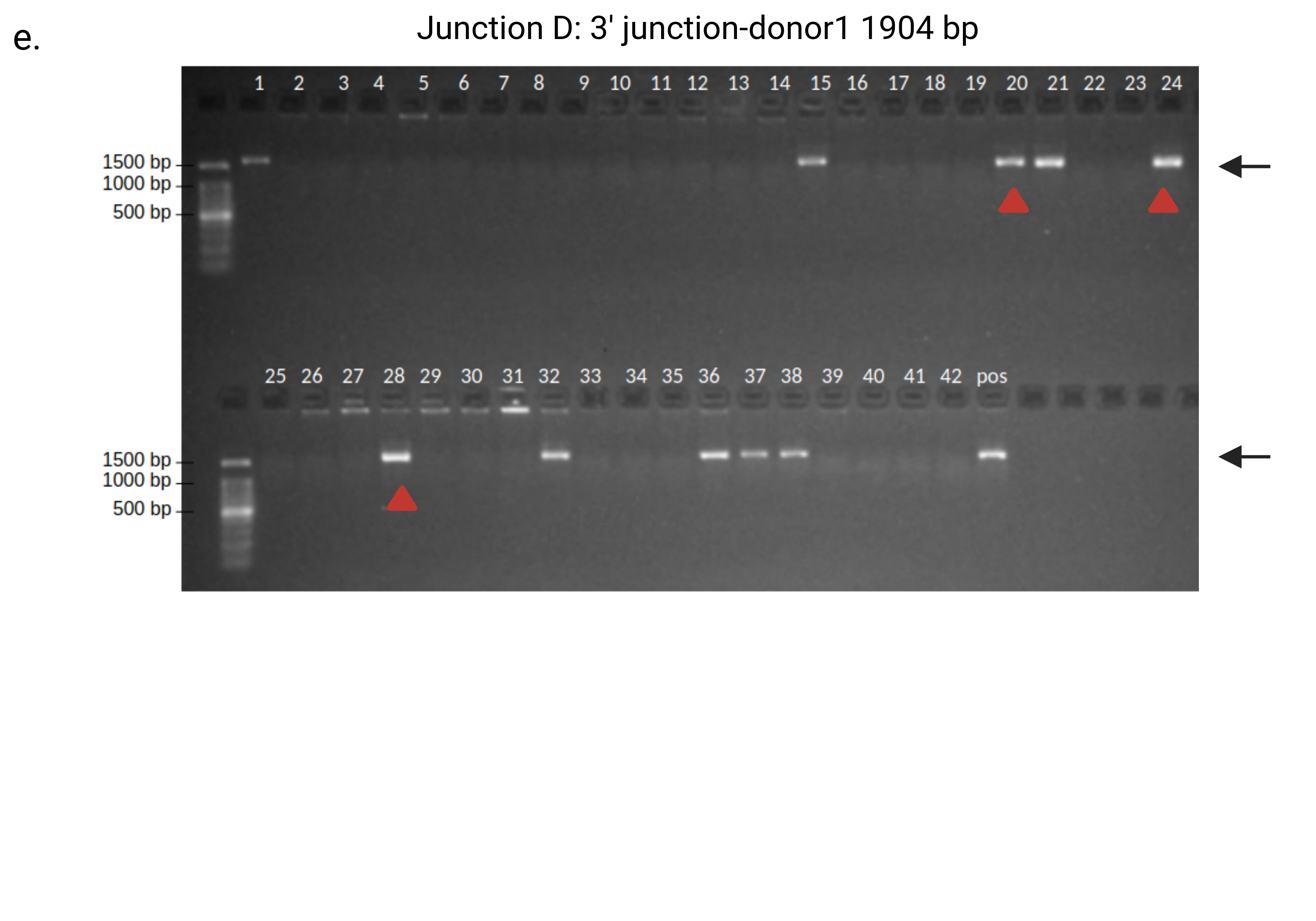

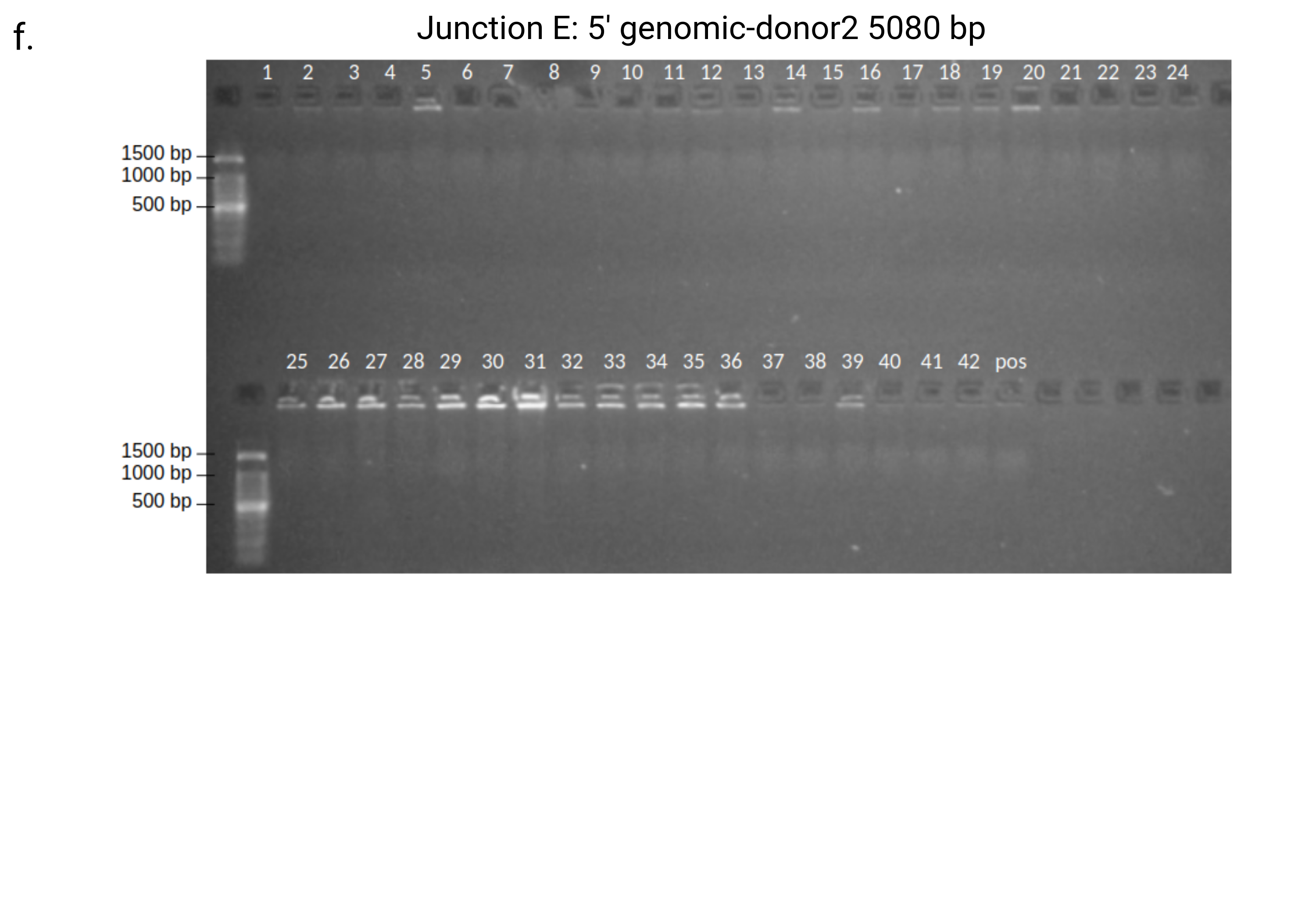

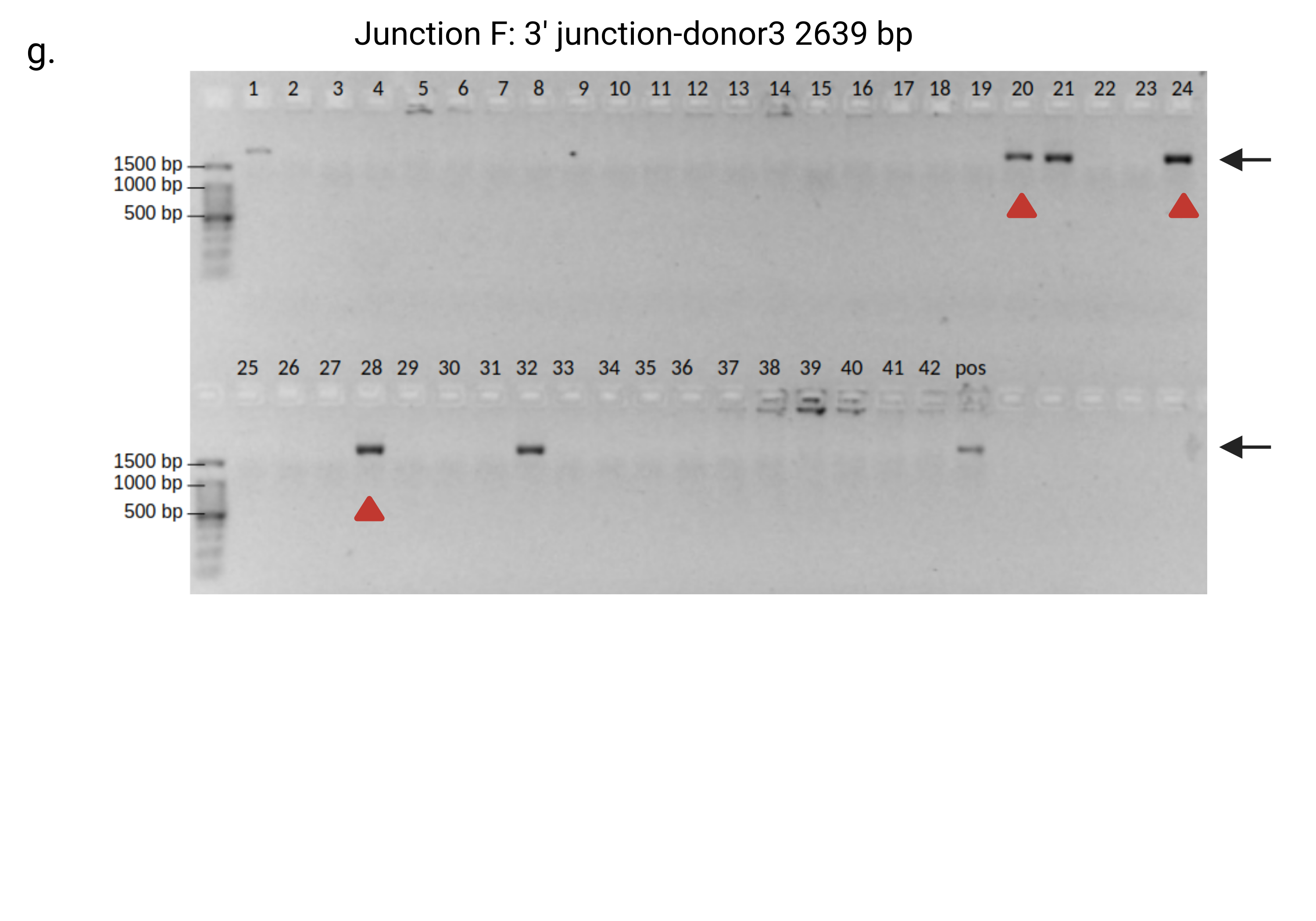

**Supplementary Figure 5.** Six PCRs are needed for screening founder animals after a 3-rAAV KI. a. Schematic of x3 rAAV donor KI for XCC113-Wt (see **Table S6**, 7.14% at 3/42 F0 pups positive) showing left and right homology arms on either side of the three donor inserts (green, yellow, purple), upstream and downstream (grey) flanking genomic regions. Primers (arrows) for two internal (B-C) and four junction PCRs (A, D, E, F) are shown. b. Gel images of PCR products for F0s screened by each set of primers. Arrows point to expected band sizes. Junction E failed to produce a positive band for positive control, suggesting further optimization is required. This junction was not used to identify putative hits. Animal 20 was negative for junction A (*) but was maintained as putative hit since positive control also failed to amplify. PCR-based screening can result in incorrectly sized bands (**, junction C high background), double banding (junction C hits), weak bands (^), and inconclusive results across difficult to amplify regions. Additionally, it is hard to determine whether all these products were amplified from the same allele.

**Figure S6**

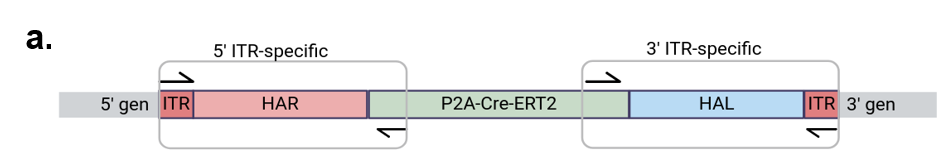

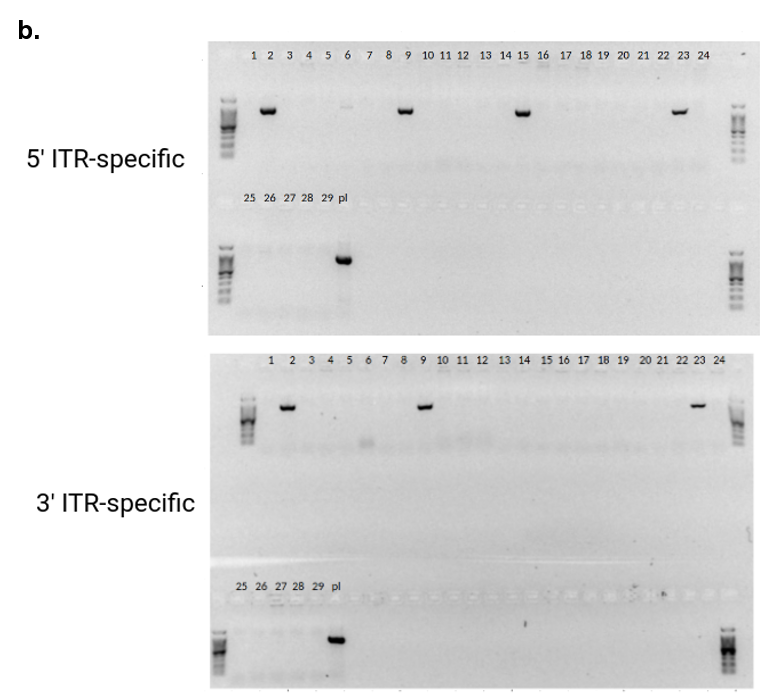

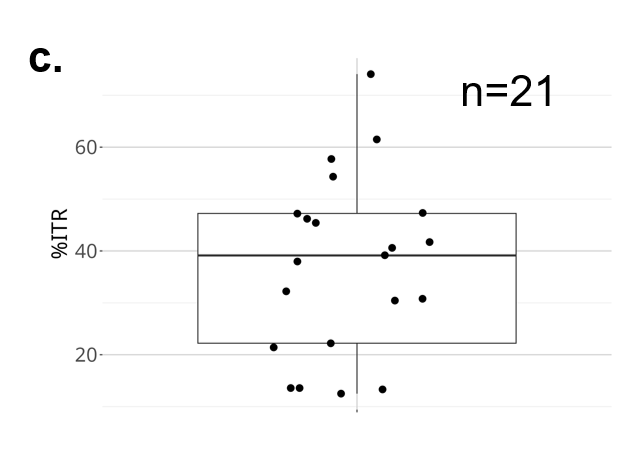

**Supplementary Figure S6.** Detection of rAAV random integration by PCR. a. Schematic of x1 rAAV donor KI showing left (red) and right (blue) homology arms on either side of donor KI (green), upstream and downstream (grey, 5’ gen and 3’ gen) flanking genomic regions. Primers (arrows) for 5’ and 3’ ITR-specific amplification are shown. b. Gel images of PCR products for junction positive F0s screened by each set of primers. c. Boxplot of percent positive for ITRs of total junction positive animals for 21 projects.

**Figure S7**

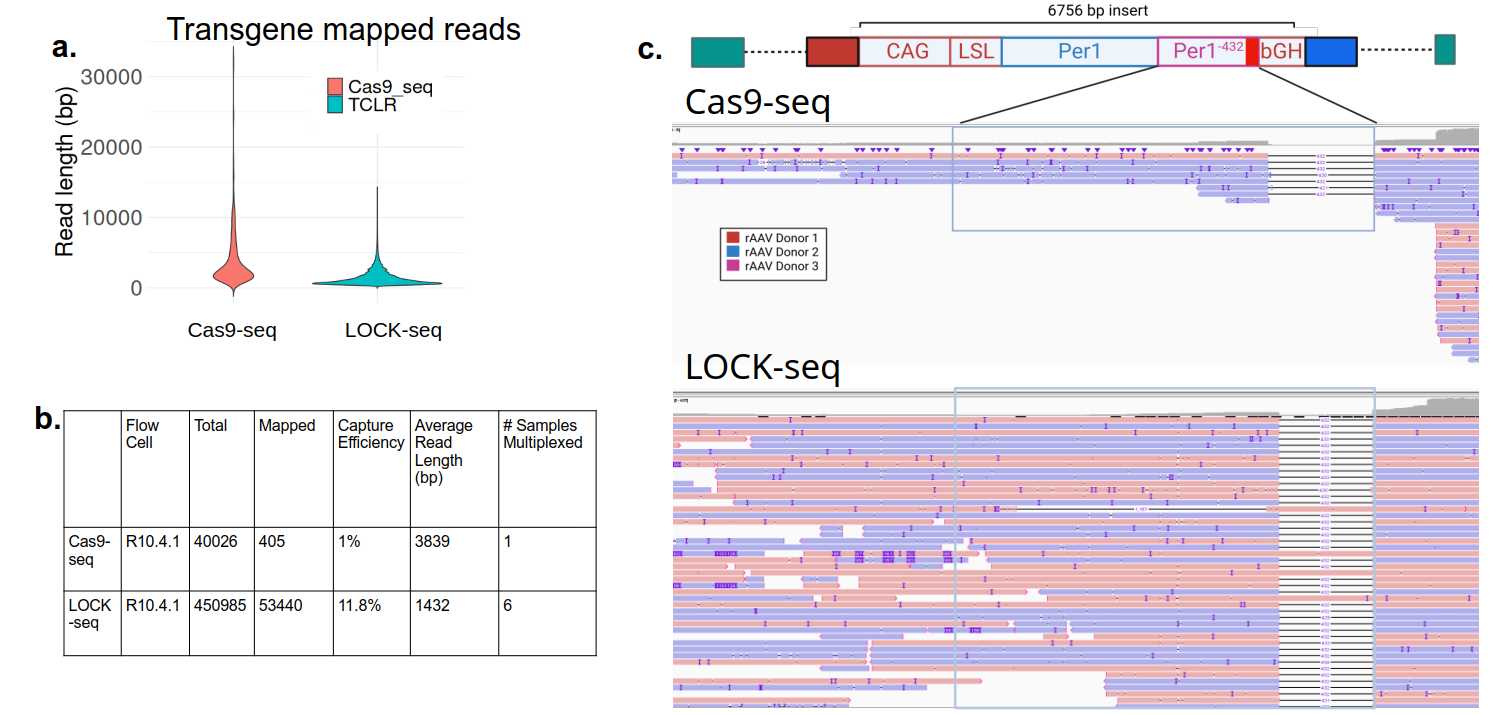

**Supplementary Figure 7.** The three rAAV KI mouse samples analyzed by Cas9-seq **a.** Violin plot of mapped read lengths from Cas9-seq compared to LOCK-seq. **b.** Table of flow cell (R10.4.1) output for Cas9-seq and LOCK-seq, capture efficiency (mapped reads/total), average read length and number of samples multiplexed on a single flow cell. **c.** Screenshot of IGV browser loaded with donor reference sequence and Cas9-seq vs. LOCK-seq bam files zoomed to expected 432 bp deletion. Samples are from project XCC113, of which LOCK-seq data shown in **Figure 4d**.

**Figure S8**

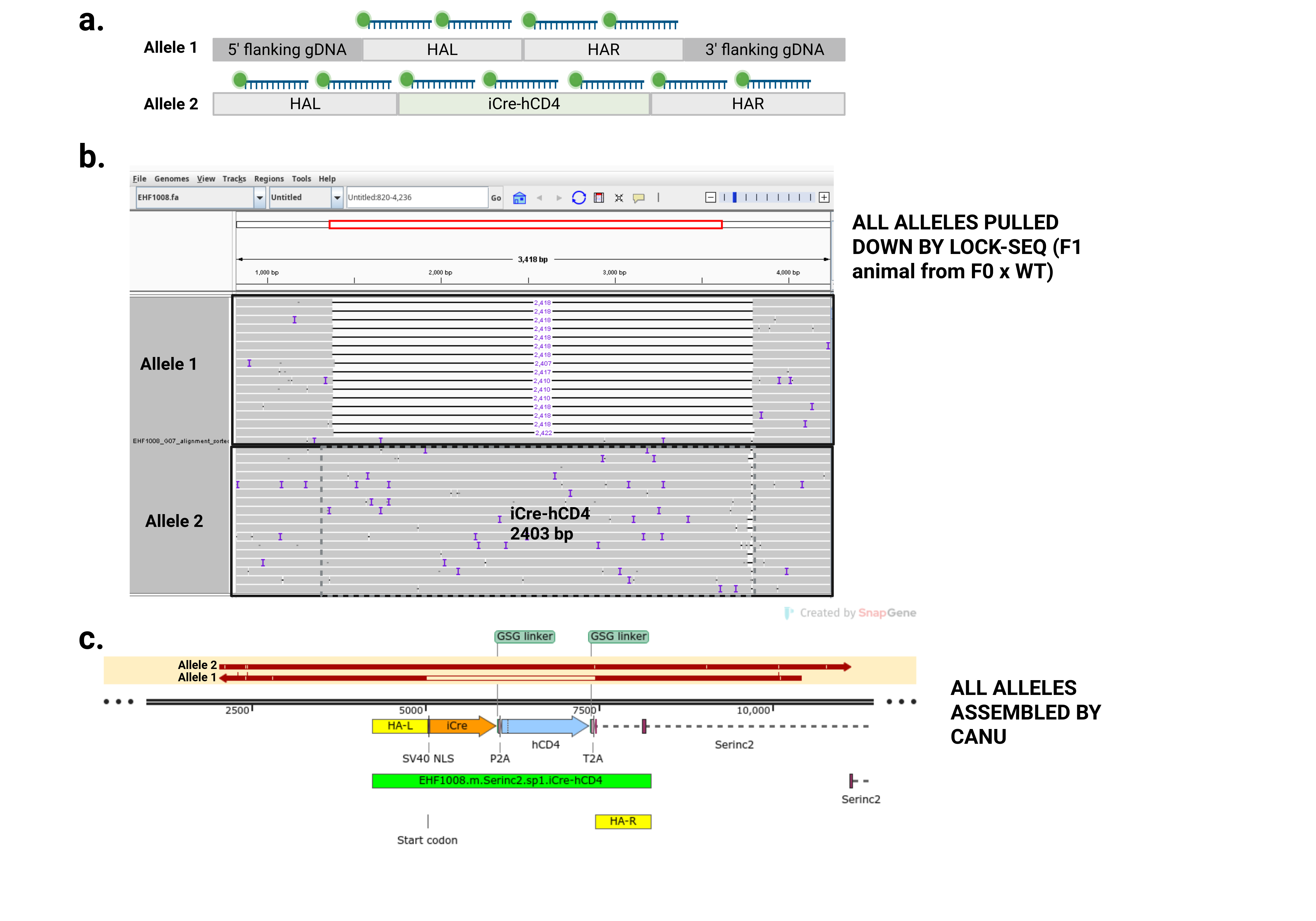

**Supplementary Figure 8** LOCK-seq probe design for pulling down all alleles, agnostic to genotype **a.** Cartoon of typical probe design spanning the full-length donor including homology arms (e.g. Allele 2). The homology arms are present on all copies since they are designed to target an endogenous genomic locus. **b.** IGV screenshot of mapped reads show the non-targeted allele (or wild-type) is missing the 2.4 kb insert but has homology arms present while the knock-in allele has the 2.4 kb insert (dashed grey box for Allele 2). **c.** Snapgene screenshot for alignments of contigs for both alleles of a heterozygous KI animal resulting from the breeding of a mosaic F0s to wild-type.

**Figure S9**

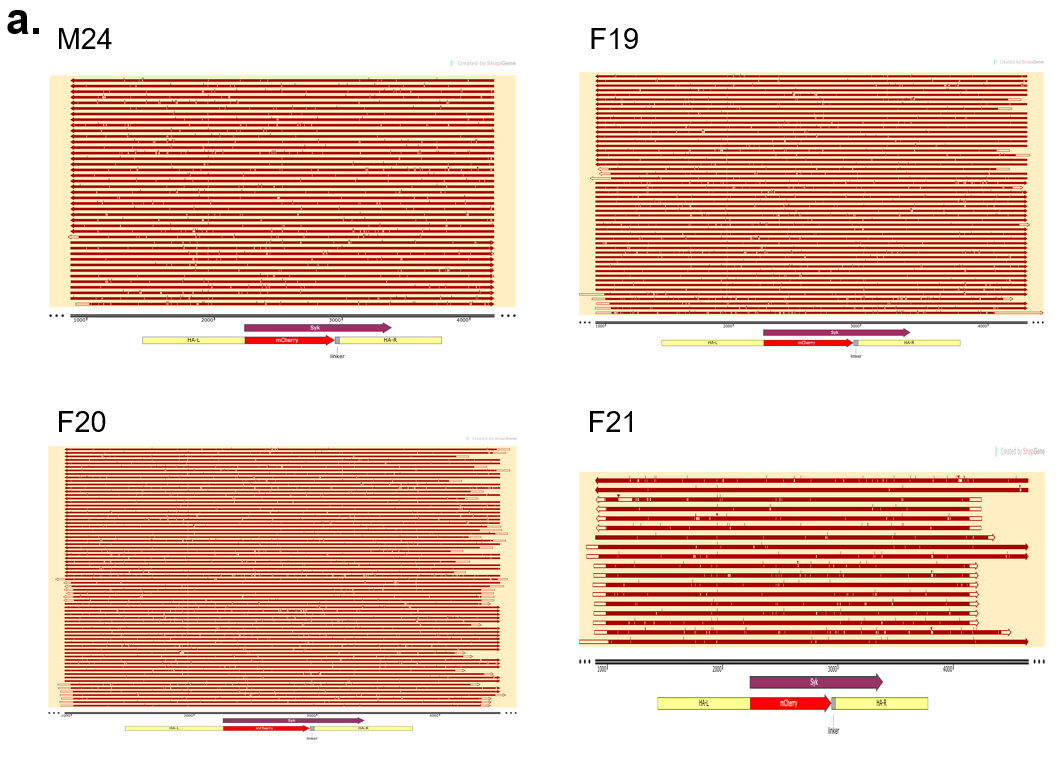

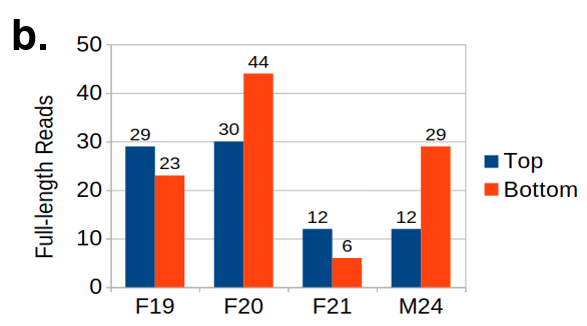

**Supplementary Figure 9.** Detailed analysis of LOCK-seq reads on the Syk-mCherry samples. **a.** Snapgene alignments of contiguous reads spanning the 5’ genomic region upstream of HA-L to 3’ genomic region downstream of HA-R for M24, F19, F20, and F21 that map entire KI cassette (homology arms in yellow, mCherry in red) and flanking genomic sequence. **b.** Barplot of full-length contiguous reads mapping to the top (blue) or bottom (red) strands of the donor (and homology arms) and flanking genomic DNA.

**Figure S1**
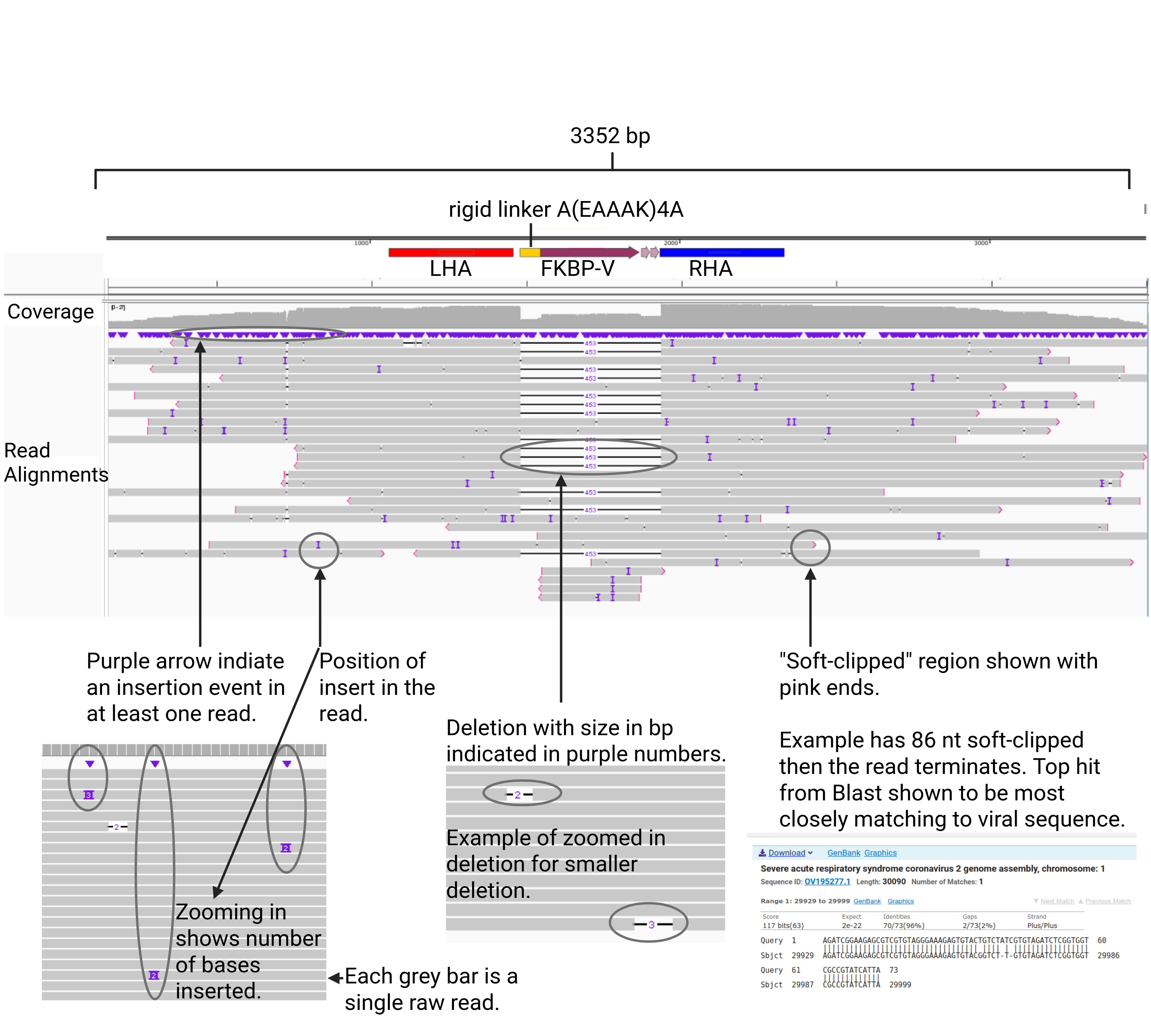
**0**

**Supplementary Figure 10.** IGV screenshot of raw reads aligned to Project MS2757 (**Table S1**) donor sequence. Purple arrows along top horizonal region below coverage plot indicate “insertion” events, or more likely sequence error. Each grey bar is a single read with purple vertical lines indicating insertion or error within the read. Deletions are shown as horizonal black bars with numbers indicating the total number of missing bases. Non-sequencing error is supported by multiple reads, as shown for the FKBP-V region, where the F1 sequence is heterozygous with roughly half of the reads containing the FKBP-V sequence and the other half not. Pink outline at the end of the reads indicates the occurrence of soft clipping where additional sequence beyond what is shown fails to align to the reference provided. In most cases, there is adapter sequence beyond that fails to align. The last three reads are described in more detail in **Fig. S11** below.

**Figure S11**

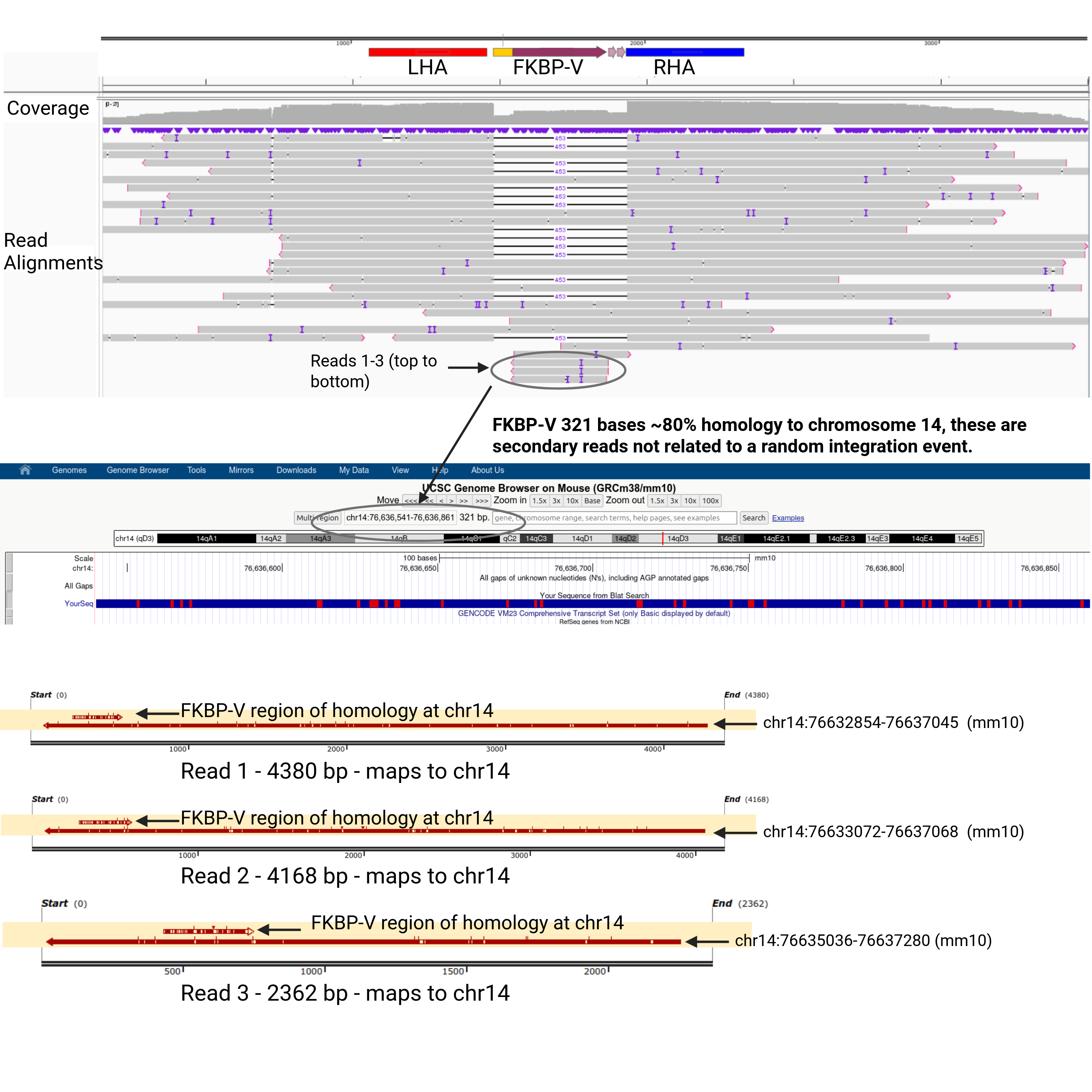

**Supplementary Figure 11.** IGV screenshot of raw reads aligned to Project MS2757 (**Table S1**) donor sequence. Pile-up of three reads at the bottom with shared soft-clipped regions is due to the partial identity (88.5%) to chromosome 14 (mm10, chr14:76636541-76636861, 321 bases). These reads have primary alignments to chr14 with the soft-clipped full-length reads mapping to the same region for all three.

**Figure S12**

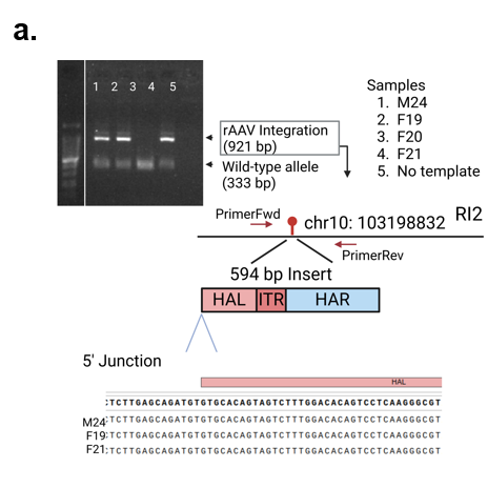

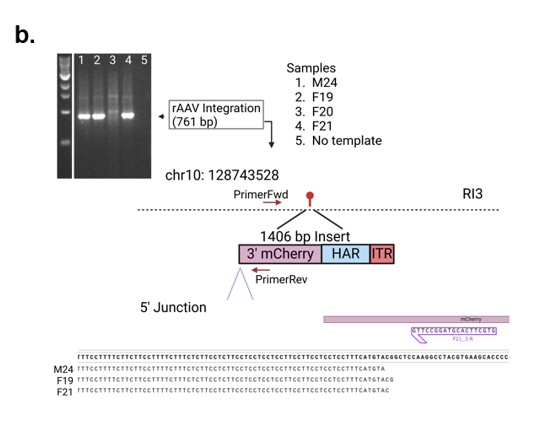

**Supplementary Figure 12.** Validation PCRs of rAAV random integration events in mSyk mouse models **a.** Gel image and sequencing data for random integration event upstream of Slc6a15 and **b.** within the first intron of Cd63.

**Figure S13**

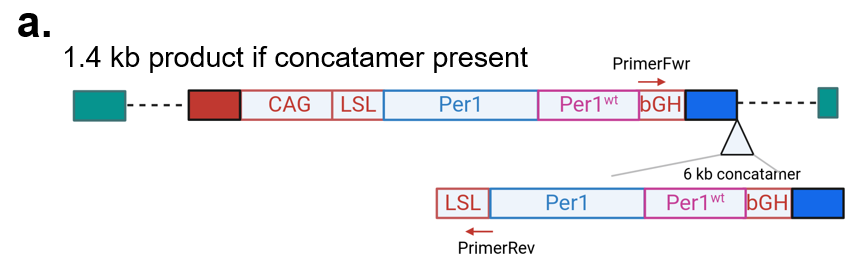

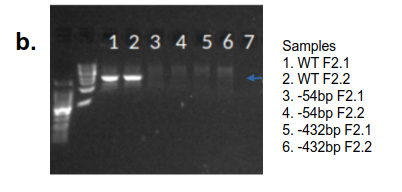

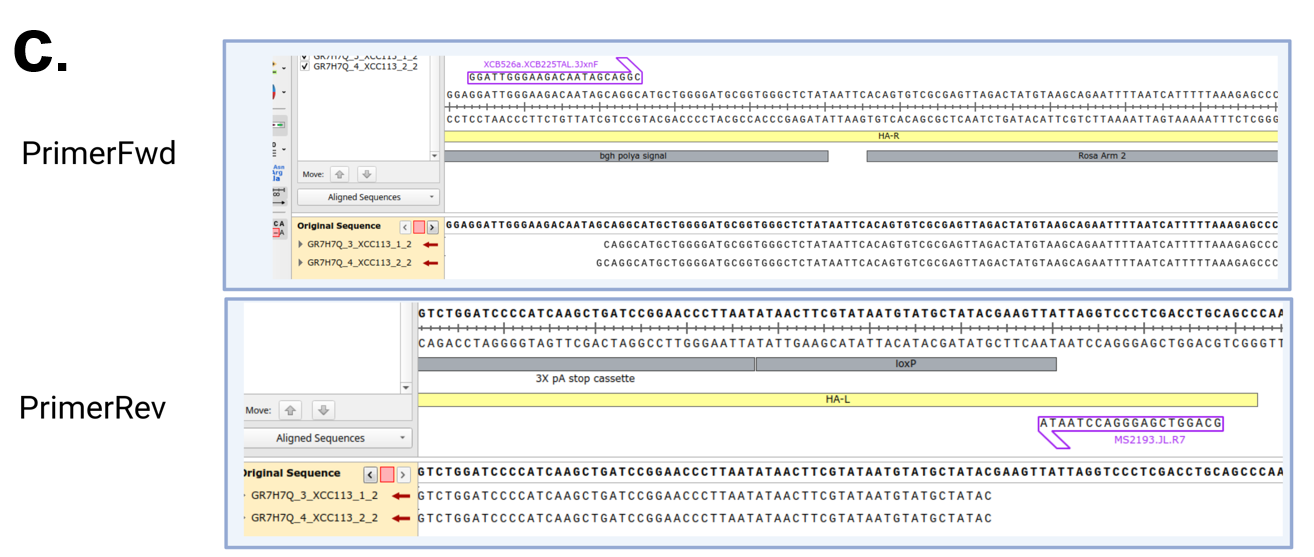

**Supplementary Figure 13.** Confirmation of donor concatenation detected by LOCK-seq. **a.** Schematic of concatenation event and primer binding sites (red arrows) for PCR validation. **b.** Gel image for validation PCRs showing positive 1.4 kb product for two F2 samples with WT Per1 cassettes. **c.** Alignments of 5’ and 3’ sequencing results for WT F2.1 (top) and WT F2.2 (bottom).

**Figure S14**

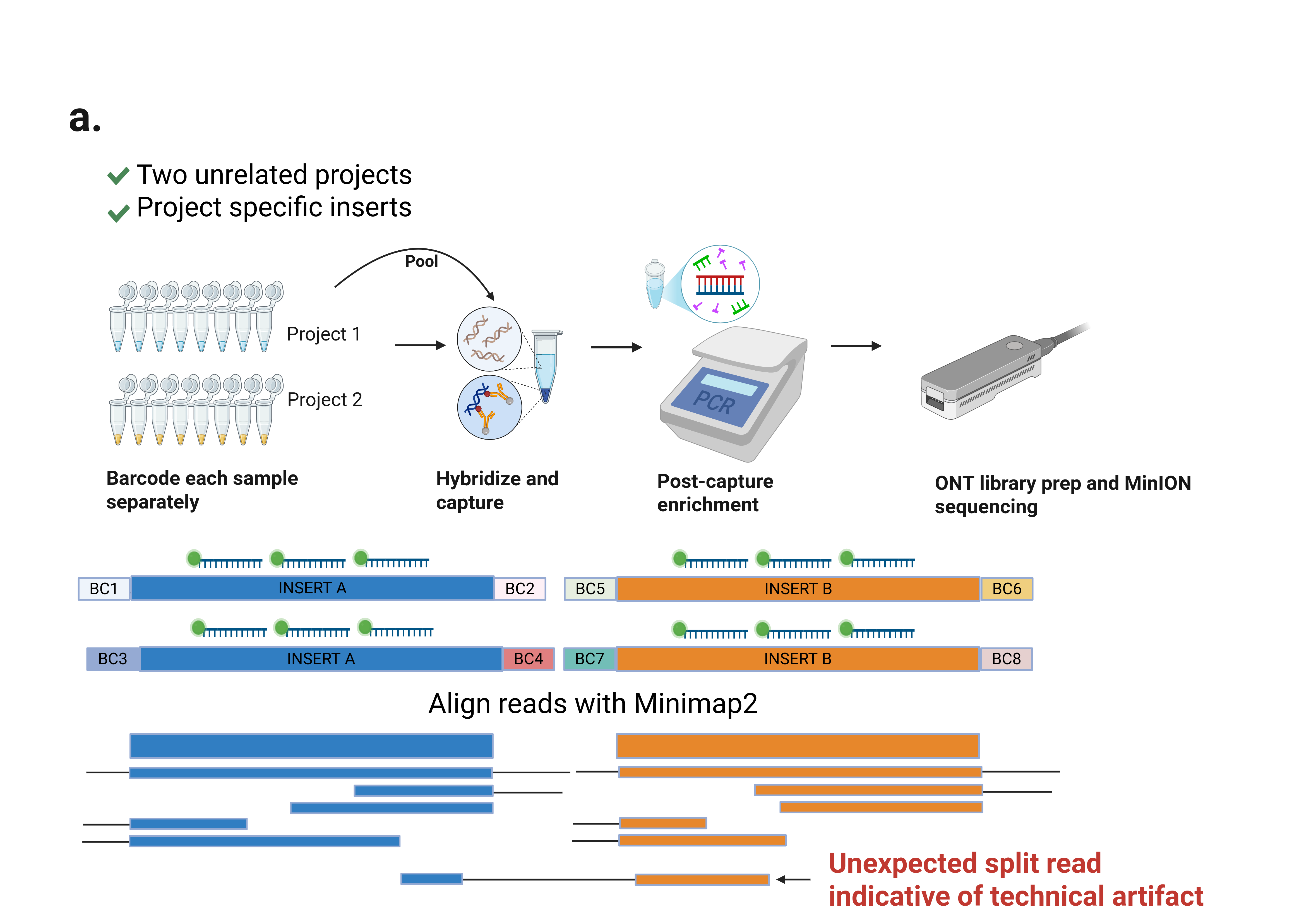

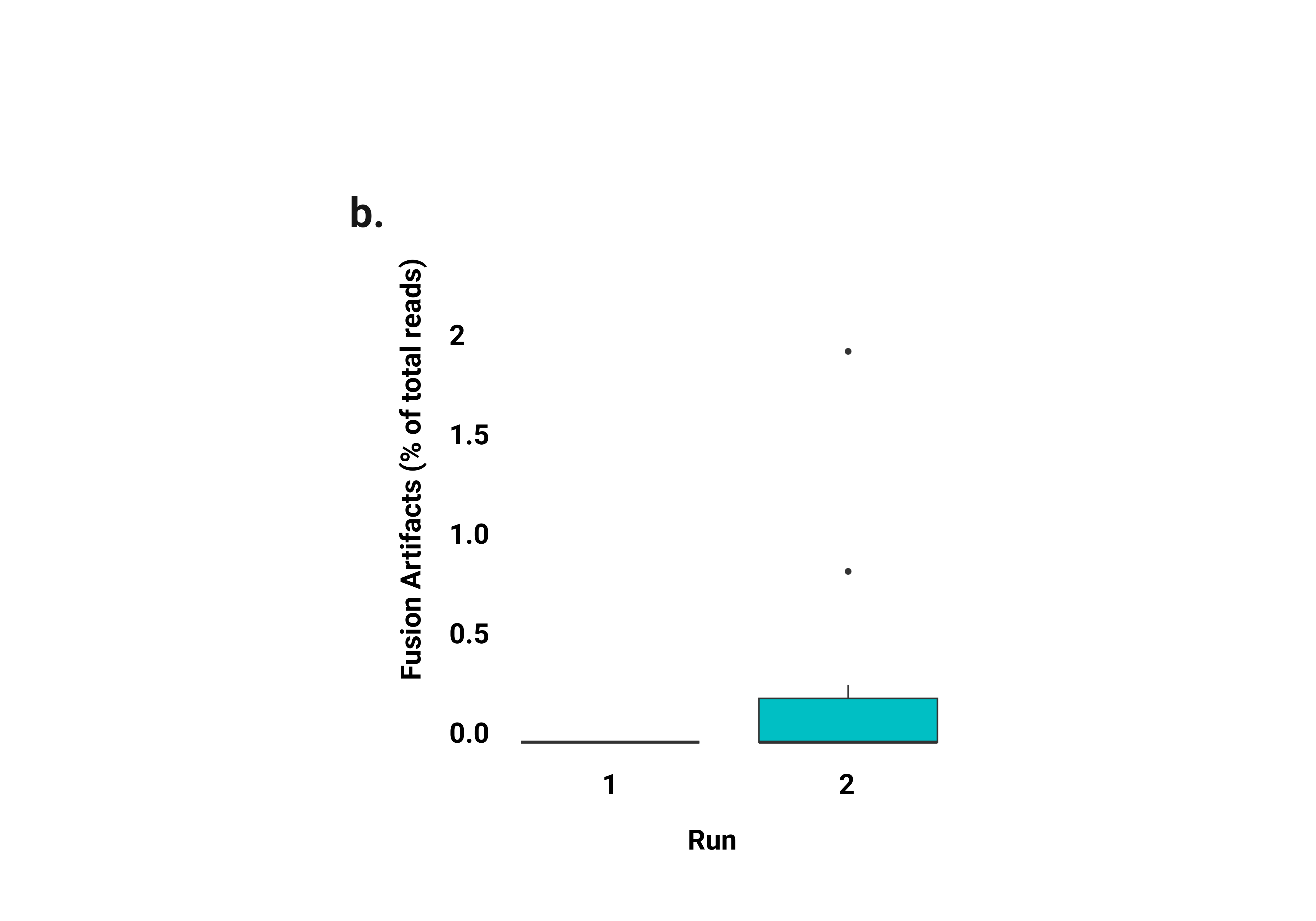

**Supplementary Figure S14** Analysis for artificial fusion reads in LOCK-seq **a.** Schematic of two unrelated KI projects with unique insert sequences (A vs. B). Following standard LOCK-seq protocol, barcoded samples for the two projects were pooled after the first round of PCR and hybridized with both sets of probes and carried through the remaining steps of LOCK-seq to result in a single library. MinION runs were analyzed to look for fusion reads where one read aligned to both unique insertion events from the two unrelated projects. **b.** Two independent runs are shown here, each with different projects and samples (projects XCF18a and JK413 for run #1, and projects MS2205_MS2206 and XCD68a for run #2, as listed in **Table S8**). In both cases, the total fusion reads across samples were <2% of mapped reads.

**Figure S15**
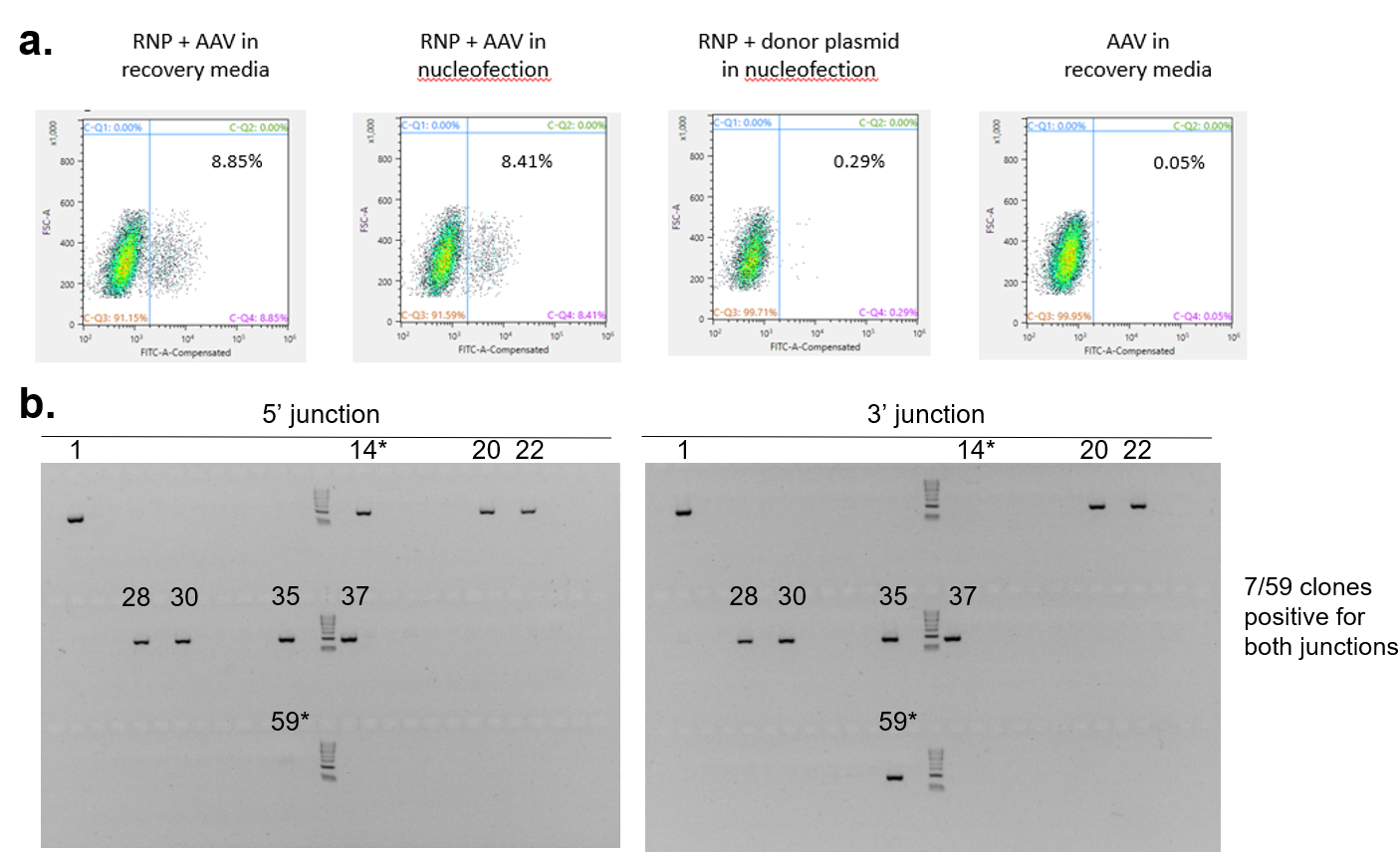

**Supplementary Figure 15.** Efficient KI in iPSCs using rAAV donors with CRISPR/Cas9. **a**. Flow cytometry of iPSCs edited with RNPs and a cDNA-GFP donor in rAAV vs plasmid format. **b.** Gel images of 5’ and 3’ junctions for 59 single-cell derived iPSC clones from a pool edited with cDNA-GFP donor in rAAV format. Numbers on top of lanes indicate clone IDs positive for 5’ and 3’ junctions, and asterisks indicate those clones only positive for a single junction. Gel image is also shown in **Fig.S19** to compare to alternative screening method.

**Figure S16**
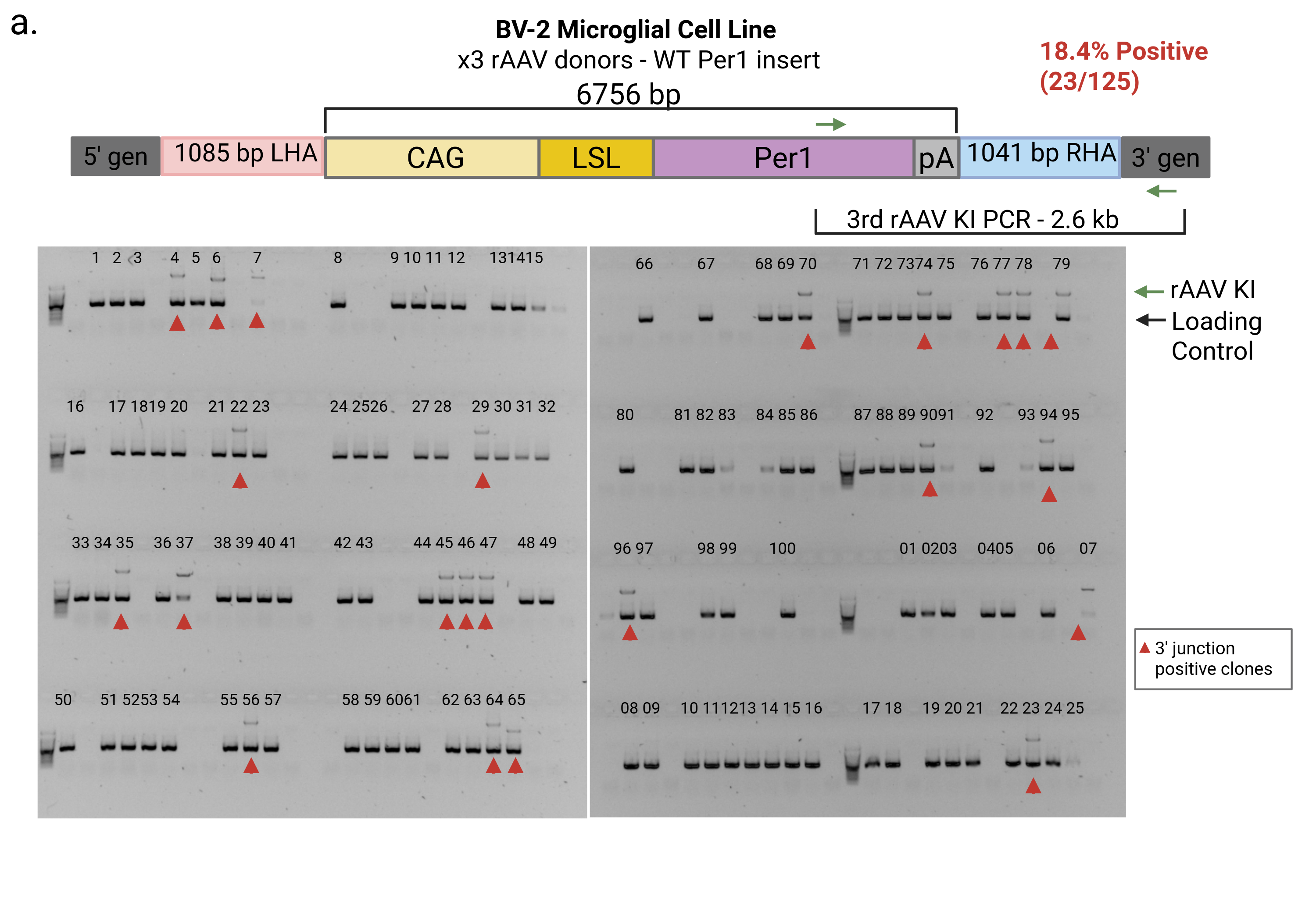

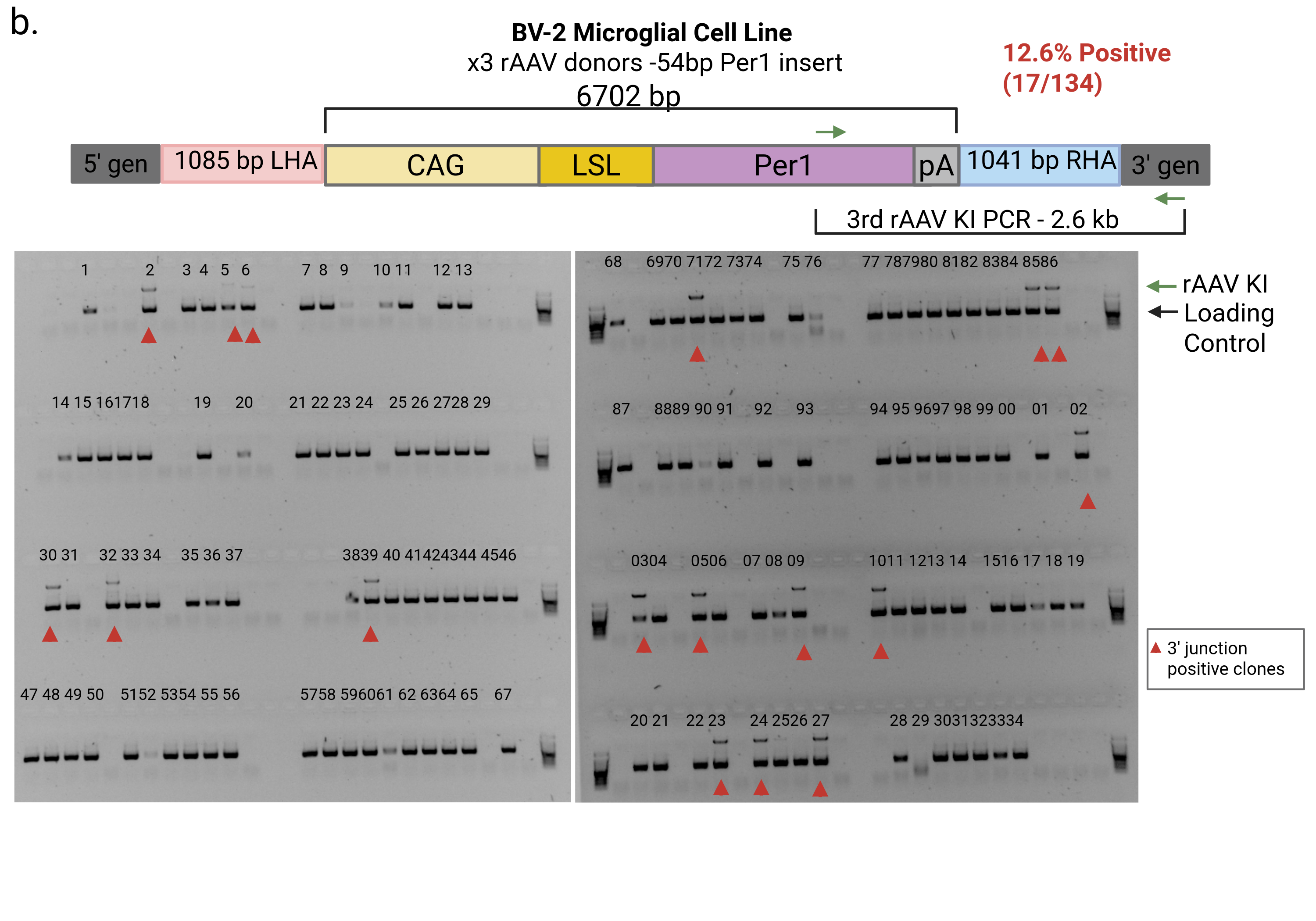

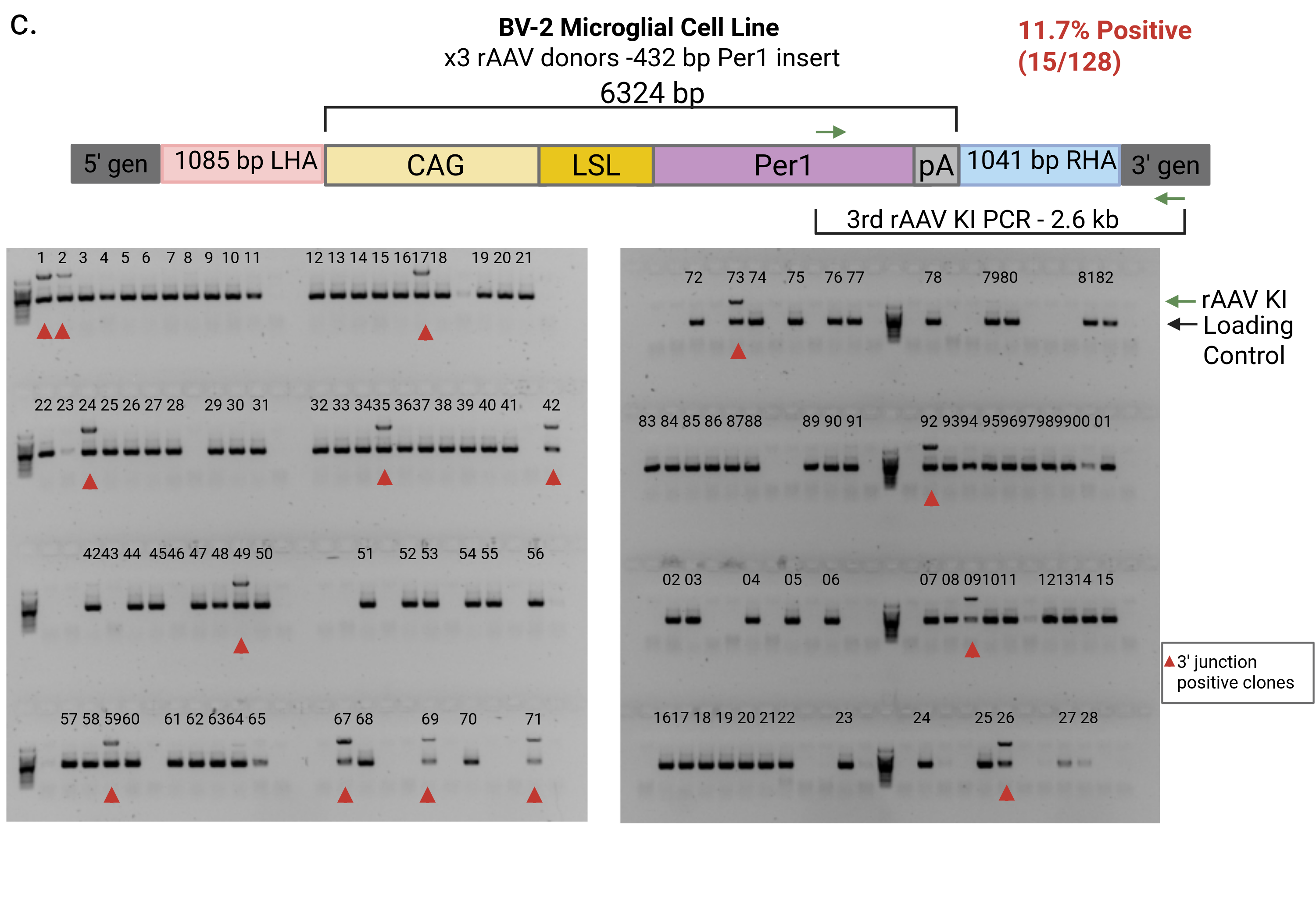

**Supplementary Figure 16.** rAAV donors enabled knockin in BV2 cells that are highly sensitive to exposure to exogenous DNA in ssODN and plasmid forms (see **Fig.6a**, where low HDR by ssODN is shown). Gel images of high efficiency and consistence of positive junction PCR amplifications from single-cell BV2 clones transduced with x3 rAAV donors for sequential knock-in of **a**. WT Per1  **b.** Per1 mutant with a 54 bp deletion or **c.** 432 bp deletion. All gels show multiplexed PCR for 3’ junction PCRs (larger 2.6 kb band) between donor 3 and 3’ genomic DNA as well as amplification of a 0.4 kb region of Nsmce3 as a loading control.

**Figure S17**

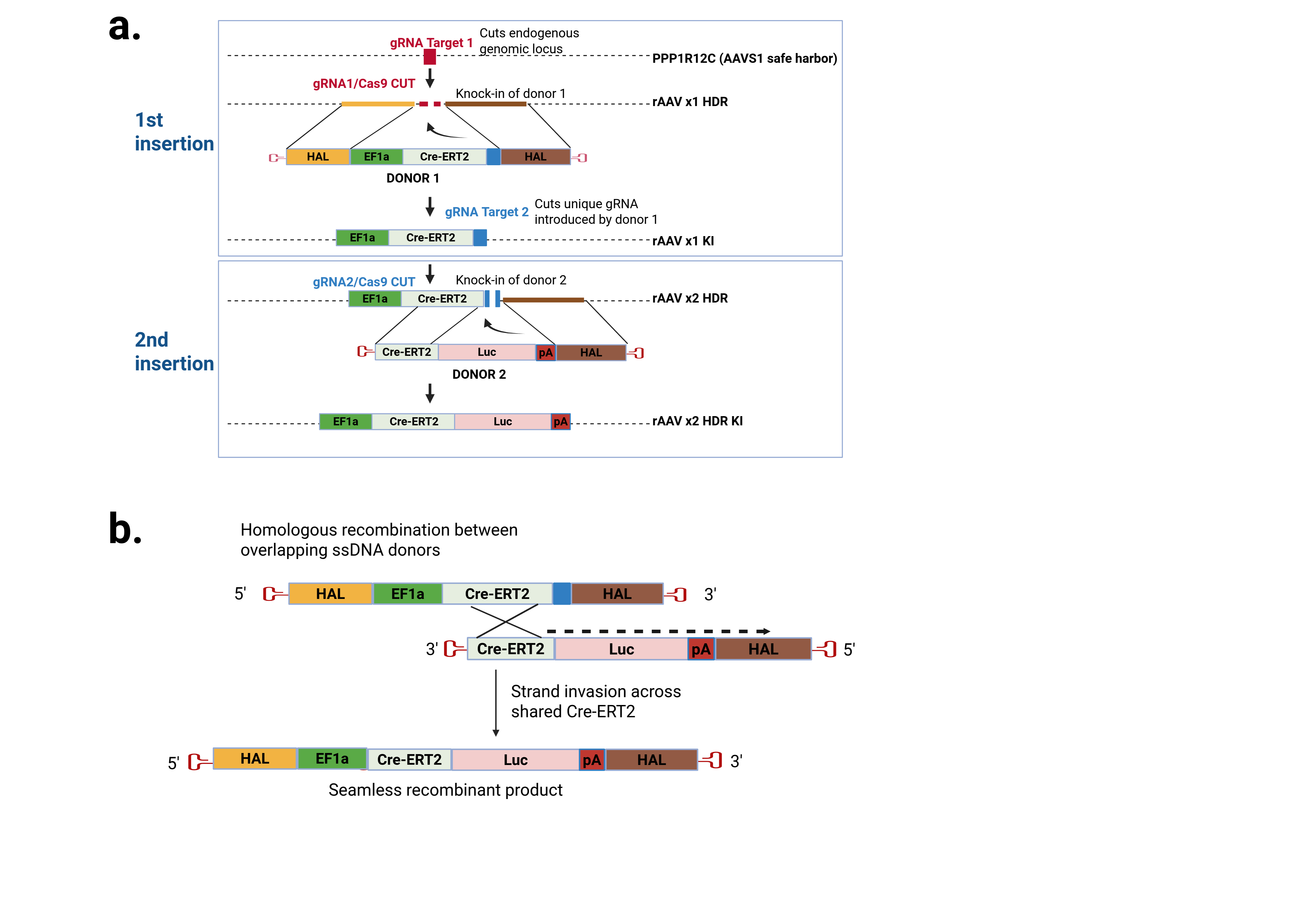

**Supplementary Figure 17.** The schematic of 2-rAAV KI single or dual gRNAs. **a.** Both rAAV donors and two RNPs are co-delivered to embryos or cells. The first gRNA cleaves the genomic target site to mediate HDR from the first donor (1^st^ insertion), and successful knock-in of the first insert introduces in the second, unique gRNA target site into the allele. The second gRNA target site is not cleavable in the single stranded rAAV genome. The second RNP then cleaves the newly introduced gRNA site, allowing the second insertion to occur via HDR (2^nd^ insertion). **b.** In the absence of the second gRNA, a seamless recombinant rAAV donor could be generated by recombination of ssDNA rAAV donors at regions of shared homology.

**Figure S18**

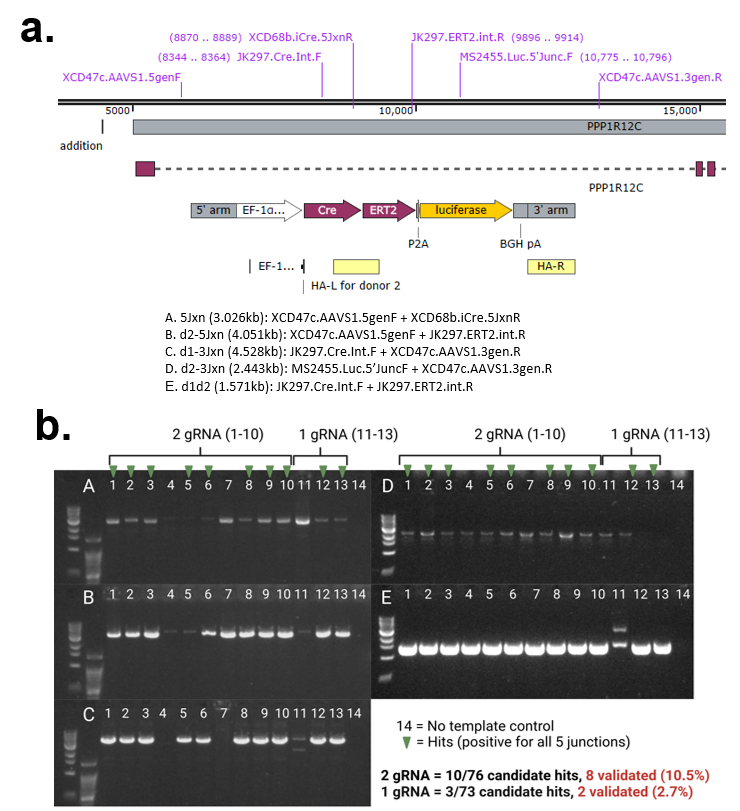

**Figure S18**
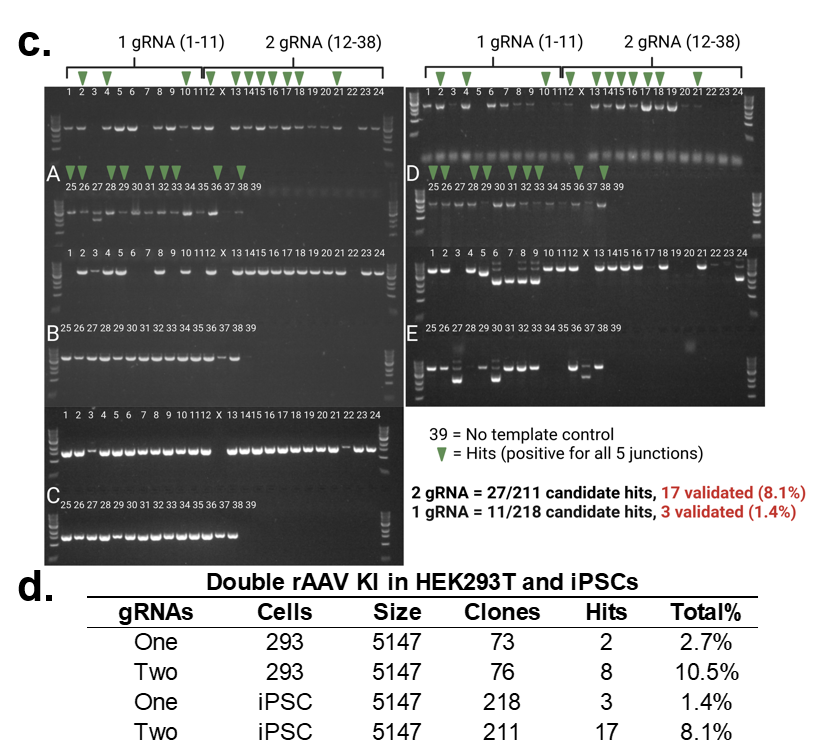

**Supplementary Figure 18.** Dual rAAV KI of XCD47c using 1 vs. 2 gRNAs in HEK293s and iPSC cells. **a**. Schematic of primers used for five junction PCRs listed A-E. Homology arms (yellow bars) of second rAAV donor are annotated. **b.** Validation PCRs of candidate hits 1-13, no template control is 14, from HEK293Ts edited using 2 gRNAs (8 positives out of 76 screened) vs. 1 gRNA (2 positive out of 73 screened). **c.** Validation PCRs of candidate hits 1-38, no template control is 39, from iPSCs edited using 2 gRNAs (17positives out of 211 screened) vs. 1 gRNA (3 positives out of 218 screened). Even though without cleavage by the gRNA brought in by donor 1, the second donor was able to insert at a lower level, implying the possibility of recombination between donors 1 and 2 prior to HDR.

**Figure S19**
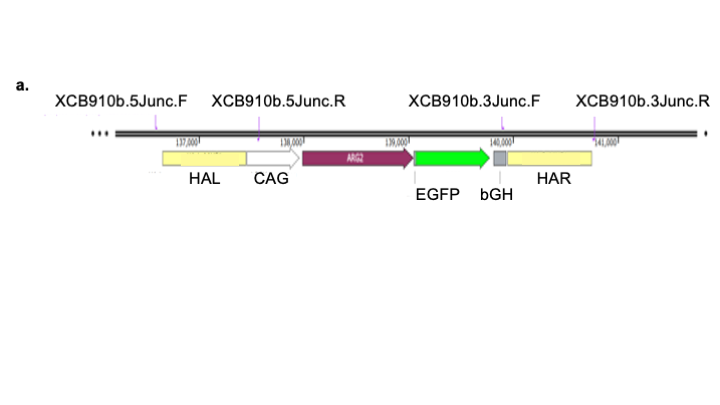

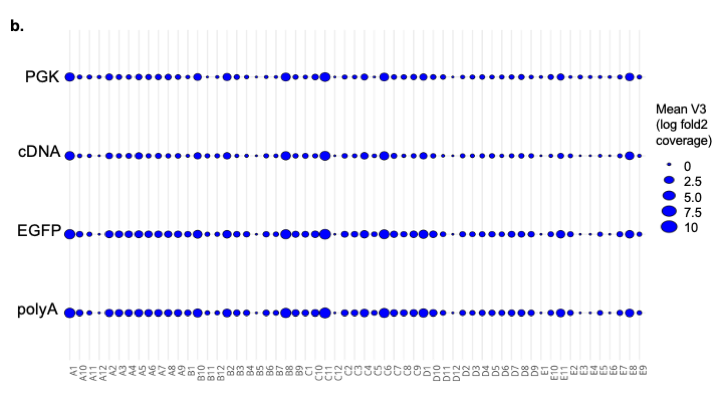

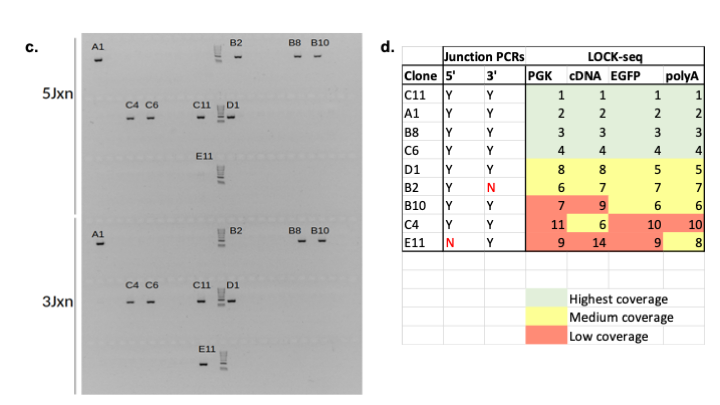

**Supplementary Figure S19** Ranking of 59 iPSC clone screening using LOCK-seq. **a**. Schematic of a single rAAV donor-mediated knock-in in iPSCs with primer binding sites shown used for 5’ and 3’ junction PCRs. **b**. Circle plot of mean log2 coverage across 59 clones screened by LOCK-seq and **c**. gel image of 5 ‘and 3’ junction PCRs (same as Fig.S4). **d**. Ranking of clones based on highest to lowest mean log2 coverage showing that LOCK-seq.

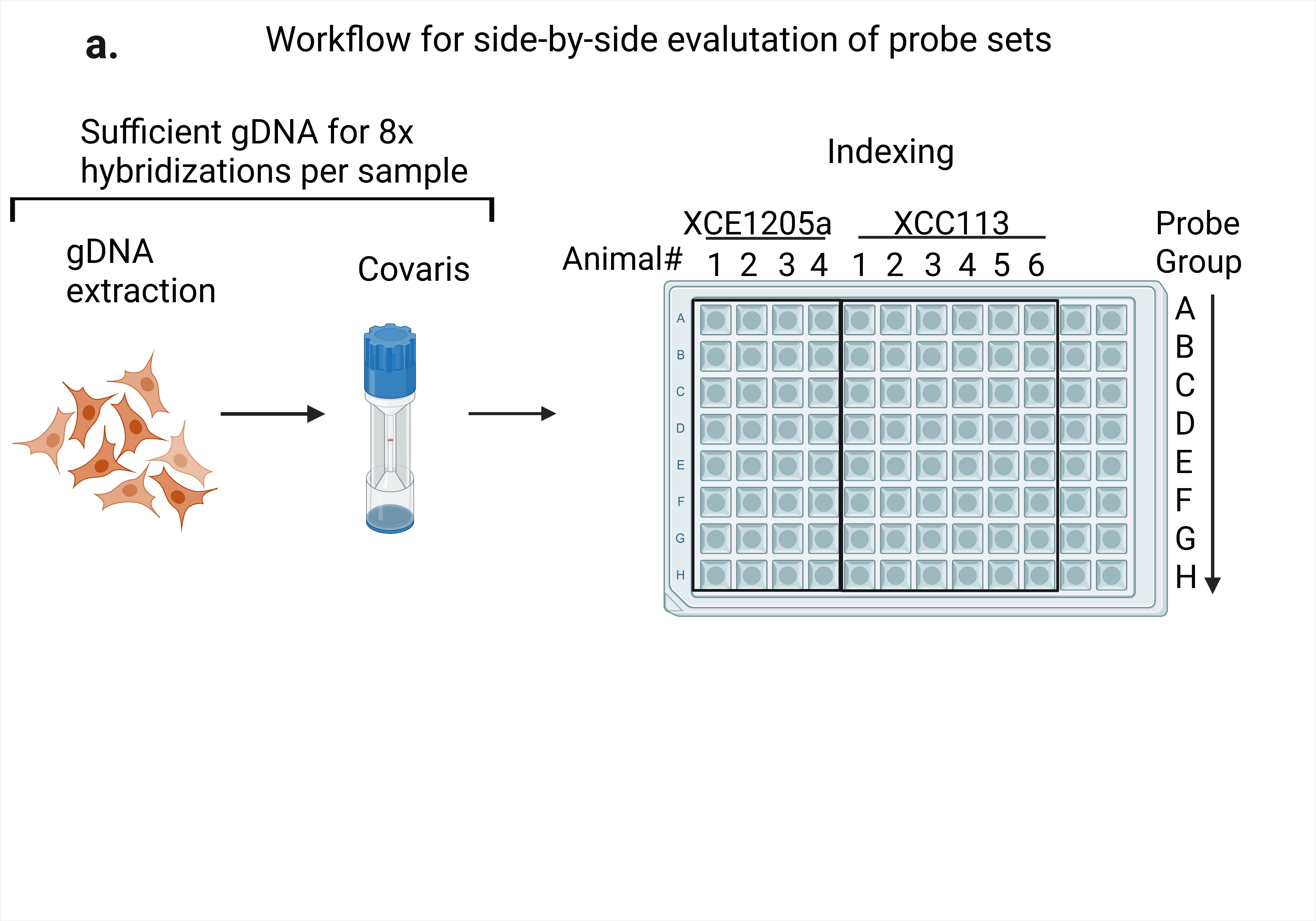

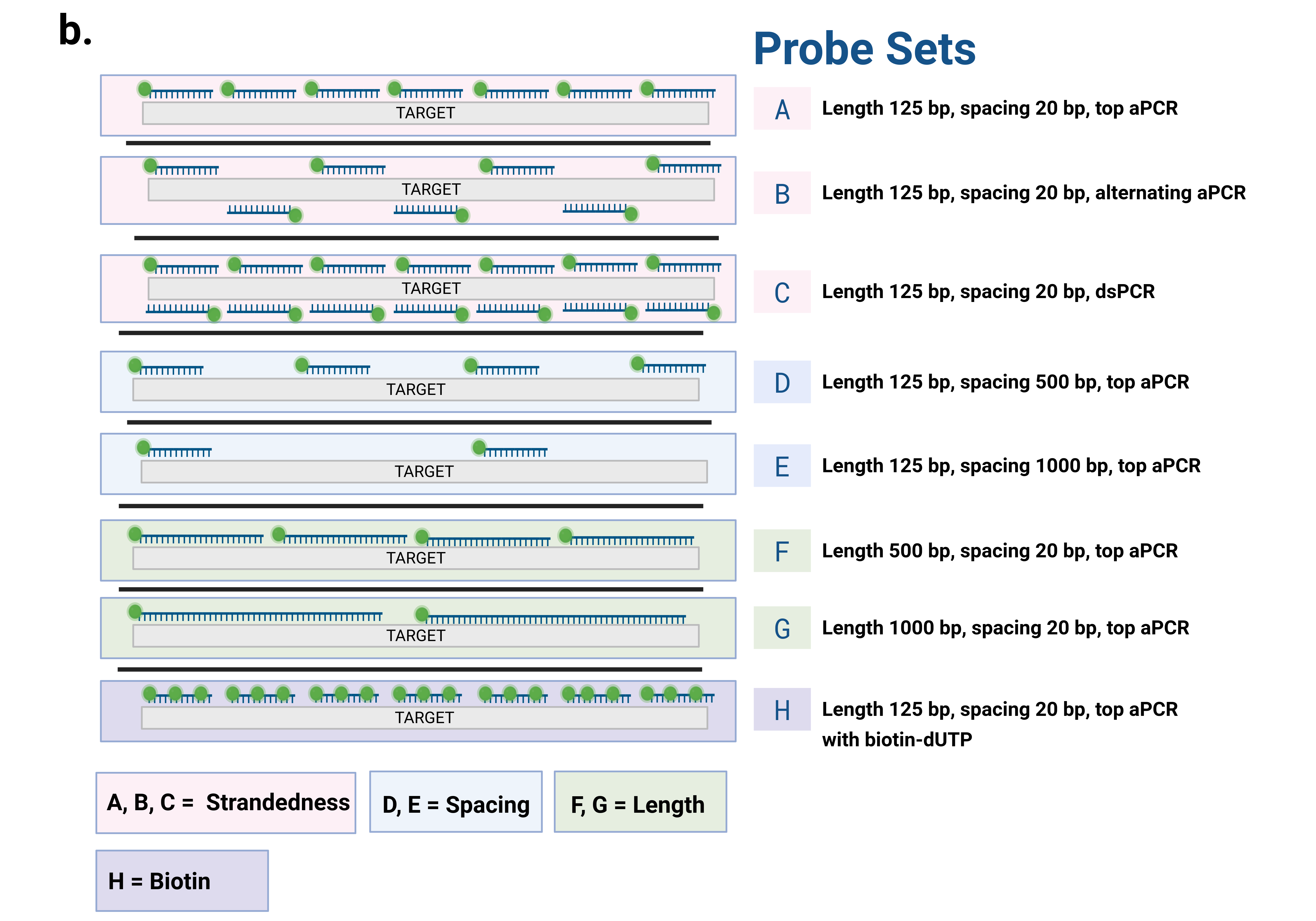

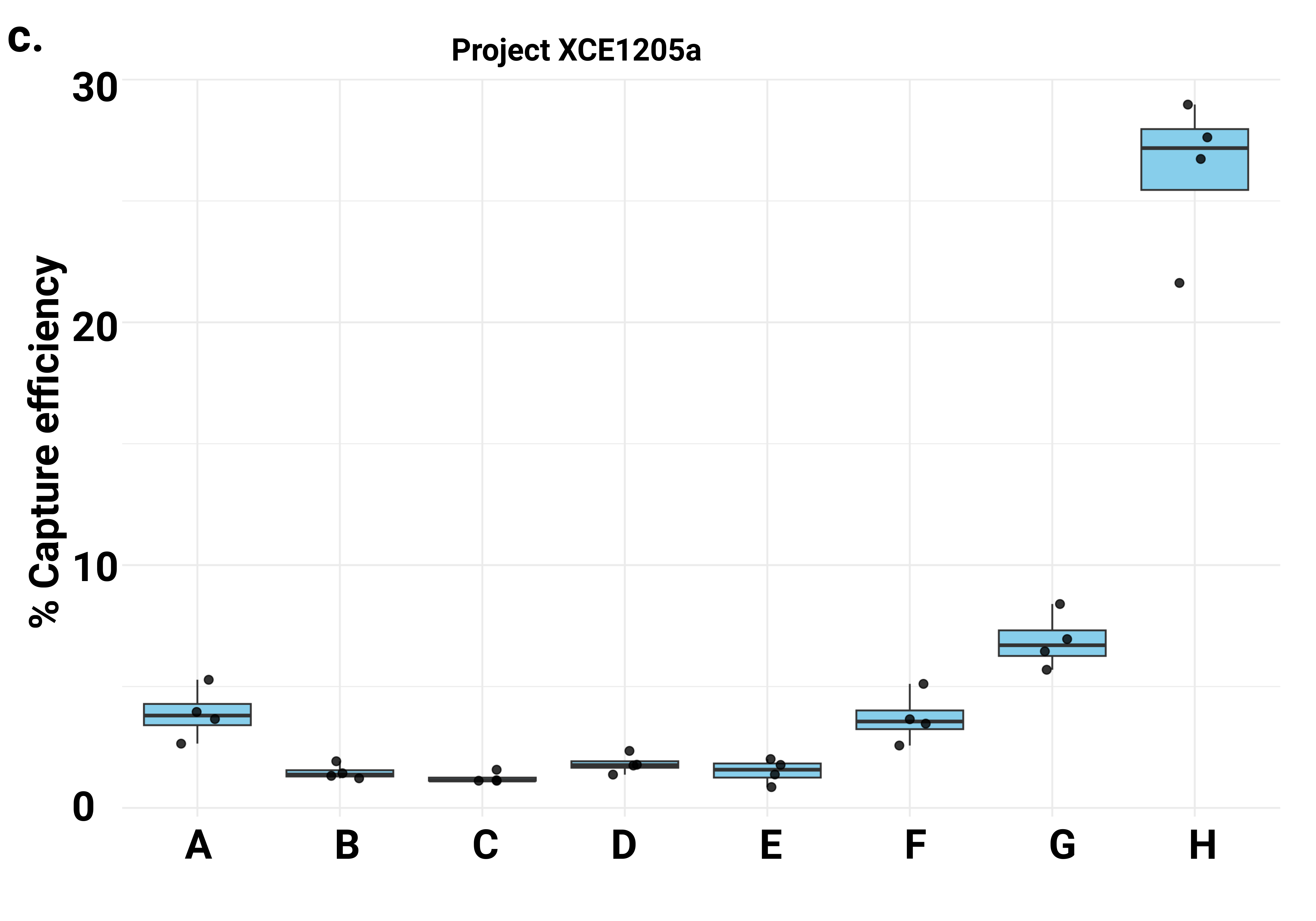

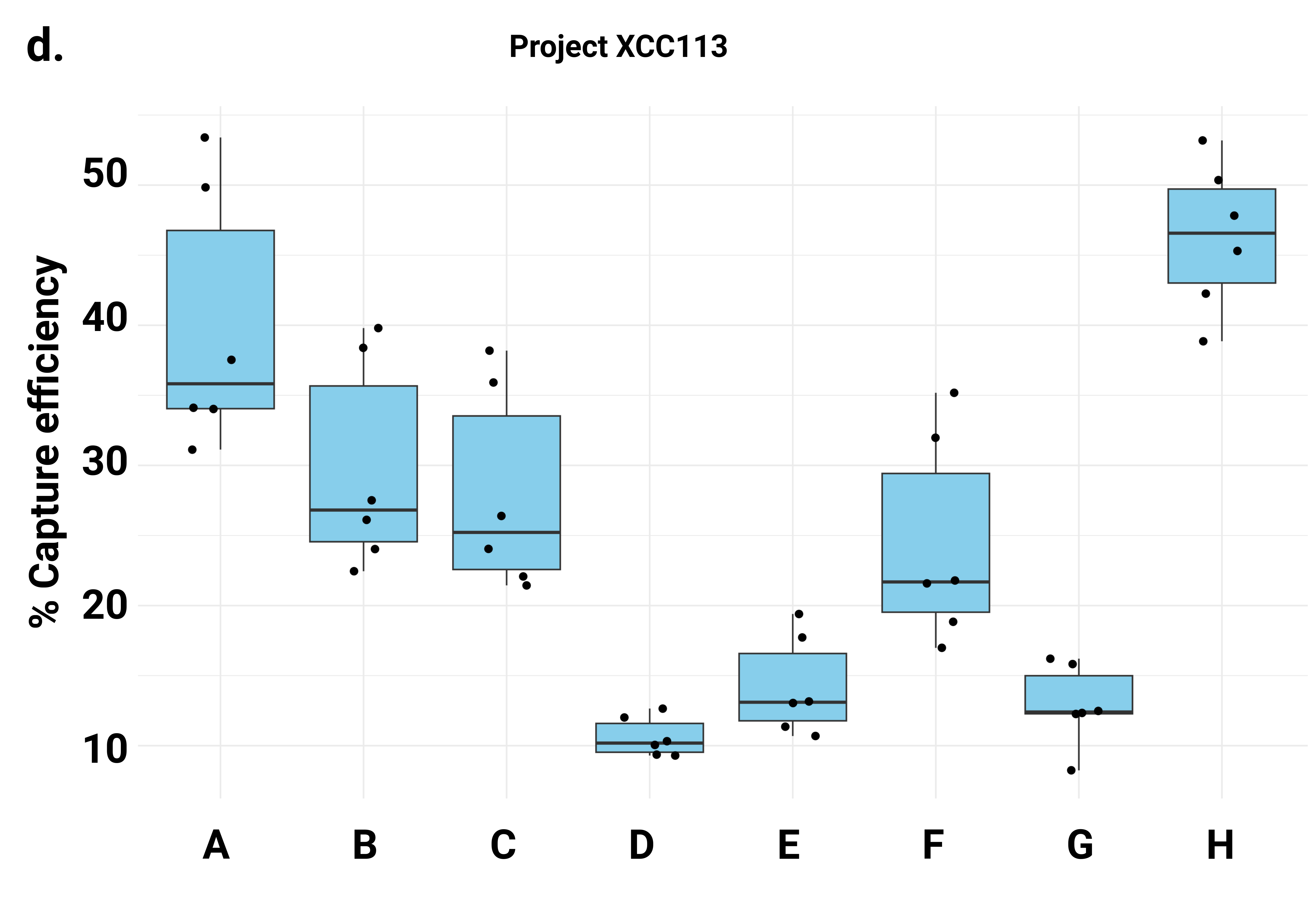

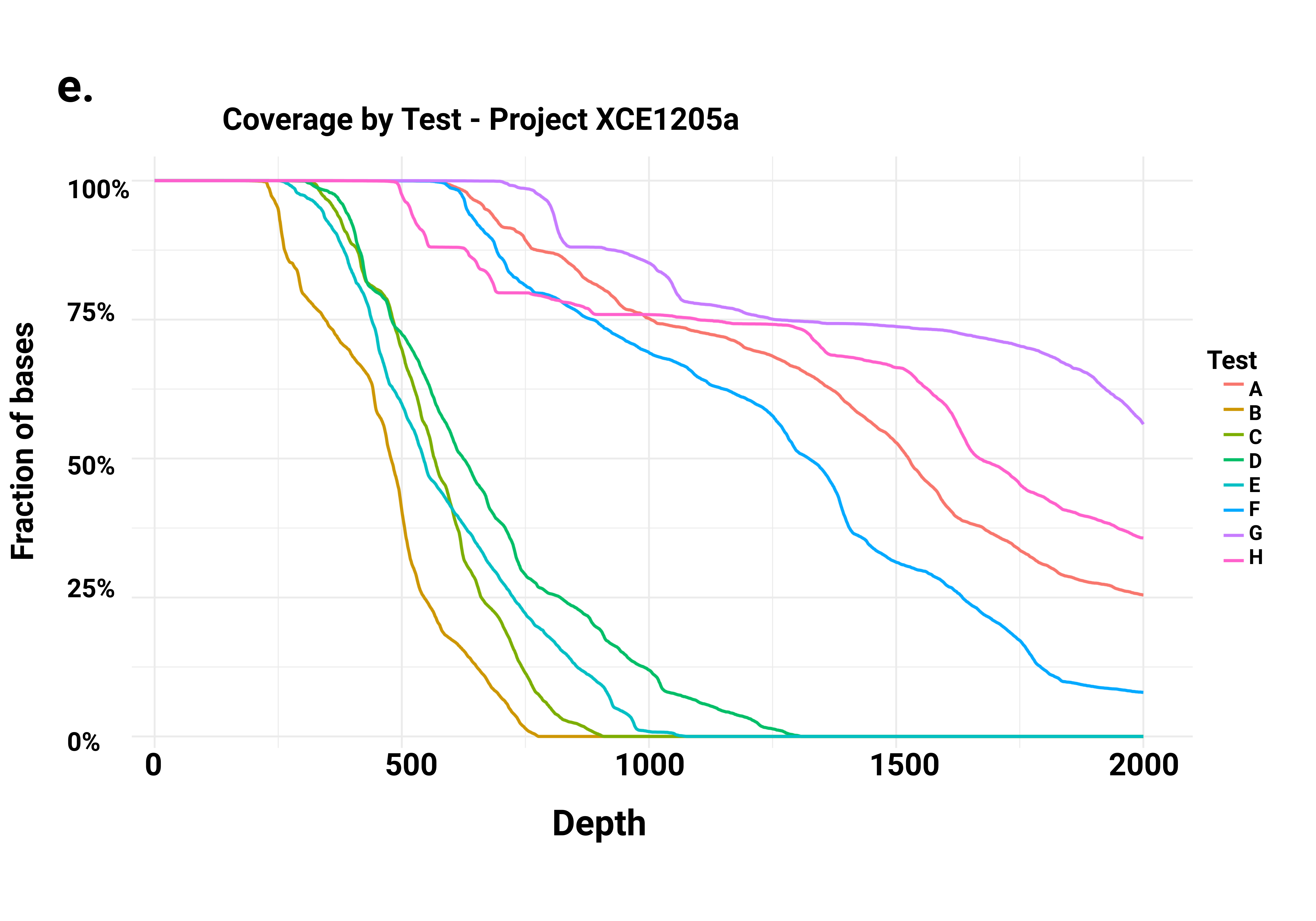

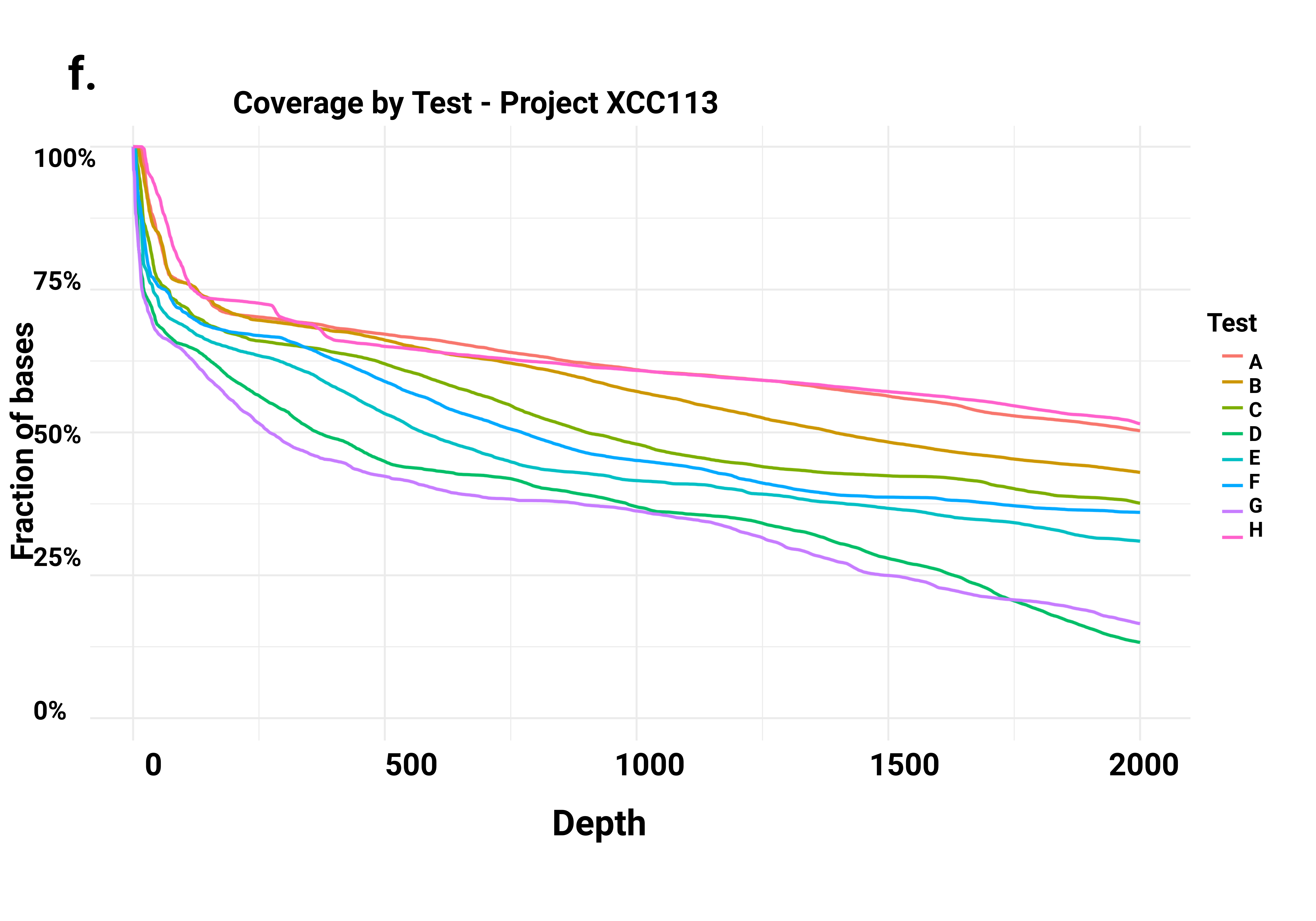

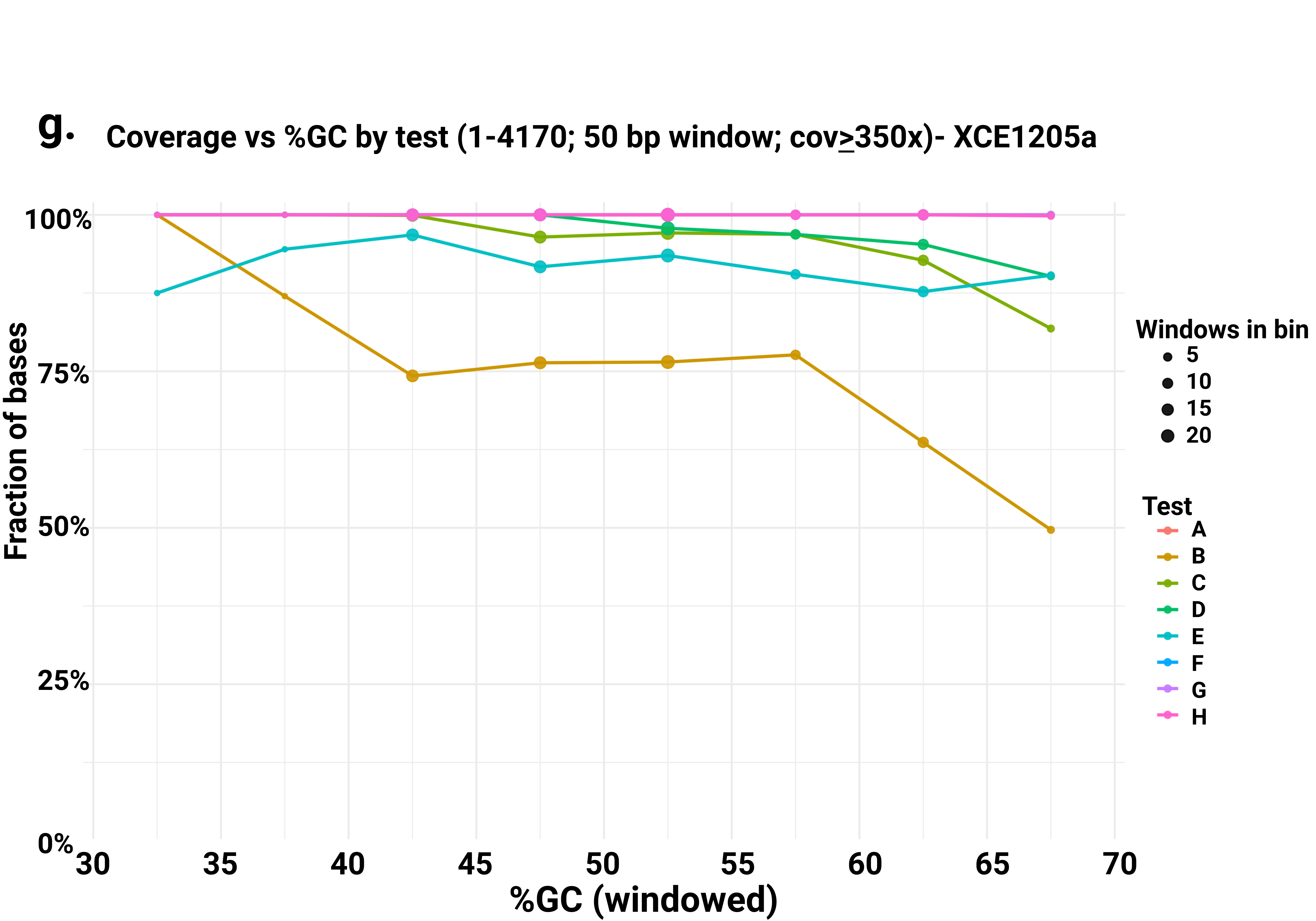

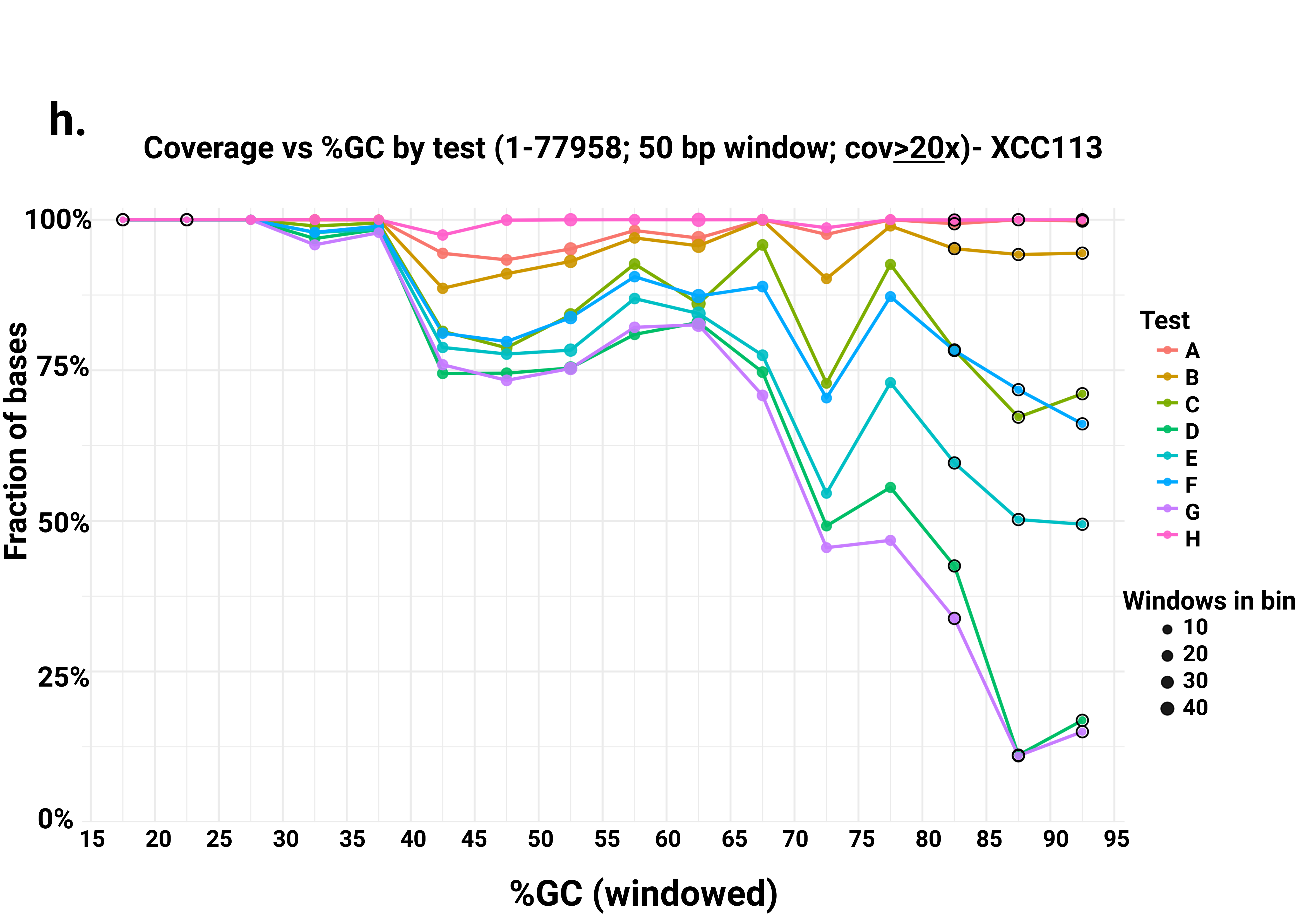

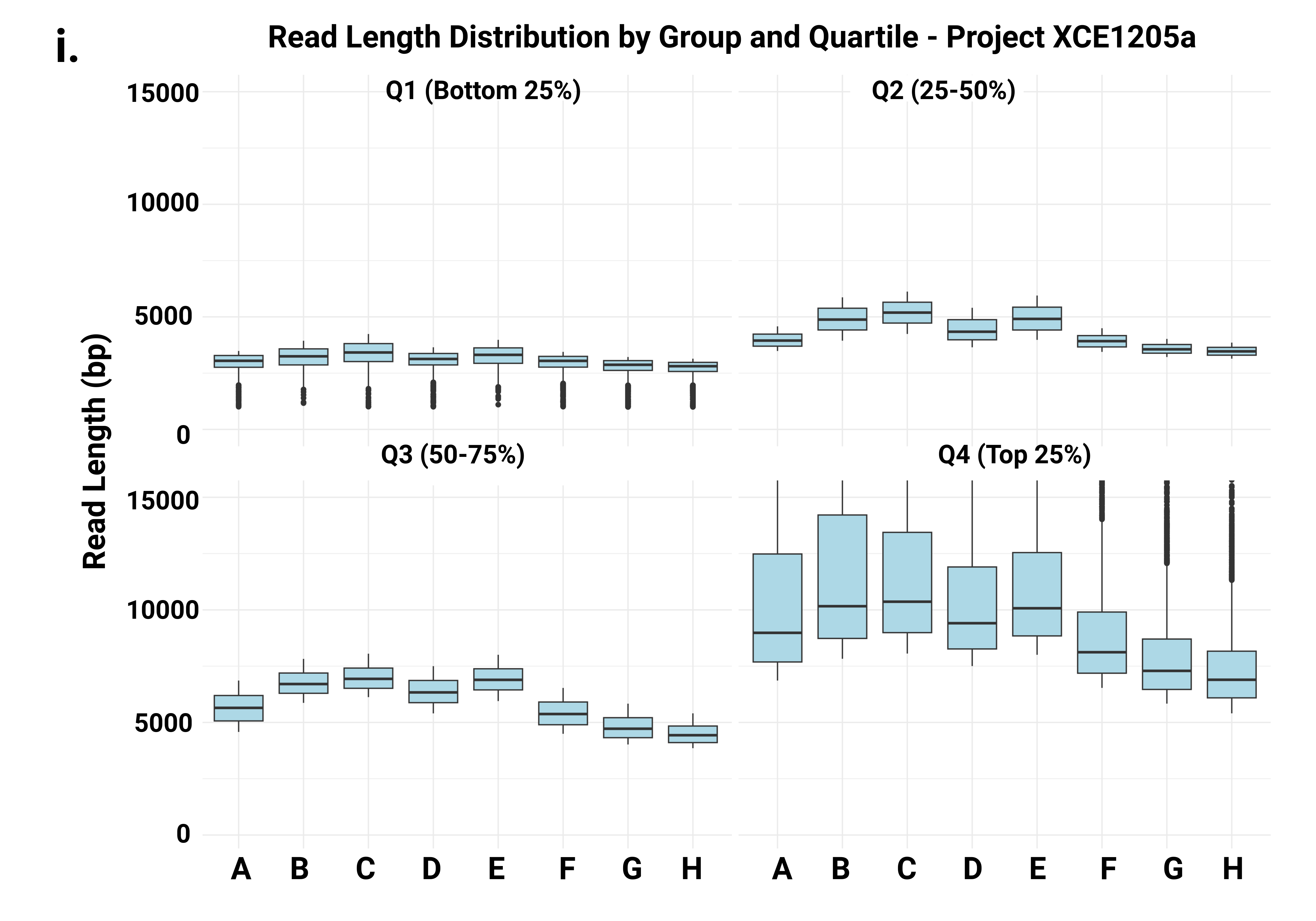

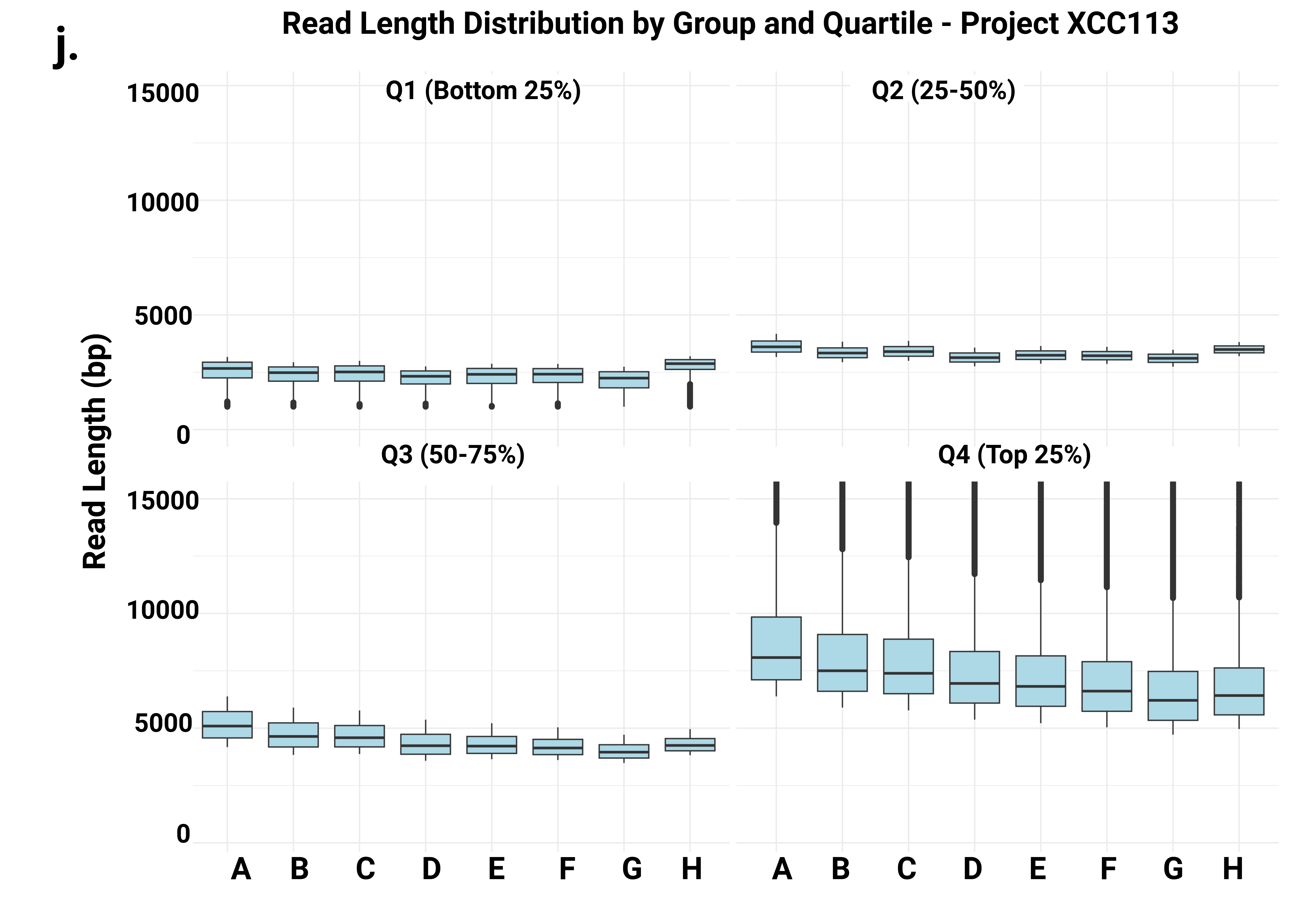

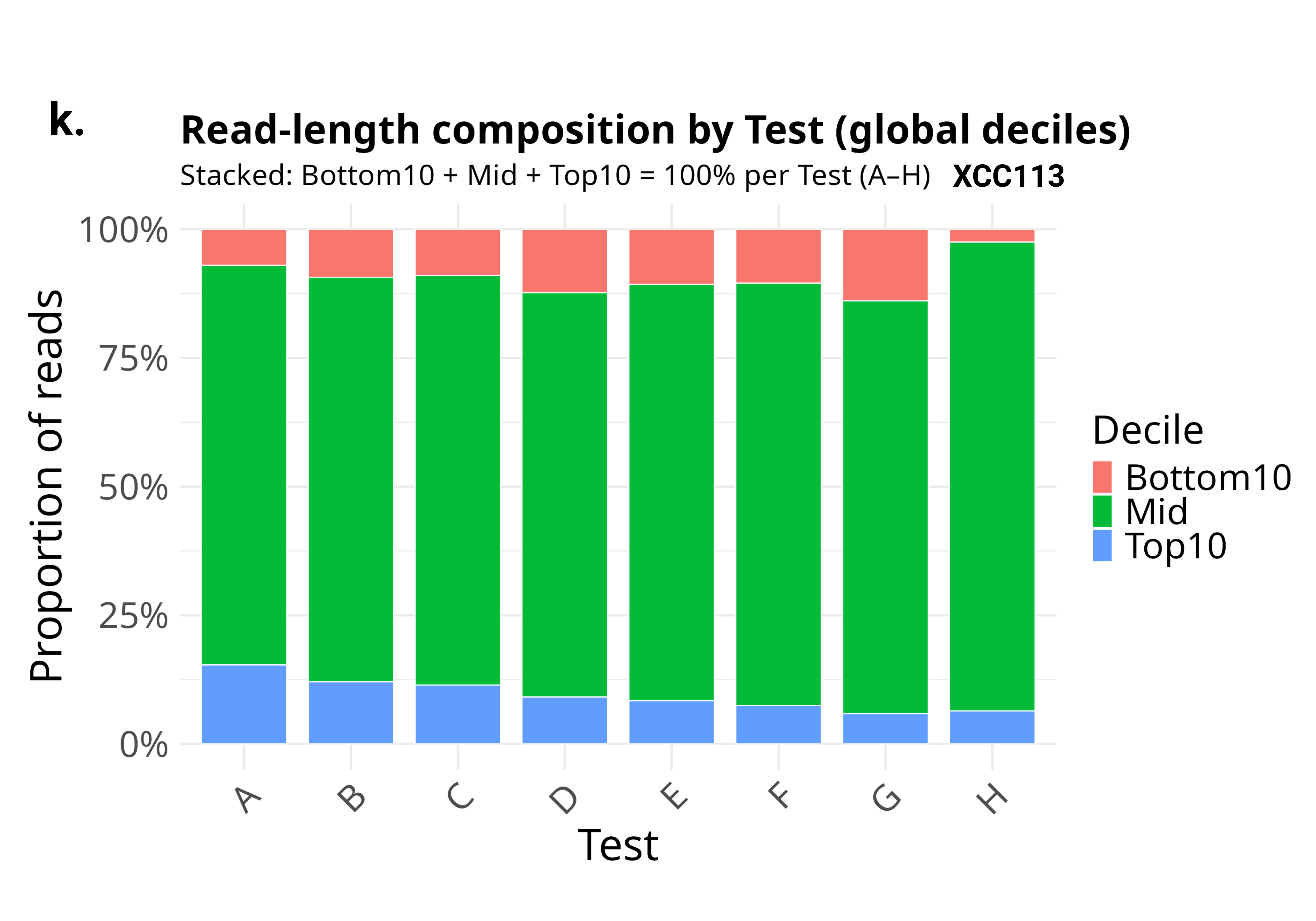

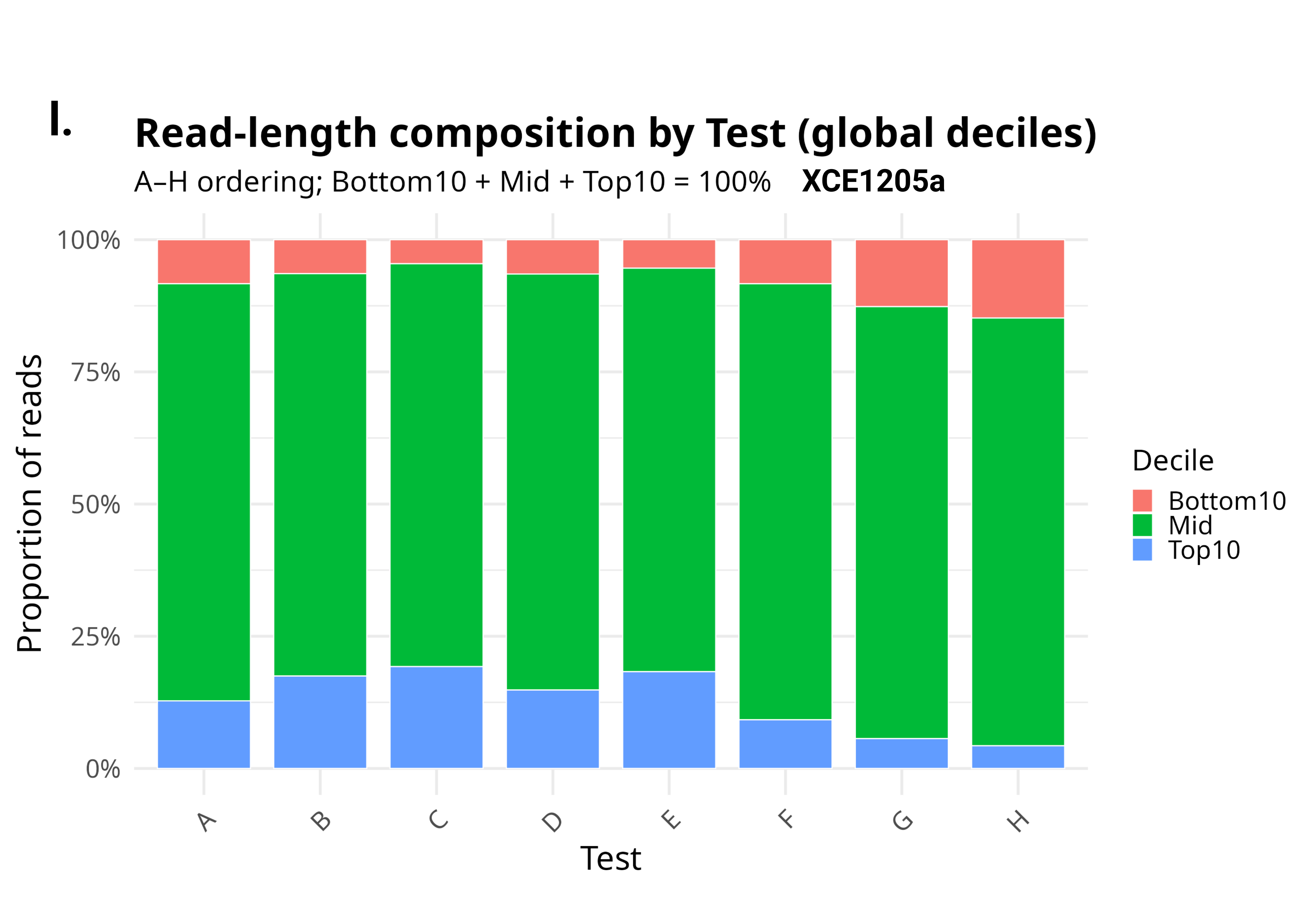

**Supplementary Figure S20.** Probe design optimization. **a.** Workflow for evaluation of eight different probe designs (A-H) in parallel, using samples from two projects: XCE1205a (n=4 F0s) and XCC113 (n=6 F1s). **b.** Various probe designs tested: the same probe length (125 bases) and density (20 bp spacing) against one strand (A), alternating strands (B) and both strands (C), lower density (D-E), longer probe length (F and G), or additional biotin moieties (H). Probe sets of 125 bases in length designed against the top strand with 20 bp spacing and additional biotin moieties (H) had the highest capture efficiency (total mapped reads/total reads, y-axis), followed by 5’-bitoin modified probes (A), across projects XCE1205a (**c**) and XCC113 (**d**). The coverage depth (fraction of total bases on the y-axis versus depth of coverage on x-axis) across XCE1205a (**e)** and XCC113 (**f**) was also highest with probe designs A and H. Coverage versus %GC content binned in 50 bp windows, fraction of bases covered (y-axis) are plotted across increasing %GC bins in XCE1205a (**g**) and XCC113 (**h**). Size of data circle scaled to size of bin (number of loci that fit that %GC). Consistent with reduced overall coverage, lower probe density (D-E) performed worst across GC-rich regions while probes with more biotin moieties achieved the most consistent coverage. **i-j.** Read length distribution by test and quartile. Read length (y-axis, bp) is shown per sample, with split distributions within samples across quartiles. **k-l.** Read length composition by test using global deciles (top 10% and bottom 10% for all samples for each project), plotting the proportion that constitutes the globally calculated deciles for each sample.
